# Cell-free prototyping of limonene biosynthesis using cell-free protein synthesis

**DOI:** 10.1101/2020.04.23.057737

**Authors:** Quentin M. Dudley, Ashty S. Karim, Connor J. Nash, Michael C. Jewett

**Affiliations:** Department of Chemical and Biological Engineering and Center for Synthetic Biology, Northwestern University, Evanston, IL 60208, USA; Earlham Institute, Norwich Research Park, Colney Lane, Norwich, NR4 7UZ, United Kingdom

**Keywords:** cell-free metabolic engineering, limonene, iPROBE, cell-free metabolic pathway prototyping, cell-free protein synthesis, synthetic biology

## Abstract

Metabolic engineering of microorganisms to produce sustainable chemicals has emerged as an important part of the global bioeconomy. Unfortunately, efforts to design and engineer microbial cell factories are challenging because design-built-test cycles, iterations of re-engineering organisms to test and optimize new sets of enzymes, are slow. To alleviate this challenge, we demonstrate a cell-free approach termed *in vitro* Prototyping and Rapid Optimization of Biosynthetic Enzymes (or iPROBE). In iPROBE, a large number of pathway combinations can be rapidly built and optimized. The key idea is to use cell-free protein synthesis (CFPS) to manufacture pathway enzymes in separate reactions that are then mixed to modularly assemble multiple, distinct biosynthetic pathways. As a model, we apply our approach to the 9-step heterologous enzyme pathway to limonene in extracts from *Escherichia coli*. In iterative cycles of design, we studied the impact of 54 enzyme homologs, multiple enzyme levels, and cofactor concentrations on pathway performance. In total, we screened over 150 unique sets of enzymes in 580 unique pathway conditions to increase limonene production in 24 hours from 0.2 to 4.5 mM (23 to 610 mg/L). Finally, to demonstrate the modularity of this pathway, we also synthesized the biofuel precursors pinene and bisabolene. We anticipate that iPROBE will accelerate design-build-test cycles for metabolic engineering, enabling data-driven multiplexed cell-free methods for testing large combinations of biosynthetic enzymes to inform cellular design.

**TOC Figure:** 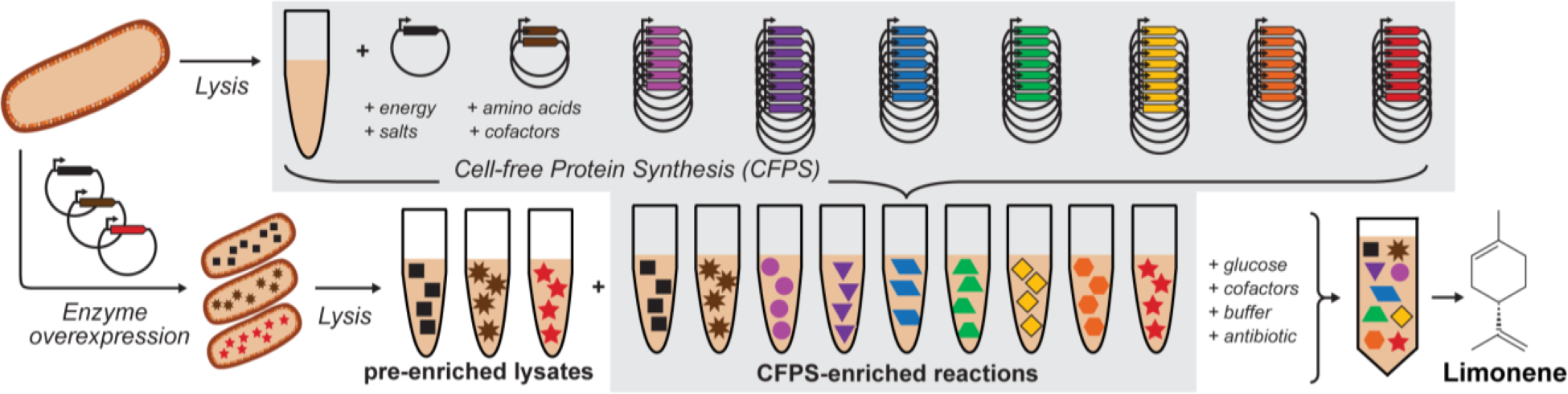

**Highlights:**

- Applied the iPROBE framework to build the nine-enzyme pathway to produce limonene
- Assessed the impact of cofactors and 54 enzyme homologs on cell-free enzyme performance
- Iteratively optimized the cell-free production of limonene by exploring more than 580 unique reactions
- Extended pathway to biofuel precursors pinene and bisabolene

## 1. Introduction

Isoprenoids are a promising class of molecules with over 40,000 known structures and potential uses as pharmaceuticals, flavors, fragrances, pesticides, disinfectants, and chemical feedstocks (Bohlmann and Keeling, 2008; George et al., 2015; Jongedijk et al., 2016). While there have been demonstrations of cellular isoprenoid production at commercial scales (Benjamin et al., 2016; Paddon et al., 2013), efforts to engineer strains for new products have proved challenging.

In particular, efforts to explore large sets of heterologous expression conditions are constrained by the need to re-engineer the cell in each iteration. This constraint has often limited cellular approaches to small sets of unique strains, with these sets focused on modifying expression conditions such as ribosome binding site strength (Li et al., 2019; Nowroozi et al., 2014), enzyme expression timing (Alonso-Gutierrez et al., 2015), and plasmid architecture (Yang et al., 2016). New developments in parallelized DNA assembly and robotic liquid handling have enabled the testing of 122 plasmid architectures for the 16-gene refactored nitrogen fixation gene cluster from *Klebsiella oxytoca* (Smanski et al., 2014), the construction of “Marionette” strains for prototyping 243 different expression profiles for lycopene pathway enzymes (Meyer et al., 2019), and the characterization of thousands of ribosome binding site combinations for tuning the production of limonene (Jervis et al., 2019) and naringenin (Zhou et al., 2019). However, these efforts have not been adapted to rapidly characterizing large combinations of enzyme homologs, which can significantly enhance performance (Ma et al., 2011; Tsuruta et al., 2009).

Cell-free metabolic engineering offers tremendous flexibility to quickly tune biosynthetic pathway enzymes, reaction substrates, and cofactors (Dudley et al., 2015; Silverman et al., 2019), giving access to test hundreds of unique pathway expression conditions. Cell-free systems using purified enzymes in particular have shown utility in prototyping metabolic pathways for a range of compounds including fatty acids (Liu et al., 2010), farnesene (Zhu et al., 2014), phenylalanine (Ding et al., 2016), and non-oxidative glycolysis (Bogorad et al., 2013). Crude lysates are becoming increasingly popular for prototyping metabolism because lysates contain endogenous metabolism, alternate substrates, and cofactors (Miguez et al., 2019; Schuh et al., 2019). Additionally, when provided with an energy source, amino acids, NTPs, and excess cofactors, crude lysates contain the translational machinery for cell-free protein synthesis (CFPS) which enables rapid production of proteins (Carlson et al., 2012; Casini et al., 2018; Chen et al., 2020; Des Soye et al., 2019; Huang et al., 2018; Jaroentomeechai et al., 2018; Jewett et al., 2008; Kightlinger et al., 2019; Lin et al., 2020; Silverman et al., 2019; Stark et al., 2019; Stark et al., 2018). CFPS decreases the time from DNA to soluble protein and can be used to synthesize functional catalytic enzymes (Karim and Jewett, 2016). The hybrid approach of cell-free protein synthesis metabolic engineering (CFPS-ME) has been successfully adapted to prototype production of polyhydroxyalkanoate (Kelwick et al., 2018), 1,4-butanediol (Wu et al., 2015; Wu et al., 2017), and styrene (Grubbe et al., 2020). Yet, even with the rapid ability to synthesize and test enzymes *in vitro*, these examples have typically utilized only a small set of enzyme homologs in their optimization strategies.

In this work, we describe a modular, high-throughput isoprenoid production platform for quickly prototyping enzyme homologs, concentrations, and reaction conditions. By expressing pathway enzymes using CFPS in separate reactions and then mixing them together in known concentrations, we modularly assemble pathway combinations for production of the monoterpenoid limonene. This conceptual approach, called *in vitro* Prototyping and Rapid Optimization of Biosynthetic Enzymes (iPROBE), has previously been shown to shorten the time to prototype 3-hydroxybutyrate and *n*-butanol production from more than 6 months to a few weeks for improving *in vivo* biosynthesis in *Clostridium* (Karim et al., 2019). Notably, iPROBE demonstrates a strong correlation between in cell and cell-free pathway performance (Karim et al., 2019).

Here we expand the iPROBE approach to longer isoprenoid pathways. We use the 9-heterologous enzyme pathway to limonene as a model pathway (**Figure 1A**). We screened 580 unique pathway combinations testing 54 different enzyme variants in several reaction (cofactor) conditions. By screening hundreds of enzyme sets and various reaction formats, we were able to improve production up to 25-fold from our initial setup. We also demonstrated pathway modularity by swapping out the isoprenoid synthetase to produce pinene and bisabolene. Our results suggest that previous cell-free isoprenoid systems, which have so far, to our knowledge reported fewer than 20 enzyme and reaction combinations per study (Chen et al., 2013; Dirkmann et al., 2018; Korman et al., 2017; Korman et al., 2014; Rodriguez and Leyh, 2014; Zhu et al., 2014), could benefit from screening more enzyme variants. iPROBE provides the ability to test dozens of enzyme homologs in hundreds of combinations without needing to re-engineer a cell or re-assemble DNA. As a result, we expect iPROBE to enhance efforts to prototype isoprenoid and other complex biosynthetic pathways for cellular or cell-free biomanufacturing.

**Figure 1.**
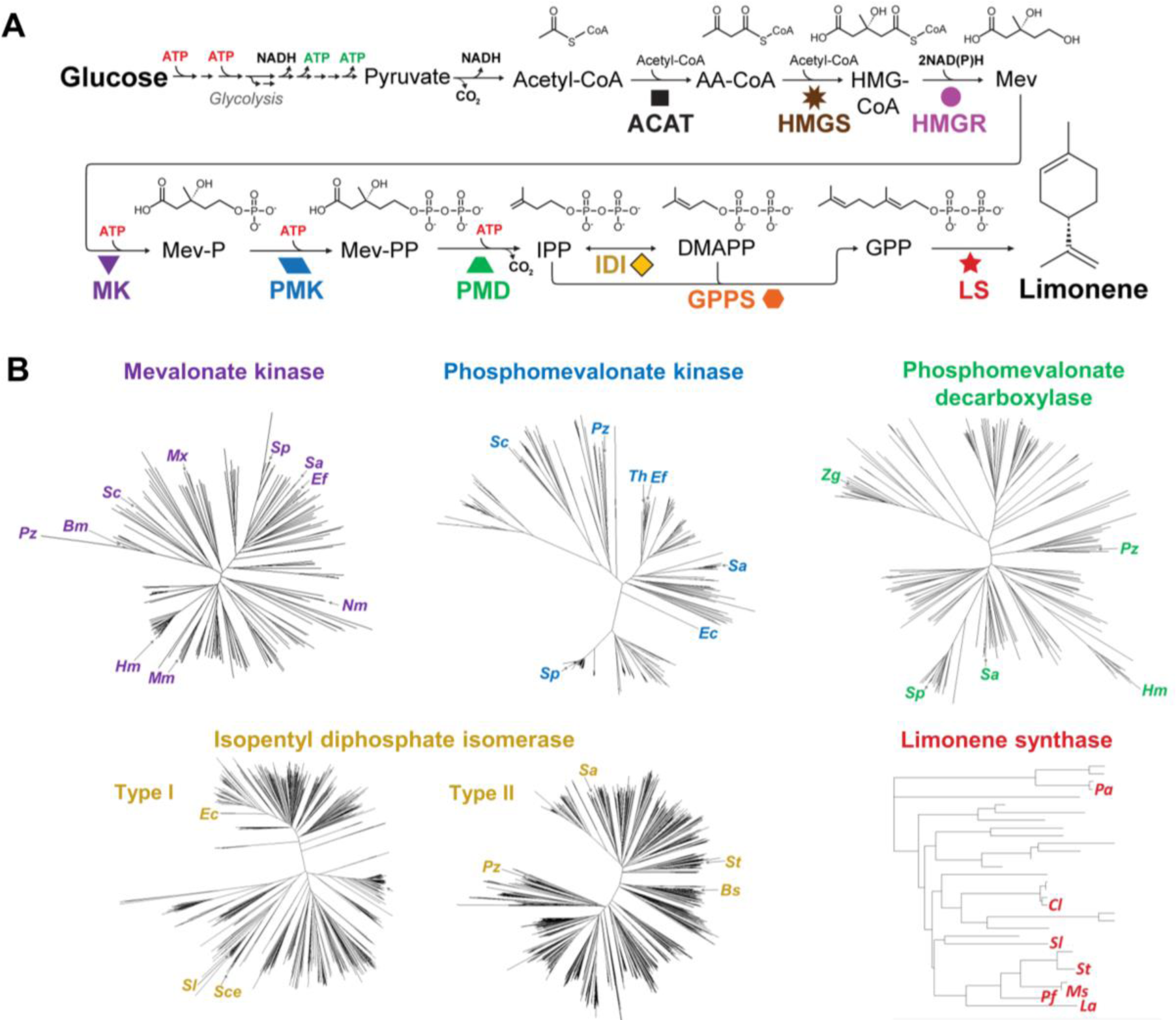
Cell-free limonene production using enzymes sourced from a variety of organisms. (A) The metabolic pathway from glucose to limonene requires nine enzymes plus glycolysis activity present in the lysate. (B) Phylogenetic comparison of enzyme sequences to select diverse enzyme homologs for testing. AA-CoA, acetoacetyl-CoA; HMG-CoA, 3-hydroxy-3-methylglutaryl-CoA; Mev, mevalonate; Mev-P, mevalonate-5-phosphate; Mev-PP, mevalonate pyrophosphate; IPP, isopentenyl pyrophosphate; DMAPP, dimethylallyl pyrophosphate; GPP, geranyl pyrophosphate; ACAT, acetyl-CoA acetyltransferase; HMGS, hydroxymethylglutaryl-CoA synthase; HMGR, hydroxymethylglutaryl-CoA reductase; MK, mevalonate kinase; PMK, phosphomevalonate kinase; PMD, pyrophosphomevalonate decarboxylase; IDI, isopentenyl pyrophosphate isomerase; GPPS, geranyl diphosphate synthase; LS, limonene synthase.

## 2. Materials and Methods

### 2.1 Strain, plasmid, and lysate preparation

All enzyme sequences tested in CFPS were cloned into the pJL1 backbone (Addgene #69496). To assemble plasmids encoding previously tested enzyme homologs (Dudley et al., 2019), the coding sequence of each enzyme was PCR-amplified using forward primer [ttaactttaagaaggagatatacatatggagaaaaaaatcNNNNNNNNNNNNNNNNNNNN where the 3’ region encodes the gene sequence starting at the second codon (i.e. right after the ATG)] and reverse primer [ttcctttcgggctttgttagcagccggtcgacNNNNNNNNNNNNNNNNNNNN where the 3’ region encodes the C-terminus of the gene sequence including the stop codon]. The forward primer adds an N-terminal expression tag to improve cell-free expression which consists of a 15 nucleotide, AT-rich sequence encoding the first five amino acids (Met-Glu-Lys-Lys-Ile, MEKKI) of chloramphenicol acetyl transferase which was used as a reporter plasmid during development of the *E. coli* CFPS system (Jewett and Swartz, 2004; Swartz et al., 2004). Addition of the MEKKI sequence demonstrably improved *in vitro* expression (**Figure S1**). The PCR fragment was then mixed with pJL1 backbone digested with the restriction enzymes NdeI and SalI along with reagents for Gibson assembly (Gibson et al., 2009).

To obtain additional sequences for testing, we first generated phylogenetic trees of pathway homologs were generated using Geneious bioinformatics software (Auckland, New Zealand). Sequences for reactions 2.7.1.36 (MK), 2.7.4.2 (PMK), 4.1.1.33 (PMD) 5.3.3.2 (IDI), and 4.3.2.3.16/19/20 (LS) were downloaded from the BRENDA database (Scheer et al., 2010) and aligned in Geneious using a Jukes-Cantor genetic distance model and a Neighbor-Joining tree build method. To choose uncharacterized homologs for testing, we selected 15 sequences from branches of the phylogenetic tree not represented by 30 previously characterized homologs (**Table S1**). These 45 sequences were then codon optimized for expression in *E. coli* synthesized by Gen9 (Cambridge, MA) or Twist Bioscience (San Francisco, CA). Sequences from Twist were delivered cloned into the pJL1 backbone. Gene sequences contained two sequential NdeI recognition sequences at the start of each gene resulting in cat<u>ATGCAT</u>ATGGAGAAAAAAATC (encoding MHMEKKI) instead of catATGGAGAAAAAAATC (encoding MEKKI). Upon comparison, the CFPS expression was equivalent for both N-terminal expression tags (**Figure S2**) and both tags ultimately used (see **Table S1**).

Lysates pre-enriched with a pathway enzyme were generated as described previously (Dudley et al., 2019). To generate CFPS S30 lysate for cell-free protein synthesis, BL21 Star(DE3) *E. coli* was grown in 1 L of 2xTYPG media in Tunair™ shake flasks at 37 °C (250 rpm). Expression of T7 TRNA polymerase was induced at OD_600_ = 0.6 by addition of 0.1 mM IPTG and cells were harvested by centrifugation at OD_600_ = 3.0. Pellets were washed twice in S30 buffer (10 mM tris acetate pH 8.2, 14 mM magnesium acetate, 60 mM potassium acetate, no dithiothreitol (DTT)), flash frozen, and stored at −80 °C. Cell pellets were then thawed on ice, resuspended in S30 buffer without DTT (0.8 mL per gram cell pellet), lysed via pressure homogenization with one pass at 20,000 psi (Avestin EmulsiFlex-B15), and centrifuged twice at 30,000 *x g* for 30 minutes. The supernatant (i.e. lysate) was transferred to a new container without disturbing the pellet, aliquoted, and flash frozen for storage at −80 °C. Optimal magnesium concentration was determined to be 8 mM based on expression of the plasmid pJL1-sfGFP (Addgene #69496) encoding superfolder Green Fluorescent Protein (sfGFP).

### 2.2 Cell-free protein synthesis reactions

All CFPS reactions used a modified PANOx-SP formula (Jewett and Swartz, 2004; Kwon and Jewett, 2015). Each reaction contains 13.3 μL of S30 extract for a 50 μL CFPS reaction (**Figure S3**) in addition to ATP (1.2 mM), GTP, UTP, CTP (0.85 mM each), folinic acid (34 µg/mL), *E. coli* tRNA mixture (170 µg/mL), 20 standard amino acids (2 mM each), NAD^+^ (33 mM), coenzyme A (CoA) (0.27 mM), oxalic acid (4 mM), spermidine (1.5 mM), putrescine (1 mM), HEPES (57 mM), potassium glutamate (134 mM), ammonium glutamate (10mM), magnesium glutamate (8 mM), phosphoenolpyruvate (PEP) (33 mM), and plasmid (13.3 µg/mL) encoding the metabolic pathway enzyme. T7 RNA polymerase was not added to CFPS reactions since extracts were induced with IPTG during cell growth. CFPS reactions of pathway enzymes are incubated for 20 hours at 30 °C or 16 °C (**Table S1**), flash frozen on liquid nitrogen, and stored at −80 °C.

### 2.3 Quantification of CFPS protein production using radioactive amino acid incorporation

To measure the amount of protein produced in a CFPS reaction,^14^C-leucine (10 µM) was supplemented in addition to the 20 standard amino acids. Reactions were centrifuge at 21,000 x *g* for 10 minutes to pellet insoluble proteins for measurement of soluble protein. Reactions were quenched by addition of equal volume 0.5 M potassium hydroxide and pipetted onto two separate 96-well fiberglass papers (PerkinElmer Printer Filtermat A 1450-421, 90×120mm) and dried. One paper was subjected to three trichloroacetic acid (TCA) washes (15 min each at 4 °C) to precipitate proteins and dried after rinsing with 100% ethanol. Scintillation wax (PerkinElmer MeltiLex A 1450-441 73×109 mm) was applied and radioactivity was measured using liquid scintillation counting via a MicroBeta2 (PerkinElmer, Waltham, MA). Protein concentration was determined as previously described (Jewett, 2004; Jewett et al., 2008; Jewett and Swartz, 2004) by comparing radioactivity in the total reaction to that of precipitated protein using Equation 1 (**Appendix S1**).

### 2.4 Mixing of CFPS reactions and pre-enriched lysates to produce limonene

All limonene synthesis reactions are 30 μL in total volume and can be divided into the “CFPS fraction” and the “substrate/lysate/cofactor fraction”. The 15 μL CFPS fraction contains six to nine CFPS reactions (thawed on ice) and mixed at concentrations dictated by the experiment. “Blank” CFPS reaction (50 μL volume containing all CFPS reagents, no plasmid, and incubated for 20 h at 30 °C, **Figure S4**) is added until the total volume of CFPS fraction is 15 µL. The 15 μL “substrate/lysate/cofactor fraction” includes the following components (concentrations are given with respect to the fully assembled 30 μL limonene synthesis reaction): 4 mM magnesium glutamate, 5 mM ammonium glutamate, 65 mM potassium glutamate, 200 mM glucose, 50 µg/mL kanamycin, and 100 mM Bis-Tris (pH 7.4). Since “CFPS fraction” already contains glutamate salts, supplementation of glutamate salts in the “substrate/lysate/cofactor fraction” maintains the overall cation concentrations at 8 mM magnesium, 10 mM ammonium, and 130 mM potassium. Optionally, cofactors ATP, NAD^+^, and CoA can be supplemented as well (concentrations are given with respect to the fully assembled 30 μL limonene synthesis reaction). The “substrate/lysate/cofactor fraction” also includes fresh S30 lysate(s) at a total protein concentration of 4 mg/mL. Total protein is measured Bradford assay with bovine serum albumin (BSA) as the standard using a microplate protocol (Bio-Rad, Hercules, CA). Typically, three S30 lysates are added each pre-enriched for a single enzyme (*Ec*ACAT, *Sc*HMGS, and *Ms*LS) (Dudley et al., 2016; Dudley et al., 2019); each is included at a total protein concentration of 1.33 mg/ mL. Based on the level of protein overexpression measured by densitometry (**Table S2**), the final concentration of enriched enzyme in the limonene synthesis reaction is estimated to be ∼4.7 μM for *Ec*ACAT, ∼5.5 μM for *Sc*HMGS, and ∼3.4 μM for *Ms*LS, respectively. “Blank lysate” (generated from BL21(DE3) containing no expression plasmid) is added in place of an enzyme-enriched lysate if the given enzyme is included as a CFPS reaction. Limonene synthesis reactions are incubated at 30 °C with a 30 µL dodecane overlay. Limonene in the overlay was quantified at 3, 4, 5, 6, and 24 hours using GC-MS as previously described (Dudley et al., 2019). See **Table S3** and **Supplementary Data File 1** for systematic description of the variable reaction components in each enzyme set tested. The TREE score is calculated using Equations 2 and 3 described in **Appendix S1**.

## 3. Results

The vision of this work was to (1) apply the iPROBE framework to build the nine-enzyme pathway to produce limonene, (2) assess the impact of cofactors on cell-free enzyme performance, (3) iteratively optimize the enzymatic production of limonene, and (4) extend the pathway to additional isoprenoids. We selected limonene biosynthesis as our model system given its importance as a high-value specialty chemical (Jongedijk et al., 2016) and our previous work with the pathway. In our previous effort, we demonstrated cell-free biosynthesis of limonene in crude cell lysates whereby the enzymes were heterologously overexpressed *in vivo* before lysate preparation (Dudley et al., 2019) (**Figure 1A**). Crude cell lysates contain endogenous enzymes for glycolysis that regenerate NADH and convert glucose to acetyl-CoA, providing the starting intermediate for limonene biosynthesis. Upon mixing together of selectively enriched lysates comprising all necessary enzymes, the full pathway could be activated. This work gave a starting point for applying iPROBE, wherein we would utilize lysates enriched not by heterologous protein expression prior to lysis, but by expressing the protein from a DNA template in the reaction via CFPS.

### 3.1 Application of iPROBE to a nine-enzyme pathway for producing limonene

We established a cell-free pathway to produce limonene and identified candidate sets of enzyme homologs to improve production. First, we built a cell-free system to produce limonene using a previously characterized set of enzymes with the iPROBE approach, where each enzyme is produced individually via CFPS and the resulting “CFPS-enriched reactions” are mixed with glucose substrate and cofactors to produce the target molecule limonene (**Figure 2A**). Nine enzymes were needed including acetyl-CoA acetyltransferase (ACAT) from *E. coli*, hydroxymethylglutaryl-CoA synthase (HMGS) from *Saccharomyces cerevisiae*, hydroxymethylglutaryl-CoA reductase (HMGR) from *Pseudomonas mevalonii*, mevalonate kinase (MK), phosphomevalonate kinase (PMK), and pyrophosphomevalonate decarboxylase (PMD) from *S. cerevisiae*, isopentenyl pyrophosphate isomerase (IDI) from *E. coli*; geranyl diphosphate synthase (GPPS) from *Picea abies*, and limonene synthase (LS) from *Mentha spicata*. (Dudley et al., 2019). This initial enzyme set 1.0 produced 0.17 mM ± 0.007 mM limonene (**Figure S5**), demonstrating the ability of the cell-free approach to reconstitute the biosynthesis pathway from cell-free synthesized enzymes.

**Figure 2.**
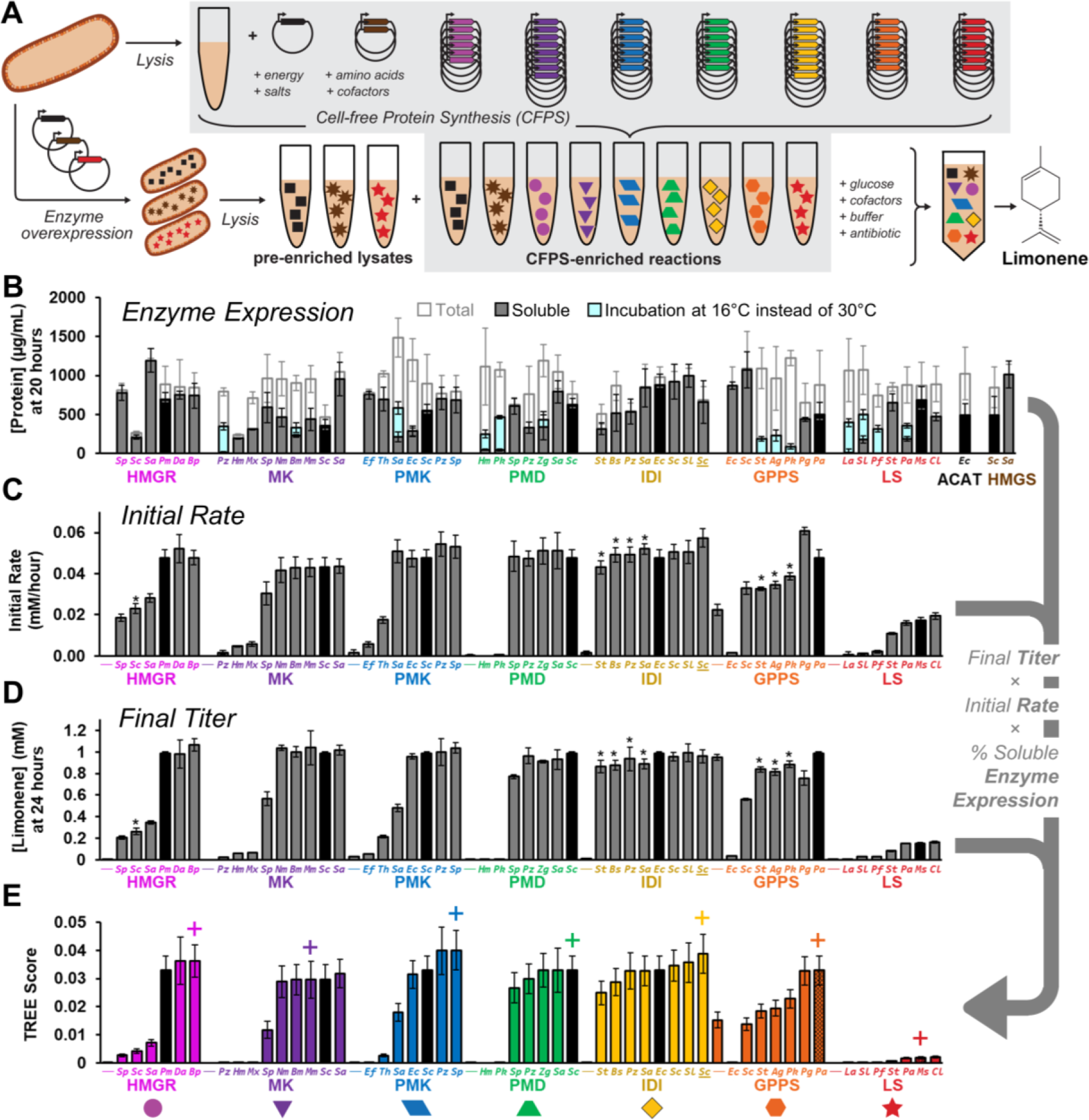
Cell-free expression of enzyme homologs and initial measurement of limonene production. (A) Cell-free protein synthesis (CFPS) generates “CFPS-enriched reactions” where each individual reaction expresses a single enzyme homolog. Additionally, enzymes can be overexpressed *in vivo* and the cell subsequently lysed to generate “pre-enriched lysates”. Mixing of glucose substrate and cofactors with enzyme-enriched extracts produces limonene. (B) CFPS yields for 54 different enzymes. CFPS reactions with soluble/total protein below 30% were incubated at 16 °C instead of 30 °C; the amount of soluble protein increased for all 14 proteins. Values represent averages (n=3) and error bars represent 1 standard deviation. (C-D) Cell-free limonene production. Using various enzyme homologs generated by CFPS, the initial rate from 3-6 hours (C) and titer at 24 hours (D) of limonene were measured. The standard reaction includes 1.0 μM *Pm*HMGR, 0.4 μM *Sc*MK, 0.4 μM *Sc*PMK, 0.4 μM *Sc*PMD, 2.0 μM *Ec*IDI, and 3.0 μM *Pa*GPPS plus pre-enriched lysates for *Ec*ACAT, *Sc*HMGS, and *Ms*LS (enzyme set 1.0). Unless noted by (*, see **Table S3** for details), each enzyme homolog is substituted for the standard enzyme at the same concentration. LS homologs are generated via CFPS and compared at 1.0 μM in a reaction lacking the *Ms*LS pre-enriched lysate. Reactions labeled (−) are negative controls without the designated enzyme. The initial rate value is the slope of a linear regression of four data points at 3, 4, 5, and 6 hours and error bars are the associated standard error (see **Appendix S1**). The values for limonene at 24 hours represent averages (n=3) and error bars represent 1 standard deviation. (E) the TREE (Titer, Rate, and Enzyme Expression) score is an empirically generated aggregate value of enzyme solubility, initial rate, and final titer that enables ranking of enzyme homologs. The (+) symbol represents homologs included in enzyme set 2.0. Error bars represent propagated error from the three components (see **Appendix S1**).

We next wanted to use iPROBE to rapidly characterize combinations of enzyme homologs, which can significantly enhance performance because catalytic rate, substrate specificity, and feedback inhibition can vary widely across related enzymes in both primary (Maeda, 2019) and secondary (Schmidt et al., 2018) metabolism. Therefore, we generated phylogenetic trees for several pathway steps using all available sequences in the BRENDA enzyme database (Scheer et al., 2010) (**Figure 1B**). We then selected 45 sequences encoding homologs of the nine enzymes of the mevalonate pathway to limonene (**Appendix S2**) and assembled them into plasmids for cell-free expression. Half of the selected homologs included uncharacterized enzymes selected from diverse clades while biasing the selection towards homologs suspected to be interesting (*e.g*., the MK, PMK, PMD, and IDI sequences from *Paracoccus zeaxanthinifaciens* ATCC 21588, a marine bacteria that produces high levels of the isoprenoid zeaxanthin (Berry et al., 2009; Berry et al., 2003)). The other half were kinetically characterized or used in previous metabolic engineering efforts (**Table S1**).

Fifty-one different pathway combinations of the candidate enzymes selected were assembled using iPROBE and screened for limonene production. CFPS generated soluble protein (>30% of total protein) for 40 of the 54 enzymes (**Figure 2B**). By decreasing the CFPS incubation temperature from 30 °C to 16 °C, we increased soluble protein yields for the remaining 14 enzymes. To test metabolic activity, iPROBE reactions were assembled containing the enzyme homolog of interest plus the remaining set of initial base-case pathway enzymes (enzyme set 1.1, which includes pre-enriched lysates of *Ec*ACAT, *Sc*HMGS, and *Ms*LS (**Figure S6**)). We measured limonene initial rate from 3 to 6 hours (**Figure 2C**) and final titer at 24 hours (**Figure 2D**) and then ranked each of these 51 different pathway combinations based on a combined Titer, Rate, and Enzyme Expression (TREE) score (**Figure 2E**). This score is a simple, empirical, aggregate value of final titer at 24 hours (mM), initial rate (mM/h), and average solubility of 9 expressed enzymes (%) (**Appendix S1**; (Karim et al., 2019)). From these data, we observed equivalent or improved activity relative to the initial base-case homolog for 2 HMGR, 5 MK, 4 PMK, 4 PMD, 7 IDI, 5 GPPS, and 3 LS homologs. This approach also served as a quick way to rule out several homologs of MK, PMK, PMD, and LS with much lower (or zero) activity; these homologs were excluded from further testing. The relative activity of different HMGRs is similar to our previous study using pre-enriched lysates (Dudley et al., 2016), as well as *in vivo* pathway testing (Ma et al., 2011) which suggests that expression of enzymes via CFPS produces proteins with similar properties to those generated with these other methods.

### 3.2 Cofactors are a key parameter in testing multi-enzyme pathways

We next wanted to compare the best enzymes from the activity screen to enzyme set 1.1. To do this, we selected six homologs as best candidates using the TREE score metric (**Figure 2E**) for enzyme set 2.0: HMGR from *Bordtella petrii*, MK from *Methanosarcina mazei*, PMK from *Enterococcus faecalis*, PMD from *S. cerevisiae*, IDI from *Streptomyces clavuligerus*, and GPPS from *Picea abies* (Norway spruce). Each selected homolog was highest performing in the TREE score with the exception of *Mm*MK, which was chosen over similarly performing homologs due to its known lack of inhibition from downstream isoprenoid metabolites (Primak et al., 2011). We then compared the initial enzyme set 1.1 to the improved set 2.0 containing new homologs for HMGR, MK, PMK, and IDI over 96 hours and found that enzyme set 2.0 did not improve limonene production (**Figure 3A**). While the first round of enzyme screening was successfully able to identify active and inactive enzyme homologs, there were only small differences between active homologs (**Figure 2E; Figure 3A**).

**Figure 3.**
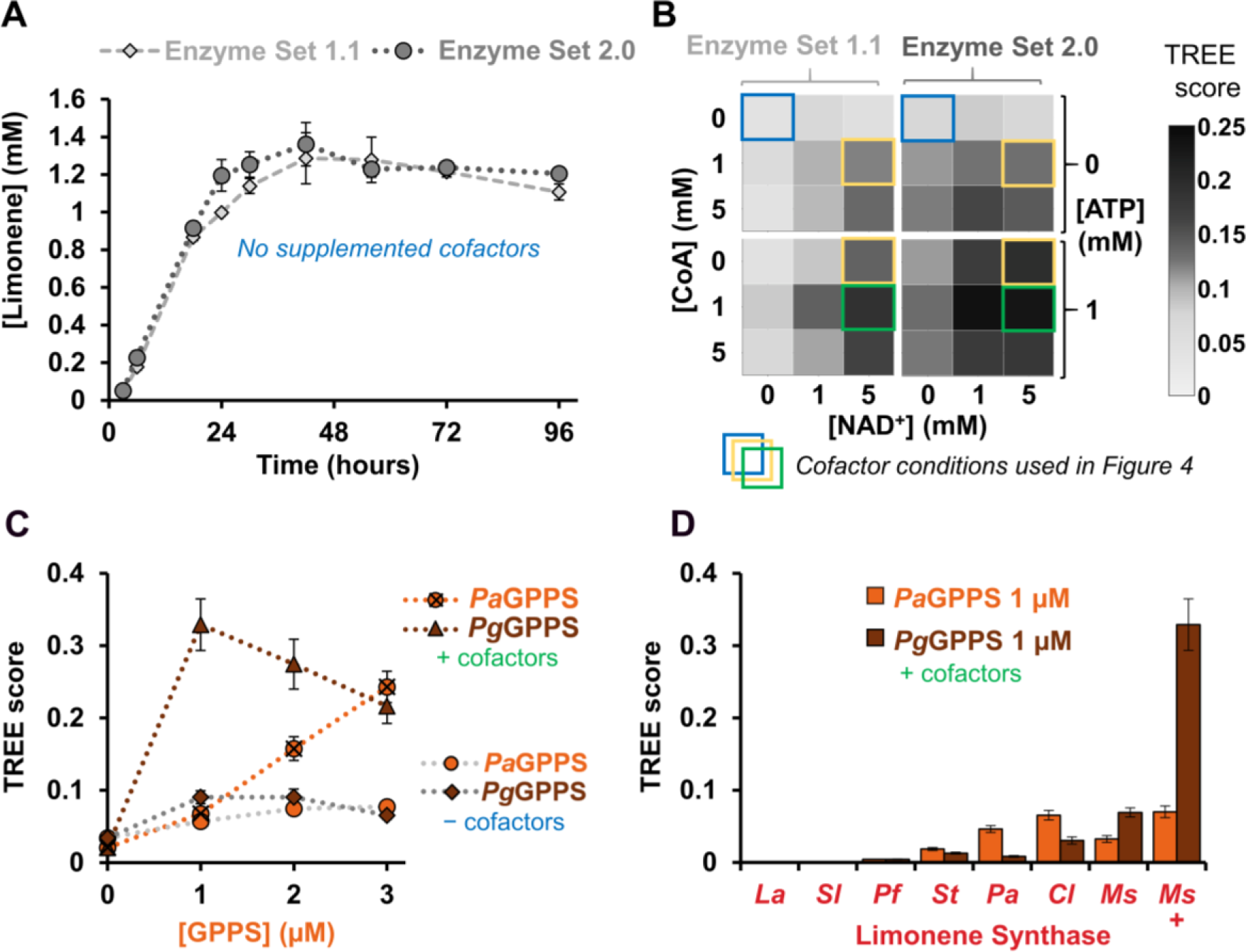
Using a new enzyme homolog set and optimizing cofactor concentrations accentuates differences between GPPS and LS homologs. (A) Comparison of initial enzyme homologs (enzyme set 1.1) and an improved set 2.0 containing new homologs for HMGR, MK, PMK, and IDI. Values represent averages (n=3) and error bars represent 1 standard deviation. (B) Supplementation of cofactors NAD^+^, CoA, and ATP improves limonene rate, titer, and TREE score (**Figure S7**) compared to no additional cofactors (blue box). Four reaction conditions (in blue, yellow, and green boxes) were selected as cofactor conditions for further experiments. (C) TREE scores of *Pa*GPPS and *Pg*GPPS (the best two GPPS homologs in **Figure 2E**) under different enzyme concentrations. These data informed the reduction of GPPS to 1.0 μM for enzyme set 2.2. (D) Re-testing of LS homologs derived from CFPS at a higher concentration (1.8 μM) and using supplemented cofactors. Values represent TREE scores and error bars represent propagated error (see **Appendix S1**).

We next sought to increase cell-free limonene synthesis by optimizing cofactors and tuning enzyme concentrations. Cofactors are important parameter in optimizing the performance of *in vitro* enzyme pathways (Dudley et al., 2016; Karim et al., 2018; O’Kane et al., 2019). We have previously shown 100-fold differences in HMG-CoA concentrations across 768 unique cofactor conditions of ATP, NAD^+^, and CoA (O’Kane et al., 2019). Therefore, we tested enzyme sets 1.1 and 2.0 in 18 different cofactor conditions (**Figure 3B**). During this optimization, we found that enzyme set 2.0 is better than or equivalent to enzyme set 1.1 in all cofactor conditions but the improvements are more pronounced at optimized cofactor conditions. The best conditions with enzyme set 2.0 increased the TREE score by two-fold with a limonene initial rate and titer at 24 hours of 0.113 ± 0.010 mM/hour and 1.73 ± 0.23 mM, respectively (**Figure S7**). Next, we wanted to reduce the concentration of enzymes to leave additional volumetric space in the reaction for other CFPS-derived enzymes and potentially differentiate less-active homologs from more-active ones. We titrated the concentration of each CFPS-derived pathway enzyme individually using enzyme set 2.0 (**Figure S8**) and found that we could reduce the concentrations of HMGR, MK, PMK, PMD, and IDI (enzyme set 2.1) to produce equivalent levels of limonene (**Figure S8G**).

We next examined GPPS, a key enzyme that directs metabolic flux towards ten-carbon GPP and competes with native metabolism which typically generates fifteen-carbon farnesyl pyrophosphate. At high protein concentrations (3 μM), *Pa*GPPS from *Picea abies* produces the highest limonene final titer, however *Pg*GPPS from *Picea glauca* has a higher productivity from 3-6 hours and produces the best TREE score at a lower concentration (1 μM) (**Figure 3C; Figure S9**). We hypothesize that *Pg*GPPS has higher catalytic rate than *Pa*GPPS but is less stable and is inactivated as the reaction proceeds. We therefore reduced the concentration of GPPS from 3.0 μM (Set 2.1) to 1.0 μM (Set 2.2) for further experiments. With increased next retested all LS homologs at a higher concentration (particularly those that did not express well in the cell-free system). The pre-enriched lysate of *Ms*LS from *Mentha spicata* again produces the highest TREE score but the difference was far more pronounced when using *Pg*GPPS rather than *Pa*GPPS (**Figure 3D; Figure S10**). This result highlights the importance of testing enzyme homologs in a variety of pathway contexts.

### 3.3 Iterative screening of active enzyme homologs under multiple cofactor conditions

To demonstrate the full potential of iPROBE we tested all active enzyme homologs in an iterative experimental approach that characterized 102 enzyme sets tested in four different cofactor concentrations for a total of 408 unique combinations (**Figure 4**). The four different cofactor conditions explore different concentrations of CoA, ATP, and NAD^+^. At this point, the primary experimental limitation was not enzyme expression or pathway assembly but product quantitation via GC-MS; this prohibited testing the full combinatorial space of 6 * 8 * 5 * 5 * 6 * 4 * 2 = 57,600 enzyme sets, which could have otherwise been achieved with automated liquid handling and assembly of cell-free biosynthesis units. The iterative testing approach started by substituting each the six active GPPS homologs and measuring the TREE score under four different cofactor conditions (24 unique conditions; Round 3.x) (**Figure 4A**). In an effort to capture enzymes that were most active under multiple cofactor conditions, the two enzyme sets with the highest average TREE score over four cofactor concentrations (**Figure 4B; Figure S11**) were selected to move forward in the screen. These sets, differing in GPPS homolog, were then used to test the eight active IDI homologs combinatorially. We assembled 64 unique pathway conditions (enzyme sets and cofactor conditions; Round 4.x) and measured TREE scores for each (**Figure 4B; Figure S11**). Four enzyme sets were selected and carried forward to be the context for combinatorial testing of all PMD homologs (80 unique conditions; Round 5.x). Thereafter, each experiment used the best scoring five conditions from the previous iteration to test active PMK (Round 6.x), PMD (Round 7.x), and HMGR (Round 8.x) homologs sequentially.

**Figure 4.**
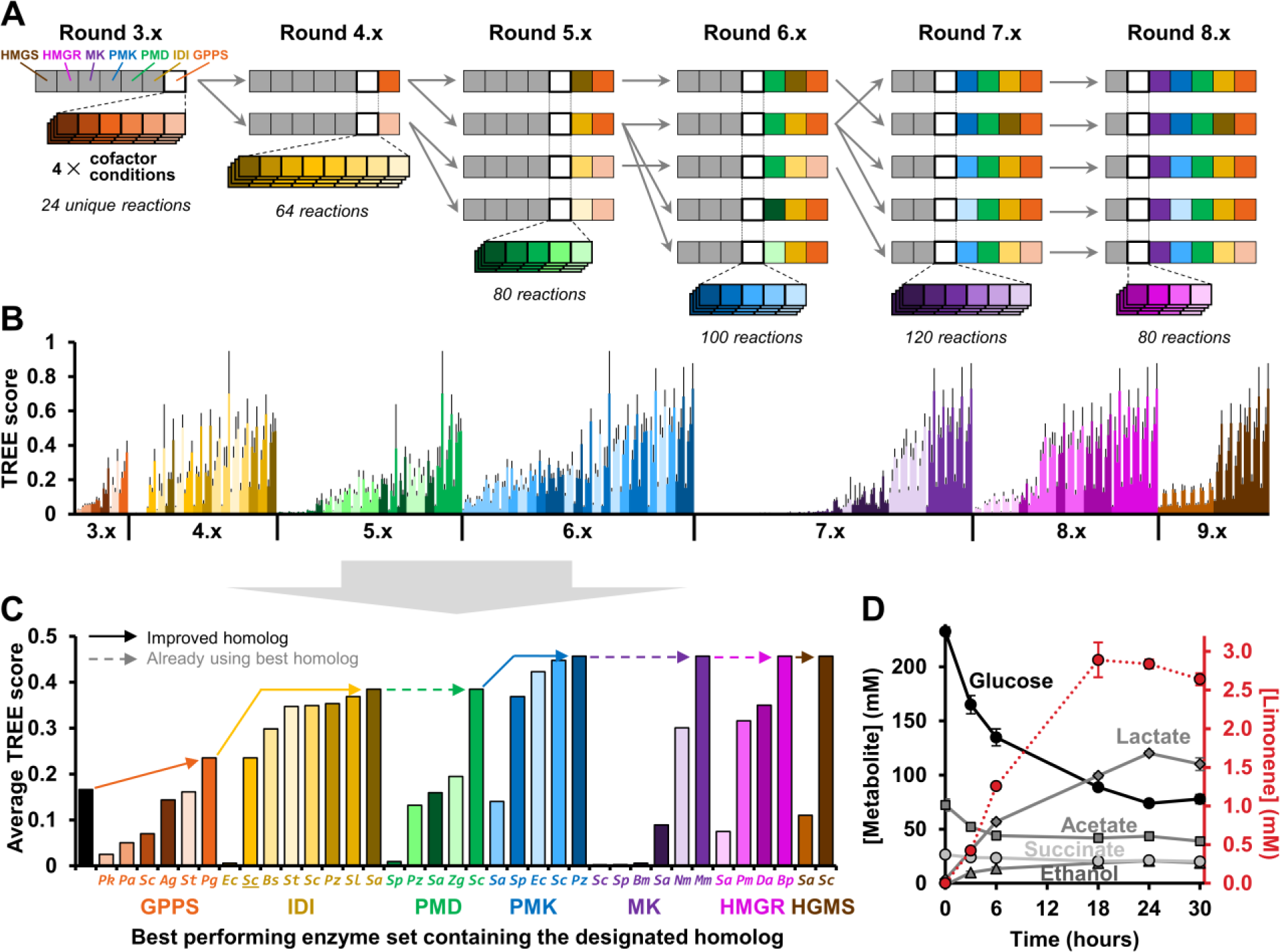
Iterative testing of enzyme sets at four different cofactor conditions identifies homologs with robust activity. (A) Conceptual approach: Starting with enzyme set 2.2, active homologs of GPPS were substituted for the default GPPS (*Pa*GPPS). Each GPPS homolog (orange) was tested under four different cofactor conditions and the resulting TREE scores averaged to rank each enzyme set. The top two sets were carried forward into the next iterative experiment in which all active homologs of IDI (yellow) were tested. The top two sets plus the best performing *St*GPPS sets were again carried forward and all active PMD homologs (green) were tested. Each subsequent iterative experiment used the top five conditions from the previous iteration to test active PMK (blue), PMD (purple), and HMGR (magenta) enzymes in sequence. (B) TREE scores of 408 reactions where each bar is a unique cofactor/enzyme set. Error bars represent propagated error (see **Appendix S1**) (C) The TREE score of each enzyme set was averaged across all four cofactor conditions and the highest score plotted to compare each homolog as a single value. The black bar represents the starting enzyme set 2.1. (D) Production of limonene (mM) and other metabolites over the course of the reaction. The concentration of each enzyme is 0.2 μM HMGR, 0.1 μM MK, 0.2 μM PMK, 0.2 μM PMD, 0.2 μM IDI, and 1.0 μM GPPS plus pre-enriched lysates for *Ec*ACAT, *Sc*HMGS, and *Ms*LS; see **Table S3** for more detailed description of the reaction components. Values represent averages (n=3) and error bars represent 1 standard deviation.

The higher limonene production and reduced enzyme concentrations proved to be a more stringent screen for comparing enzyme homologs relative to **Figure 2**. The iterative experiment found clear differences between MK and PMD homologs that produced the same amount of limonene in earlier experiments (**Figure 4C, Figure S12**). Additionally, we found that *Ec*IDI from *E. coli* (one of the most common enzymes used for *in vivo* terpenoid production) is far less active at reduced concentration compared to other IDI homologs. Finally, we wish to highlight that some enzyme sets showed a variable ranking depending on which cofactor concentration was used. For example, *Sa*HMGR, *Sa*MK, and *Pz*PMK prefer no supplemental CoA while *St*GPPS is highly active under optimal cofactors but quite low without cofactor supplementation (**Figure S13**). The final enzyme set 9.0 included *Ec*ACAT, *Sc*HMGS, *Bp*HMGR, *Mm*MK, *Pz*PMK, *Sc*PMD, *Sl*IDI, *Pg*GPPS, and *Ms*LS (**Figure 4D**). Relative to our baseline enzyme set 1.1, the final conditions improved limonene initial rate and final titer by 6-fold and 4-fold, respectively.

### 3.4 Bioproduction of additional terpenoids

With an optimized set of enzymes at hand, we aimed to demonstrate the modularity and flexibility of the cell-free system by extending the pathway to additional products. To achieve this goal, we replaced the final enzyme of the pathway with different synthases to generate the biofuel precursors pinene and bisabolene (**Figure 5A**). Pinene is a useful precursor molecule for various perfumes and bisabolene can be chemically hydrogenated to bisabolane which has nearly identical properties to D2 diesel fuel (Peralta-Yahya et al., 2011). We compared two pinene synthase homologs which produce similar levels of pinene while increased ratios of β-pinene compared to previous *in vitro* and *in vivo* studies (**Figure S14**). When using 3.8 µM of CFPS-derived monoterpene synthase, the limonene synthase produces higher amounts of product compared to the pinene synthases (**Figure 5B**). A cell-free system composed of two pre-enriched lysates and seven CFPS-enriched reactions (including 1.0 µM *Ec*GPPS/FPPS (ispA) and 1.6 µM *Ag*BS) produces 4.94 ± 0.73 mM (1010 mg/L) bisabolene after 72 hours (**Figure 5C, Figure S14**). This is similar to the best bisabolene titer achieved in cells (912 mg/L) (Peralta-Yahya et al., 2011).

**Figure 5.**
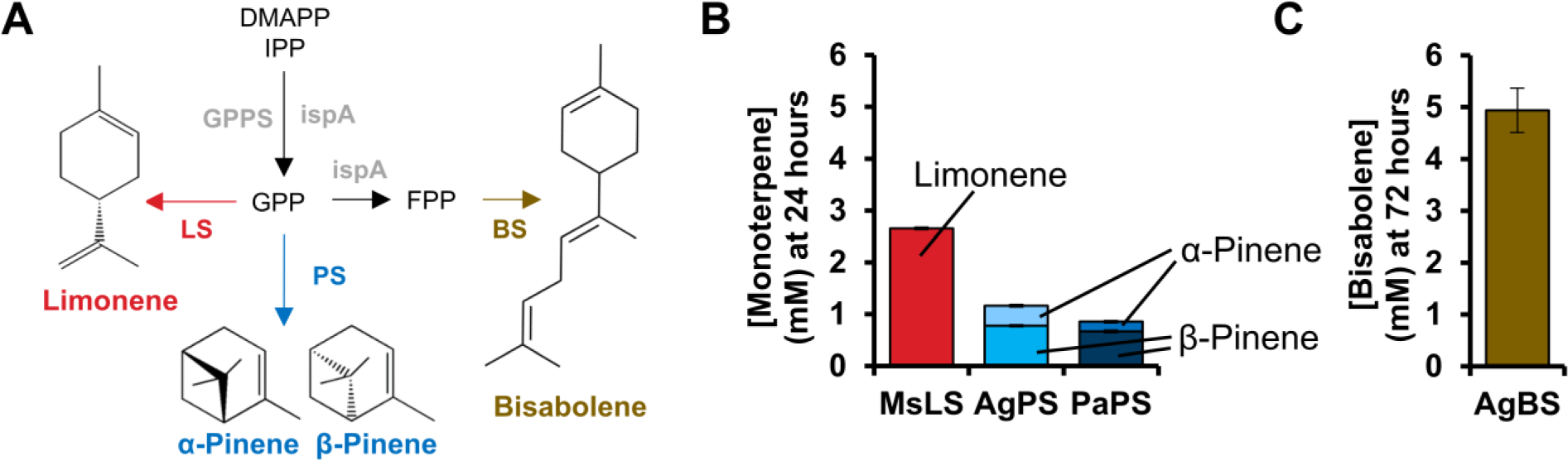
Synthesis of additional terpenes including biofuel precursors pinene and bisabolene. (A) Various terpene synthases can produce different products from the common GPP precursor. (B) Comparison of limonene and pinene production from glucose using CFPS-derived terpene synthases. Pinene synthases (PS) were encoded by sequences from *Abies grandis* (Grand fir) and *Picea abies* (Norway spruce). (C) Production of bisabolene from glucose by CFPS-enriched reactions containing farnesyl diphosphate synthase (EcFPPS encoded by *ispA*) from *E. coli* and bisabolene synthase (BS) from *Abies grandis*. Limonene, pinene, and bisabolene synthesis reactions use enzyme set 10.0 (see **Table S3**) and are supplemented with 5 mM NAD^+^, 1 mM CoA, and 1 mM ATP. Values represent averages (n=3) and error bars represent 1 standard deviation. **Figure S14** depicts the accumulation of terpenoids over time.

## Discussion

In this work, we demonstrated the potential of iPROBE to accelerate Design-Build-Test cycles for metabolic engineering. We applied iPROBE to the limonene biosynthesis pathway, representing the longest pathway utilized by iPROBE (nine steps) to date. We tested over 580 conditions of different enzyme homologs, enzyme concentrations, and cofactor conditions; this dataset is available in **Supplementary Data File 1**. The best performing reaction (using enzyme set 9.0) improved production of limonene by 25-fold from our initial conditions (enzyme set 1.0), producing 4.49 ± 0.14 mM (610 mg/L) limonene in 24 hours (**Figure S14**). These rates and titers are similar to recent *in vivo* efforts (**Table S4**), though lower than a cell-free system utilizing purified enzymes that lacks competing side reactions (Korman et al., 2017).

Our results have several key features. First, our rapid, cell-free framework facilitates the study of large number of pathway combinations. Already, cell-free prototyping has proved useful for optimizing several enzymatic pathways including isoprenoids (Dudley et al., 2016), fatty acids (Liu et al., 2010), farnesene (Zhu et al., 2014), phenylalanine (Ding et al., 2016), non-oxidative glycolysis (Bogorad et al., 2013), polyhydroxyalkanoates (Kelwick et al., 2018), 1,4-butanediol (Wu et al., 2015; Wu et al., 2017), styrene (Grubbe et al., 2020), and 3 hydroxybutyrate/n-butanol (Karim et al., 2019). Here, we used CFPS to express and study numerous enzyme homologs. The CFPS approach permitted production of active enzyme in hours without requiring protein purification or extensive expression optimization. This enabled the creation of multiple cell-free biosynthesis “units” that could be assembled modularly, in a mix-and-match fashion, to explore hundreds of distinct biosynthetic pathways. As a result, we found several enzyme steps including GPPS, IDI, and MK that showed strong differences between homologs. Specifically, we identified *Pg*GPPS, *Sl*IDI, and *Mm*MK (**Figure S11**) as promising candidates for further efforts to improve *in vivo* isoprenoid titers. Supporting our discoveries, two recent efforts utilized *Mm*MK in place of *Sc*MK to produce 1.29 g/L limonene (Wu et al., 2019) and to improve isoprene titers 1.4-fold (Li et al., 2019).

Another important feature of our work is that we identified enzyme sets with high activity across a range of cofactor concentrations. A key advantage of cell-free systems is that they enable the experimenter to directly manipulate the system, bypassing limitations on molecular transport across the cell wall. We used this feature to perform pathway optimizations under different cofactor concentrations to select enzymes that were robust and active under multiple conditions. To our knowledge, this kind of iterative, combinatorial strategy has yet to be pursued. While trends of relative homolog activity are similar across different cofactor conditions, there are exceptions such as *Pz*PMK, *Sa*MK, *Sa*HMGR, and *St*GPPS (**Figure S11; Figure S13**). Coordinately tuning enzyme homologs, enzyme concentrations, and cofactor concentrations together using data-driven design could facilitate improved pathway performance in metabolic engineering campaigns (Karim et al., 2019). Additional studies in multiple, distinct pathways, guided by machine learning could elucidate new approaches toward efficient engineering efforts.

Future developments will seek to build upon and improve our efforts to design and optimize biosynthetic pathways using iPROBE. For example, correlations between cell-free optimized pathways and cellular designs could be established. In addition, enhancements to the TREE score, which consolidates three performance parameters (final titer, initial rate, and enzyme solubility) into a single metric, are warranted. While the TREE score provides an easy-to-use strategy for pathway ranking, it is sensitive to small changes in initial rate and the effect of enzyme solubility is relatively small. Thus, refining the TREE score (*e.g*., by weighting each factor) to better correlate *in vitro* and *in vivo* data will help identify the minimal amount of cell-free data needed to inform pathway design. Finally, our isoprenoid prototyping platform could be adapted for alternate pathway routes that utilize neryl pyrophosphate as a substrate for monoterpene synthases rather than GPP (Cheng et al., 2019; Ignea et al., 2019; Wu et al., 2019) or produce IPP and DMAPP using prenol or isoprenol as an intermediate (Chatzivasileiou et al., 2019; Clomburg et al., 2019; Lund et al., 2019; Ward et al., 2019).

Looking forward, we anticipate that cell-free prototyping will facilitate rapid design-build-test cycles for identifying the best sets of enzymes to carry out a defined molecular transformation. In addition, the iPROBE framework could be adapted to other pathways for screening enzyme activity, testing for interacting effects between component parts, and analysis of byproduct pathways. iPROBE is particularly well suited for this endeavor because it can assess large numbers (500+) of enzyme combinations and cofactor conditions without requiring large-scale DNA assembly or metabolic engineering of living cells. It is also able to identify potential synergies between enzymatic steps that can only come from studying sets of enzymes in the context of the full biosynthetic pathway. These abilities will be synergistically enhanced as new methods are developed for measuring target metabolites at high throughput (O’Kane et al., 2019). The approach reported here will advance efforts to design, build, and test any metabolic pathway of choice.

## Author contributions

QMD and MCJ designed the study; QMD, ASK, and MCJ developed the cell-free framework; CJN and QMD assembled the plasmids, QMD performed all cell-free experiments; QMD and ASK analyzed cell-free data; QMD, ASK, and MCJ wrote the manuscript.

## Acknowledgements

We gratefully acknowledge the Department of Energy (BER grant: DE-SC0018249), the Joint Genome Institute Community Science Program (Project 503280), the David and Lucile Packard Foundation (2011-37152), and the Dreyfus Teacher-Scholar Program for funding and support. QMD is funded, in part, by the Northwestern Molecular Biophysics Training Program supported by NIH via NIGMS (5T32 GM008382). We also thank Will Bothfeld for helpful discussions and Do Soon Kim for advice on developing supplemental figures.

