## Supplemental Information for "Cell-free prototyping of limonene biosynthesis using cell-free protein synthesis"

#### Supplementary Information

Supplementary Tables S1-4
Supplementary Figures S1-17
Supplementary References
Appendix S1 (equations)
Appendix S2 (DNA sequences)

### Supplementary Tables

#### **Table S1.** DNA sequences and plasmids used in this study.

| Enzyme | # | Abbr. | Source organism | uniprot ID | Plasmid name | N-terminal tag | Molecular weight (Da) | CFPS expression temperature | Reference |
|---|---|---|---|---|---|---|---|---|---|
| EcACAT | 1 | Ec | *Escherichia coli* | P76461 | 354_pJL1-atoB (ACAT_Eco) | none | 40607 | 30 °C | Dueber *et al. Nat Biotech* (2009), Dudley *et al. ACS Syn Bio* (2016), many others |
| ScHMGS | 2a | Sc | *Saccharomyces cerevisiae* | P54839 | 310_pJL1-(CAT5aa)-HMGS_Sce | MEKKI | 55756 | 30 °C | Dueber *et al. Nat Biotech* (2009), Dudley *et al. ACS Syn Bio* (2016), many others |
| SaHMGS | 2b | Sa | *Staphylococcus aureus* | A0A114PFY9 | 311_pJL1-(CAT5aa)-HMGS_Sau | MEKKI | 43706 | 30 °C | Tsuruta *et al. PloS One* (2009), Dudley *et al. ACS Syn Bio* (2016) |
| PmHMGR | 3a | Pm | *Pseudomonas mevalonii* | P13702 | 312_pJL1-(CAT5aa)-HMGR_Sce | MEKKI | 46091 | 30 °C | Ma *et al. Met Eng* (2011), Dudley *et al. ACS Syn Bio* (2016) |
| ScHMGR | 3b | Sc | *Saccharomyces cerevisiae* | P12683 | 313_pJL1-(CAT5aa)-HMGR_Sau | MEKKI | 53632 | 30 °C | Dueber *et al. Nat Biotech* (2009), Dudley *et al. ACS Syn Bio* (2016), many others |
| SaHMGR | 3c | Sa | *Staphylococcus aureus* | A0A0Y9XRG8 | 314_pJL1-(CAT5aa)-HMGR_Pme | MEKKI | 46814 | 30 °C | Tsuruta *et al. PloS One* (2009), Dudley *et al. ACS Syn Bio* (2016) |
| SpHMGR | 3d | Sp | *Streptococcus pneumoniae* | M5KNC3 | 315_pJL1-(CAT5aa)-HMGR_Spn | MEKKI | 46752 | 30 °C | Yoon *et al. J Biotech* (2009), Dudley *et al. ACS Syn Bio* (2016) |
| BpHMGR | 3e | Bp | *Bordetella petrii* | A9HWZ9 | 316_pJL1-(CAT5aa)-HMGR_Bpe | MEKKI | 45178 | 30 °C | Ma *et al. Met Eng* (2011), Dudley *et al. ACS Syn Bio* (2016) |
| DaHMGR | 3f | Da | *Delftia acidovorans* | S2WYR6 | 317_pJL1-(CAT5aa)-HMGR_Dac | MEKKI | 45653 | 30 °C | Ma *et al. Met Eng* (2011), Dudley *et al. ACS Syn Bio* (2016) |
| ScMK | 4a | Sc | *Saccharomyces cerevisiae* | P07277 | 281_pJL1-(CAT5aa)-MK1_Sce | MEKKI | 48959 | 30 °C | Dueber *et al. Nat Biotech* (2009), Dudley *et al. ACS Syn Bio* (2016), many others |
| SaMK | 4b | Sa | *Staphylococcus aureus* | W8TNT9 | 321_pJL1-(CAT7aa)-MK_Sau | MHMEKKI | 33687 | 30 °C | Voynova *et al. J Bacteriol* (2004), many others * |
| SpMK | 4c | Sp | *Streptococcus pneumoniae* | A0A0E8Z6G1 | 322_pJL1-(CAT7aa)-MK_Spn | MHMEKKI | 32238 | 30 °C | Zurbriggen *et al. BioEnergy Res* (2012), Yang *et al. Microbial Cell Factories* (2016) |
| MmMK | 4d | Mm | *Methanosarcina mazei* | Q8PW39 | 323_pJL1-(CAT7aa)-MK_Mma | MHMEKKI | 32180 | 30 °C | Primak *et al. Appl Enviro Microbiol* (2011) |
| PzMK | 4e | Pz | *Paracoccus zeaxanthinifaciens* | Q8L1I2 | 324_pJL1-(CAT7aa)-MK_Pze | MHMEKKI | 40067 | 16 °C | Bai *et al.* US7422884 (2008), Berry *et al.* US20090226986 (2009) |
| HmMK | 4f | Hm | *Haloferax mediterranei* | I3R889 | 325_pJL1-(CAT7aa)-MK_Hme | MHMEKKI | 37656 | 30 °C | sequence not previously characterized |
| NmMK | 4g | Nm | *Nitrosopumilus maritimus* | A9A427 | 326_pJL1-(CAT7aa)-MK_Nma | MHMEKKI | 35199 | 30 °C | sequence not previously characterized |
| MxMK | 4h | Mx | *Myxococcus xanthus* | Q1D2E9 | 327_pJL1-(CAT7aa)-MK_Mxa | MHMEKKI | 32719 | 30 °C | sequence not previously characterized |
| BmMK | 4i | Bm | *Bacopa monniera* | M1F2U2 | 328_pJL1-(CAT7aa)-MK_Bmo | MHMEKKI | 41697 | 16 °C | Kumari *et al. Intl J Biol Macromol* (2015) |
| ScPMK | 5a | Sc | *Saccharomyces cerevisiae* | P32377 | 282_pJL1-(CAT5aa)-PMK1_Sce | MEKKI | 50955 | 30 °C | Dueber *et al. Nat Biotech* (2009), Dudley *et al. ACS Syn Bio* (2016), many others |
| SaPMK | 5b | Sa | *Staphylococcus aureus* | A0A0K6VEI6 | 329_pJL1-(CAT7aa)-PMK_Sau | MHMEKKI | 40985 | 16 °C | Yang *et al. Microbial Cell Factories* (2016) |
| SpPMK | 5c | Sp | *Streptococcus pneumoniae* | Q8DR49 | 330_pJL1-(CAT7aa)-PMK_Spn | MHMEKKI | 37794 | 30 °C | Zurbriggen *et al. BioEnergy Res* (2012), Rodriguez and Leyh *PLoS One* (2014) |
| EfPMK | 5d | Ef | *Enterococcus faecalis* | A0A059MZ18 | 331_pJL1-(CAT7aa)-PMK_Efa | MHMEKKI | 41279 | 30 °C | Doun *et al. Protein Sci* (2005) |
| PzPMK | 5e | Pz | *Paracoccus zeaxanthinifaciens* | Q8L1I1 | 332_pJL1-(CAT7aa)-PMK_Pze | MHMEKKI | 32352 | 30 °C | sequence not individually characterized, see Berry *et al.* US20090226986 (2009) |
| ThPMK | 5f | Th | *Tetragenococcus halophilus* | G4L4J3 | 333_pJL1-(CAT7aa)-PMK_Tha | MHMEKKI | 40891 | 30 °C | sequence not previously characterized |
| EcPMK | 5g | Ec | *Eremococcus coleocola* | E4KRA4 | 334_pJL1-(CAT7aa)-PMK_Eco | MHMEKKI | 41564 | 16 °C | sequence not previously characterized |
| ScPMD | 6a | Sc | *Saccharomyces cerevisiae* | P32377 | 283_pJL1-(CAT5aa)-PMD1_Sce | MEKKI | 44616 | 30 °C | Dueber *et al. Nat Biotech* (2009), Dudley *et al. ACS Syn Bio* (2016), many others |
| SaPMD | 6b | Sa | *Staphylococcus aureus* | X5E5B5 | 335_pJL1-(CAT7aa)-PMD_Sau | MHMEKKI | 37591 | 30 °C | Yang *et al. Microbial Cell Factories* (2016) |
| SpPMD | 6c | Sp | *Streptococcus pneumoniae* | D6ZQL9 | 336_pJL1-(CAT7aa)-PMD_Spn | MHMEKKI | 36315 | 30 °C | Zurbriggen *et al. BioEnergy Res* (2012), Yang *et al. Microbial Cell Factories* (2016) |
| PkPMD | 6d | Pk | *Picrorhiza kurrooa* | S5Y9Q9 | 337_pJL1-(CAT7aa)-PMD_Pku | MHMEKKI | 47000 | 16 °C | sequence not previously characterized |
| PzPMD | 6e | Pz | *Paracoccus zeaxanthinifaciens* | Q8L1I0 | 338_pJL1-(CAT7aa)-PMD_Pze | MHMEKKI | 36190 | 30 °C | sequence not individually characterized, see Berry *et al.* US20090226986 (2009) |
| HmPMD | 6f | Hm | *Haloferax mediterranei* | I3R4N4 | 339_pJL1-(CAT7aa)-PMD_Hme | MHMEKKI | 36432 | 16 °C | sequence not previously characterized |
| ZgPMD | 6g | Zg | *Zobellia galactanivorans* | G0LAG7 | 340_pJL1-(CAT7aa)-PMD_Zga | MHMEKKI | 41366 | 16 °C | sequence not previously characterized |

* *Sa*MK (Lefurgy et al., 2010; Rodriguez and Leyh, 2014; Voynova et al., 2004; Weaver et al., 2014)

**Table S1.** *continued*

| Enzyme | # | Abbr. | Source organism | uniprot ID | Plasmid name | N-terminal tag | Molecular weight (Da) | CFPS expression temperature | Reference |
|---|---|---|---|---|---|---|---|---|---|
| EcIDI | 7a | Ec | *Escherichia coli* (Type I) | Q46822 | 284_pJL1-(CAT5aa)-IDI_Eco | MEKKI | 21008 | 30 °C | Dueber *et al. Nat Biotech* (2009), Dudley *et al. ACS Syn Bio* (2016), many others |
| BsIDI | 7b | Bs | *Bacillus subtilis* (Type II) | P50740 | 285_pJL1-(CAT5aa)-IDI_Bsu | MEKKI | 37720 | 30 °C | Wang *et al. Process Biochem* (2015) |
| SceIDI | 7c | Sc | *Saccharomyces cerevisiae* (Type I) | P15496 | 341_pJL1-(CAT7aa)-IDI_Sce | MHMEKKI | 34119 | 30 °C | Anderson *et al. J Biol Chem* (1989) |
| SlIDI | 7d | Sl | *Solanum lycopersicym (tomato)* (Type I) | A9LRT7 | 342_pJL1-(CAT7aa)-IDI_Sly | MHMEKKI | 27954 | 30 °C | Berthelot *et al. Biochimie* (2016) |
| StIDI | 7e | St | *Streptomyces sp strain CL190* (Type II) | Q9KWG2 | 343_pJL1-(CAT7aa)-IDI_Str | MHMEKKI | 39610 | 30 °C | Kaneda *et al. Proceedings of the National Academy of Sciences* (2001) |
| PzIDI | 7f | Pz | *Paracoccus zeaxanthinifaciens* (Type II) | Q8L1I4 | 344_pJL1-(CAT7aa)-IDI_Pze | MHMEKKI | 38072 | 30 °C | sequence not individually characterized, see Berry *et al.* US20090226986 (2009) |
| SaIDI | 7g | Sa | *Staphylococcus aureus* (Type II) | P58052 | 345_pJL1-(CAT7aa)-IDI_Sau | MHMEKKI | 39568 | 30 °C | Kao *et al. Organic Letters* (2005) |
| SclIDI | 7h | Sc | *Streptomyces clavuligerus* (Type I) | E2PYC0 | 346_pJL1-(CAT7aa)-IDI_Scl | MHMEKKI | 23009 | 30 °C | sequence not previously characterized |
| PaGPPS | 8a | Pa | *Picea abies* (Norway spruce) | B1A9K6 | 286_pJL1-(CAT5aa)-GPPS_Agr_F3F | MEKKI | 33025 | 30 °C | Schmidt and Gershenzon *Phytochem* (2008), Sarria *et al. ACS Syn Bio* (2014) |
| AgGPPS | 8b | Ag | *Abies grandis* (grand fir) | Q8LKJ2 | 287_pJL1-(CAT5aa)-GPPS2_Pab | MEKKI | 32919 | 16 °C | Sarria *et al. ACS Syn Bio* (2014) and many others** |
| StGPPS | 8c | St | *Streptomyces sp. strain KO-3988* | Q2L6D1 | 288_pJL1-(CAT5aa)-GPPS_Str | MEKKI | 38580 | 16 °C | Willrodt *et al. Biotech J* (2014) |
| PgGPPS | 8d | Pg | *Picea glauca* (white spruce) | V9Y2D8 | 347_pJL1-(CAT7aa)-GPPS_Pgl | MHMEKKI | 33002 | 30 °C | sequence not previously characterized |
| PkGPPS | 8e | Pk | *Picrorhiza kurrooa* | Q5G1J1 | 348_pJL1-(CAT7aa)-GPPS_Pku | MHMEKKI | 41160 | 16 °C | Singh *et al. Gene* (2013) |
| Sc*GPPS | 8f | Sc* | *Saccharomyces cerevisiae* (K197E) | P08524 | 349_pJL1-(CAT7aa)-GPPS*_Sce | MHMEKKI | 41252 | 30 °C | Fischer *et al. Biotech Bioeng* (2011) |
| EcFPPS | 8g | Ec | *Escherichia coli* | P22939 | 318_pJL1-(CAT5aa)-ispA_Ec | MEKKI | 32659 | 30 °C | Dudley *et al. Syn Bio* (2019) |
| MsLS | 9a | Ms | *Mentha spicata* (spearmint) | Q40322 | 246_pJL1-(CAT5aa)-LS_Msp | MEKKI | 64317 | 30 °C | Dudley *et al. Syn Bio* (2019) and many others*** |
| ClLS | 9b | Cl | *Citrus limon* (lemon) | Q8L5K3 | 247_pJL1-(CAT5aa)-LS_Cli | MEKKI | 65350 | 30 °C | Lücker *et al. Eur J Biochem* (2002), Jongedijk *et al. Yeast* (2015) |
| PfLS | 9c | Pf | *Perilla frutescens* (wild sesame) | Q9AXM7 | 248_pJL1-(CAT5aa)-LS_Pfr | MEKKI | 64449 | 16 °C | Yuba *et al. Arch Biochem Biophys* (1996), Jongedijk *et al. Yeast* (2015) |
| StLS | 9d | St | *Schizonepeta tenufolio* (Jap. catnip) | Q9FUW5 | 350_pJL1-(CAT7aa)-LS_Ste | MHMEKKI | 65095 | 30 °C | Kiyota *et al. J Biotechnol* (2014) |
| LaLS | 9e | La | *Lavandula angustifolia* (lavender) | Q2XSC6 | 351_pJL1-(CAT7aa)-LS_Lan | MHMEKKI | 64819 | 16 °C | Landmann *et al. Arch Biochem Biophys* (2007) |
| SlLS | 9f | Sl | *Solanum lycopersicum* (tomato) | G1JUH1 | 352_pJL1-(CAT7aa)-LS_Sly | MHMEKKI | 65586 | 16 °C | Falara *et al. Plant Physiol* (2011) |
| PaLS | 9g | Pa | *Picea abies* (Norway spruce) | Q675L1 | 353_pJL1-(CAT7aa)-LS-Pab | MHMEKKI | 66801 | 16 °C | sequence not previously characterized |
| AgBS | | Ag | *Abies grandis* (grand fir) | O81086 | 320_pJL1-(CAT5aa)-BS_Agr | MEKKI | 94338 | 16 °C | Peralta-Yahya *et al. Nat Comm* (2011) |
| AgPS | | Ag | *Abies grandis* (grand fir) | O24475 | 249_11a_pJL1_CATrbs(5aa)_PS_Agr | MEKKI | 68104 | 30 °C | Sarria *et al. ACS Syn Bio* (2014) |
| PaPS | | Pa | *Picea abies* (Norway spruce) | Q675L3 | 250_11b_pJL1_CATrbs(5aa)_PS_Pab | MEKKI | 65454 | 30 °C | Sarria *et al. ACS Syn Bio* (2014) |

** *Ag*GPPS (Alonso-Gutierrez et al., 2013; Burke and Croteau, 2002; Carter et al., 2003; Kang et al., 2014; Willrodt et al., 2014; Zhao et al., 2016)

*** *Ms*LS, see (Jongedijk et al., 2016) for list of papers using this enzyme

References (Anderson et al., 1989; Berry et al., 2009; Berthelot et al., 2016; Doun et al., 2005; Dueber et al., 2009; Falara et al., 2011; Fischer et al., 2011; Jongedijk et al., 2015; Kaneda et al., 2001; Kao et al., 2005; Kiyota et al., 2014; Kumari et al., 2015; Landmann et al., 2007; Lücker et al., 2002; Schmidt and Gershenzon, 2008; Singh et al., 2013; Tsuruta et al., 2009; Wang et al., 2015; Willrodt et al., 2014; Yang et al., 2016; Yoon et al., 2009; Yuba et al., 1996; Zurbriggen et al., 2012)

**Table S2. Estimation of overexpresesd protein concentration in pre-enriched lysates.** To estimate the concentration of overexpressed proteins in pre-enriched (*i.e.,* CFME) lysates, we used densitometry to estimate the band percentage in a Coomassie stained SDS-PAGE gel using ImageLab software. These percentages are multiplied by the concentration of pre-enriched lysates and converted to a molar concentration using the molecular weight of the protein in question.

| **Protein enriched in the lysate** | **Band %** | **overexpressed protein molecular weight (Da)** | **Amount of crude lysate typically loaded into cell-free reaction (mg/mL total protein)** | **µM overexpressed protein in cell-free reaction *** | **Reference for original SDS-PAGE gel** | |
|---|---|---|---|---|---|---|
| ACAT_Eco (i.e. EcACAT) | 14.2% | 40607 | 1.33 | 4.6 | Dudley *et al. ACS Syn Bio* (2016) Fig. S | Lane 2 |
| HMGS_Sce (i.e. ScHMGS) | 22.9% | 55756 | 1.33 | 5.5 | Dudley *et al. ACS Syn Bio* (2016) Fig. S | Lane 3 |
| HMGS_Sau (i.e. SaHMGS) | 12.6% | 43706 | 1.33 | 3.8 | Dudley *et al. ACS Syn Bio* (2016) Fig. S | Lane 5 |
| HMGR_Pme (i.e. PmHMGR) | 15.4% | 46091 | 1.33 | 4.5 | Dudley *et al. ACS Syn Bio* (2016) Fig. S | Lane 8 |
| ACAT_Eco (i.e. EcACAT) | 8.1% | 40607 | 4 | 8.0 | Dudley *et al. Syn Bio* (2019) Fig. S4 | Lane 3 |
| HMGS_Sce (i.e. ScHMGS) | 8.4% | 55756 | 4 | 6.0 | Dudley *et al. Syn Bio* (2019) Fig. S4 | Lane 3 |
| HMGR_Pme (i.e. PmHMGR) | 6.8% | 46091 | 4 | 5.9 | Dudley *et al. Syn Bio* (2019) Fig. S4 | Lane 3 |
| MK_Sce (i.e. ScMK) | 10.1% | 48959 | 1 | 2.1 | Dudley *et al. Syn Bio* (2019) Fig. S4 | Lane 4 |
| PMK_Sce (i.e. ScPMK) | 16.1% | 50955 | 1 | 3.2 | Dudley *et al. Syn Bio* (2019) Fig. S4 | Lane 5 |
| PMD_Sce (i.e. ScPMK) | 3.7% | 44616 | 1 | 0.8 | Dudley *et al. Syn Bio* (2019) Fig. S4 | Lane 6 |
| IDI_Eco (i.e. EcIDI) | 18.2% | 21008 | 1 | 8.7 | Dudley *et al. Syn Bio* (2019) Fig. S4 | Lane 7 |
| LS_Msp (i.e. MsLS) | 16.3% | 64317 | 1 | 2.5 | Dudley *et al. Syn Bio* (2019) Fig. S4 | Lane 9 |
| LS_Msp (i.e. MsLS) | 16.3% | 64317 | 1.33 | 3.4 | Dudley *et al. Syn Bio* (2019) Fig. S4 | Lane 9 |

** = % overexpressed band of total protein * crude lysate loaded / overexpressed protein molecular weight*

**Table S3.** Details of enzyme and cofactor concentrations used in each experiment.

| Enzyme Set | Figure | cofactors (mM) | | | pre-enriched lysates | | | | CFPS reactions | | | | | | | | | | | | | |
|---|---|---|---|---|---|---|---|---|---|---|---|---|---|---|---|---|---|---|---|---|---|---|
| | | NAD+ | CoA | ATP | EcACAT | ScHMGS | MsLS | "blank" | HMGR | µM | MK | µM | PMK | µM | PMD | µM | IDI | µM | GPPS | µM | synthase | µM |
| Set 1.0 | S5 | 0 | 0 | 0 | - | - | - | + | PmHMGR | 0.4 | ScMK | 0.4 | ScPMK | 0.4 | ScPMD | 0.4 | EcIDI | 1.0 | PaGPPS | 0.6 | MsLS | 1.6 |
| - | 2C-E†† | 0 | 0 | 0 | + | + | ** | ** | * PmHMGR | 1.0†† | * ScMK | 0.4 | * ScPMK | 0.4 | * ScPMD | 0.4 | * EcIDI | 2.0†† | * PaGPPS | 3.0†† | * MsLS | ** |
| Set 1.1 | 3A | 0 | 0 | 0 | + | + | + | - | PmHMGR | 1.0 | ScMK | 0.4 | ScPMK | 0.4 | ScPMD | 0.4 | EcIDI | 2.0 | PaGPPS | 3.0 | | |
| Set 2.0 | 3A | 0 | 0 | 0 | + | + | + | - | BpHMGR | 1.0 | MmMK | 0.4 | SpPMK | 0.4 | ScPMD | 0.4 | SclIDI | 2.0 | PaGPPS | 3.0 | | |
| Set 1.1 | 3B | * | * | * | + | + | + | - | PmHMGR | 1.0 | ScMK | 0.4 | ScPMK | 0.4 | ScPMD | 0.4 | EcIDI | 2.0 | PaGPPS | 3.0 | | |
| Set 2.0 | 3B | * | * | * | + | + | + | - | BpHMGR | 1.0 | MmMK | 0.4 | SpPMK | 0.4 | ScPMD | 0.4 | SclIDI | 2.0 | PaGPPS | 3.0 | | |
| Set 2.1 | S8 | 5 | 1 | 1 | + | + | + | - | BpHMGR | 0.2 | MmMK | 0.1 | SpPMK | 0.2 | ScPMD | 0.2 | SclIDI | 0.2 | PaGPPS | 3.0 | | |
| Set 2.2 | 4 | Four cofactor conditions: α = no supple-mented cofactors; β = 5 mM NAD+ 1 mM CoA 0 mM ATP; γ = 5 mM NAD+ 0 mM CoA 1 mM ATP; δ = 5 mM NAD+ 1 mM CoA 1 mM ATP | | | + | + | + | - | BpHMGR | 0.2 | MmMK | 0.1 | SpPMK | 0.2 | ScPMD | 0.2 | SclIDI | 0.2 | * PaGPPS | 1.0 | | |
| Set 3.1 | 4 | | | | + | + | + | - | BpHMGR | 0.2 | MmMK | 0.1 | SpPMK | 0.2 | ScPMD | 0.2 | * SclIDI | 0.2 | PgGPPS | 1.0 | | |
| Set 3.2 | 4 | | | | + | + | + | - | BpHMGR | 0.2 | MmMK | 0.1 | SpPMK | 0.2 | ScPMD | 0.2 | * SclIDI | 0.2 | StGPPS | 1.0 | | |
| Set 4.1 | 4 | | | | + | + | + | - | BpHMGR | 0.2 | MmMK | 0.1 | SpPMK | 0.2 | * ScPMD | 0.2 | SaIDI | 0.2 | PgGPPS | 1.0 | | |
| Set 4.2 | 4 | | | | + | + | + | - | BpHMGR | 0.2 | MmMK | 0.1 | SpPMK | 0.2 | * ScPMD | 0.2 | SlIDI | 0.2 | PgGPPS | 1.0 | | |
| Set 4.3 | 4 | | | | + | + | + | - | BpHMGR | 0.2 | MmMK | 0.1 | SpPMK | 0.2 | * ScPMD | 0.2 | SceIDI | 0.2 | StGPPS | 1.0 | | |
| Set 4.4 | 4 | | | | + | + | + | - | BpHMGR | 0.2 | MmMK | 0.1 | SpPMK | 0.2 | * ScPMD | 0.2 | BsIDI | 0.2 | StGPPS | 1.0 | | |
| Set 5.1 | 4 | | | | + | + | + | - | BpHMGR | 0.2 | MmMK | 0.1 | * SpPMK | 0.2 | ScPMD | 0.2 | SaIDI | 0.2 | PgGPPS | 1.0 | | |
| Set 5.2 | 4 | | | | + | + | + | - | BpHMGR | 0.2 | MmMK | 0.1 | * SpPMK | 0.2 | ScPMD | 0.2 | SlIDI | 0.2 | PgGPPS | 1.0 | | |
| Set 5.3 | 4 | | | | + | + | + | - | BpHMGR | 0.2 | MmMK | 0.1 | * SpPMK | 0.2 | ScPMD | 0.2 | SceIDI | 0.2 | StGPPS | 1.0 | | |
| Set 5.4 | 4 | | | | + | + | + | - | BpHMGR | 0.2 | MmMK | 0.1 | * SpPMK | 0.2 | SaPMD | 0.2 | SlIDI | 0.2 | PgGPPS | 1.0 | | |
| Set 5.5 | 4 | | | | + | + | + | - | BpHMGR | 0.2 | MmMK | 0.1 | * SpPMK | 0.2 | ZgPMD | 0.2 | SlIDI | 0.2 | PgGPPS | 1.0 | | |
| Set 6.1 | 4 | | | | + | + | + | - | BpHMGR | 0.2 | * MmMK | 0.1 | PzPMK | 0.2 | ScPMD | 0.2 | SlIDI | 0.2 | PgGPPS | 1.0 | | |
| Set 6.2 | 4 | | | | + | + | + | - | BpHMGR | 0.2 | * MmMK | 0.1 | PzPMK | 0.2 | ScPMD | 0.2 | SaIDI | 0.2 | PgGPPS | 1.0 | | |
| Set 6.3 | 4 | | | | + | + | + | - | BpHMGR | 0.2 | * MmMK | 0.1 | ScPMK | 0.2 | ScPMD | 0.2 | SlIDI | 0.2 | PgGPPS | 1.0 | | |
| Set 6.4 | 4 | | | | + | + | + | - | BpHMGR | 0.2 | * MmMK | 0.1 | EcPMK | 0.2 | ScPMD | 0.2 | SlIDI | 0.2 | PgGPPS | 1.0 | | |
| Set 6.5 | 4 | | | | + | + | + | - | BpHMGR | 0.2 | * MmMK | 0.1 | ScPMK | 0.2 | ScPMD | 0.2 | SceIDI | 0.2 | StGPPS | 1.0 | | |
| Set 7.1 | 4 | | | | + | + | + | - | * BpHMGR | 0.2 | MmMK | 0.1 | PzPMK | 0.2 | ScPMD | 0.2 | SlIDI | 0.2 | PgGPPS | 1.0 | | |
| Set 7.2 | 4 | | | | + | + | + | - | * BpHMGR | 0.2 | MmMK | 0.1 | PzPMK | 0.2 | ScPMD | 0.2 | SaIDI | 0.2 | PgGPPS | 1.0 | | |
| Set 7.3 | 4 | | | | + | + | + | - | * BpHMGR | 0.2 | MmMK | 0.1 | ScPMK | 0.2 | ScPMD | 0.2 | SlIDI | 0.2 | PgGPPS | 1.0 | | |
| Set 7.4 | 4 | | | | + | + | + | - | * BpHMGR | 0.2 | MmMK | 0.1 | EcPMK | 0.2 | ScPMD | 0.2 | SlIDI | 0.2 | PgGPPS | 1.0 | | |
| Set 7.5 | 4 | | | | + | + | + | - | * BpHMGR | 0.2 | MmMK | 0.1 | ScPMK | 0.2 | ScPMD | 0.2 | SceIDI | 0.2 | StGPPS | 1.0 | | |
| Set 8.1 | 4 | | | | + | § | + | - | BpHMGR | 0.2 | MmMK | 0.1 | PzPMK | 0.2 | ScPMD | 0.2 | SlIDI | 0.2 | PgGPPS | 1.0 | | |
| Set 8.2 | 4 | | | | + | § | + | - | BpHMGR | 0.2 | MmMK | 0.1 | ScPMK | 0.2 | ScPMD | 0.2 | SlIDI | 0.2 | PgGPPS | 1.0 | | |
| Set 8.3 | 4 | | | | + | § | + | - | BpHMGR | 0.2 | MmMK | 0.1 | EcPMK | 0.2 | ScPMD | 0.2 | SlIDI | 0.2 | PgGPPS | 1.0 | | |
| Set 9.0 | - | 5 | 1 | 1 | + | + | + | - | BpHMGR | 0.2 | MmMK | 0.1 | PzPMK | 0.2 | ScPMD | 0.2 | SlIDI | 0.2 | PgGPPS | 1.0 | | |
| Set 10.0 | 5 | 5 | 1 | 1 | + | + | - | + | BpHMGR | 0.2 | MmMK | 0.1 | PzPMK | 0.2 | ScPMD | 0.2 | SlIDI | 0.2 | PgGPPS | 1.0 | MsLS | 3.8 |
| Set 10.0_AgPS | 5 | 5 | 1 | 1 | + | + | - | + | BpHMGR | 0.2 | MmMK | 0.1 | PzPMK | 0.2 | ScPMD | 0.2 | SlIDI | 0.2 | PgGPPS | 1.0 | AgPS | 3.8 |
| Set 10.0_PaPS | 5 | 5 | 1 | 1 | + | + | - | + | BpHMGR | 0.2 | MmMK | 0.1 | PzPMK | 0.2 | ScPMD | 0.2 | SlIDI | 0.2 | PgGPPS | 1.0 | PaPS | 3.8 |
| Set 10.1_AgBS | 5 | 5 | 1 | 1 | + | + | - | + | BpHMGR | 0.2 | MmMK | 0.1 | PzPMK | 0.2 | ScPMD | 0.2 | SlIDI | 0.2 | EcFPPS | 1.0 | AgBS | 1.6 |

+ indicates a pre-enriched lysate

* variable paramter, listed homolog is the default homolog used as reference or carried in from prior iteration

** MsLS is typically present as a pre-enriched lysate. When testing LS homologs, LS was added as a CFPS reaction (1.0 µM) and "blank" lysate without a plamid was used in place of the MsLS-enriched lysate

†† Instead of 1.0 µM, ScHMGR is tested at 0.5 µM.

†† Instead of 2.0 µM, select IDI homologs are included at 1.1 µM (BsIDI), 0.5 µM (StIDI), 1.2 (PzIDI), and 1.5 µM (SaIDI).

†† Instead of 3.0 µM, select GPPS homologs are included at 1.3 µM (AgGPPS), 1.0 µM (StGPPS), and 0.7 µM (PkGPPS)

§ HMGS homologs tested as both pre-enriched lysate (SaHMGS ~ 3.8 µM, ScHMGS ~5.5 µM) and as CFPS reactions (1.5 µM)

Note that some enzyme sets have multiple names (for example sets 6.1, 7.1, and 9.0 are identical)

**Table S4.** Metabolic engineering for the production of monoterpenes.

| Monoterpenoid product | Host | Pathway | Synthase enzyme source organism | Maximum titer (mg/L) | Volumetric productivity (mg/L/day) | Reference |
|---|---|---|---|---|---|---|
| 3-Carene | *E. coli* | MEP/DXP | *Picea abies* | 0.003 | 0.01 | Reiling *et al. Biotechnol Bioeng* (2004) |
| Linalool | *S. cerevisiae* | MVA | *Lavandula angustifolia* | 0.1 | n/a | Amiri *et al. Biotechnol Letters* (2016) |
| α/β-Pinene | *C. glutamicum* | MEP/DXP | *Abies grandis* | 0.2 | 0.1 | Kang *et al. Biotechnol Letters* (2014) |
| Limonene | *Synechocystis* | MEP/DXP | *Schizonepeta tenuifolia* | 0.4 | 0.1 | Kiyota *et al. J Biotechnol* (2014) |
| Limonene | *E. coli* | MVA, glucose | *Mentha spicata* | 0.3 | 0.2 | Willrodt *et al. Biotechnol J* (2014) |
| Limonene | *S. cerevisiae* | MVA | *Citrus limon, Perilla frutescens* | 0.5 | 0.2 | Jongedijk *et al. Yeast* (2015) |
| Sabinene | *S. cerevisiae* | MVA | *Slavia pomifera* | 17.5 | n/a | Ignea *et al. ACS Syn Biol* (2014) |
| Limonene | *S. cerevisiae* | MVA | *Mentha spicata, Citrus limon* | 1.5 | 0.3 | Behrendorff *et al. Microb Cell Fact* (2011) |
| Limonene | *S. cerevisiae* | MVA | *Citrus limon* (H570Y) | 130 | 0.9 | Ignea *et al. Nat Comm* (2019) |
| Limonene | *Synechococcus* | MEP/DXP | *Mentha spicata* | 4.0 | 1.3 | Davies *et al. Front Bioeng Biotechnol* (2014) |
| Limonene | *E. coli* | MVA | *Mentha spicata* | 57.0 | 2.4 | Dunlop *et al. Mol Syst Biol* ( 2011) |
| Limonene | *E. coli* | MEP/DXP | *Mentha spicata* | 5.0 | 4.0 | Carter *et al. Phytochem* (2003) |
| α/β-Pinene | *E. coli* | MVA | *Abies grandis* | 32.4 | 10.8 | Sarria *et al. ACS Syn Bio* (2014) |
| Limonene | *E. coli* | MEP/DXP | *Mentha spicata* | 35.8 | 11.9 | Du *et al. Biores Bioproc* (2014) |
| Geraniol | *E. coli* | MVA | *Camptotheca acuminata* | 48 | - | Chen *et al. J Ind Microbiol Biotechnol* (2016) |
| Myrcene | *E. coli* | MVA | *Quercus ilex* | 58.2 | 19.4 | Kim *et al. J Agric Food Chem* (2015) |
| Cineole | *S. cerevisiae* | MVA | *Salvia fruticosa* | 1100 | 57.9 | Ignea *et al. Microb Cell Fact* (2011) |
| Limonene | *E. coli* | MVA | *Mentha spicata* | 230 † | 77 | Jervis *et al. ACS Syn Biol* (2019) |
| Sabinene | *E. coli* | MVA | *Slavia pomifera* | 82.2 | 82.2 | Zhang *et al. Microb Cell Fact* (2014) |
| Limonene | Cell-free* | MVA | *Mentha spicata* | 90.2 | 90.2 | Dudley *et al. Syn Bio* (2019) |
| Geraniol | *E. coli* | MVA | *Ocimum basilicum* | 183 | 91.3 | Zhou *et al. J Biotechnol* (2014) |
| Geraniol | *S. cerevisiae* | MVA | *Valeriana officinalis* | 293 | 97.7 | Zhao *et al. Appl Microbiol Biotechnol* (2016) |
| Limonene | *S. cerevisiae* | MVA | *Citrus limon* | 917 | 183 | Cheng *et al. ACS Syn Biol* (2019) |
| Limonene | *E. coli* | MVA | *Mentha spicata* | 604 | 202 | Alonso-Gutierrez *et al. Met Eng* (2015) |
| Limonene | *E. coli* | MVA | *Citrus limon* | 1290 | 369 | Wu *et al. J Agric Food Chem* (2019) |
| **Limonene** | **Cell-free**** | **MVA** | ***Mentha spicata*** | **610** | **610** | **This work** |
| Geraniol | *E. coli* | MVA | *Ocimum basilicum* | 2000 | 706 | Liu *et al. Biotechnol Biofuels* (2016) |
| Limonene | *E. coli* | MVA, glycerol | *Mentha spicata* | 1350 | 736 | Willrodt *et al. Biotech J* (2014) |
| Limonene | Cell-free*** | MVA | *Mentha spicata* | 12500 | 1786 | Korman *et al. Nat Comm* (2017) |

† 1150 mg/L organic phase * 20% overlay

* CFME approach using pre-enriched *E. coli* lysates

** CFPS-ME approaching combining pre-enriched *E. coli* lysates with CFPS-enriched lysates

*** purified enzymes

| Monoterpenoid product | Host | Pathway | Synthase enzyme source organism | Titer (µg/g fresh weight) | Reference |
|---|---|---|---|---|---|
| Linalool | *A. thaliana* | MVA + MEP/DXP | *Fragaria × ananassa* | 83 | Aharoni *et al. Plant Cell* (2003) |
| Geraniol | *N. benthamiana* | MVA + MEP/DXP | *Ocimum basilicum* | 110 | Fischer *et al. J Biotechnol* (2013) |
| Geraniol | *N. benthamiana* | MVA + MEP/DXP | *Valeriana officinalis* | 129 | Dong *et al. New Phytol* (2016) |

References (Aharoni et al., 2003; Alonso-Gutierrez et al., 2015; Amiri et al., 2016; Behrendorff et al., 2013; Carter et al., 2003; Chen et al., 2016; Cheng et al., 2019; Davies et al., 2014; Dong et al., 2016; Du et al., 2014; Dudley et al., 2019; Dunlop et al., 2011; Fischer et al., 2013; Ignea et al., 2011; Ignea et al., 2014; Jervis et al., 2019; Jongedijk et al., 2015; Kang et al., 2014; Kim et al., 2015; Kiyota et al., 2014; Korman et al., 2017; Liu et al., 2016; Reiling et al., 2004; Sarria et al., 2014; Willrodt et al., 2014; Wu et al., 2019; Zhang et al., 2014; Zhao et al., 2016; Zhou et al., 2014)

#### Supplementary Figures

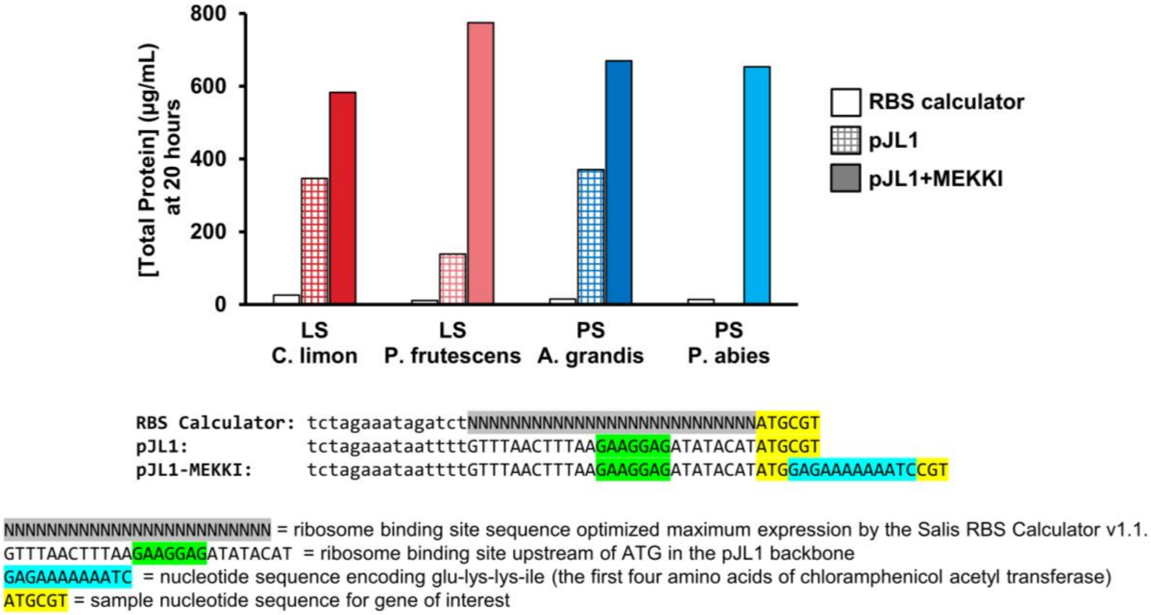


**Figure S1. Addition of N-terminal expression tag encoding MEKKI improves cell-free expression of monoterpene synthases.** Each limonene synthase (LS) or pinene synthase (PS) was expressed in CFPS using three different DNA plasmids. The “RBS calculator” version does not contain an N-terminal expression tag but used the Salis RBS Calculator v1.1 (Salis et al., 2009) to optimize the 27 base pair sequence (colored gray and containing a ribosome binding site) for maximum expression. The “pJL1” version does not contain a variable RBS or N-terminal expression tag but uses the RBS encoded by the pJL1 plasmid (Addgene #69496, pJL1 is derived from pY71 (Bundy and Swartz, 2010; Swartz et al., 2004)). The “pJL1-MEKKI” version places the coding sequence of the protein into pJL1 behind an N-terminal tag known to improve *in vitro* expression. The N-terminal tag is a 15 nucleotide, AT-rich sequence which encodes the first five amino acids (Met-Glu-Lys-Lys-Ile, MEKKI) of chloramphenicol acetyl transferase. Chloramphenicol acetyl transferase has been previously used as a reporter protein during the development of the *E. coli* CFPS system (Jewett and Swartz, 2004; Swartz et al., 2004).

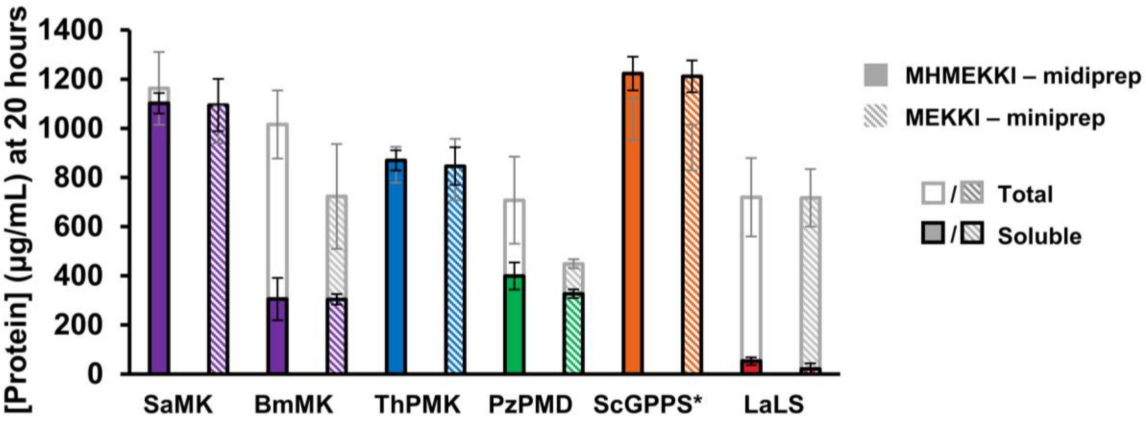


**Figure S2. Two variants of the N-terminal expression tag give similar expression for five different enzymes.** Gene sequences were unintentionally synthesized with two different N-terminal expression tags. The first tag sequence is catATGGAGAAAAAAATC (encoding MEKKI) and the second tag sequence is catATGCATATGGAGAAAAAAATC (encoding MHMEKKI). Comparison of the two N-terminal tags (using five different example enzymes) show similar soluble protein concentrations and both tags ultimately used (see **Table S1**). Values represent averages (n=3) and error bars represent 1 standard deviation.

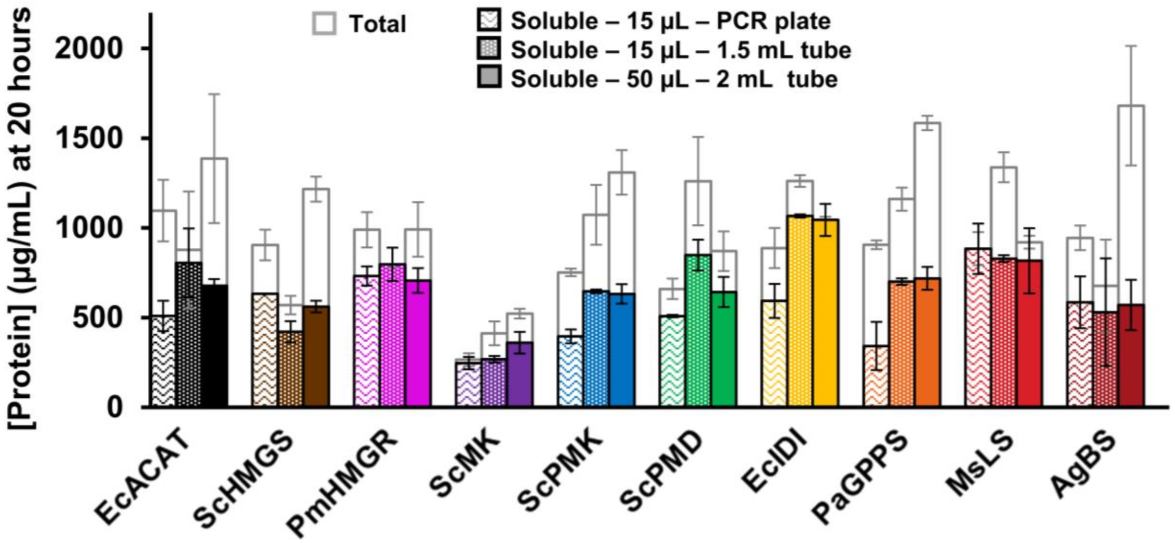


**Figure S3. Protein expression yields for ten representative proteins remain consistent when scaling up reaction volumes from 15 µL to 50 µL.** The standard 15 µL reaction is typically incubated in a 1.5 mL microcentrifuge tube. Due to the large amounts of CFPS reactions required for this manuscript, we wanted to scale up the reaction to minimize pipetting. To keep the reaction environment consistent at a larger reaction volume, 2 mL microcentrifuge tubes are used to incubate 50 µL CFPS reactions. Enzyme expression levels are similar between 15 µL and 50 µL reactions. Alternatively, we tried incubating 15 µL reactions in a PCR plate covered with foil lid (to minimize plastic/reduce pipetting) but found reduced the relative protein expression for some enzymes (*Pa*GPPS) compared to incubation in microcentrifuge tubes. Throughout the manuscript, all CFPS reactions were run at the 50 µL scale. Values represent averages (n=3) and error bars represent 1 standard deviation.

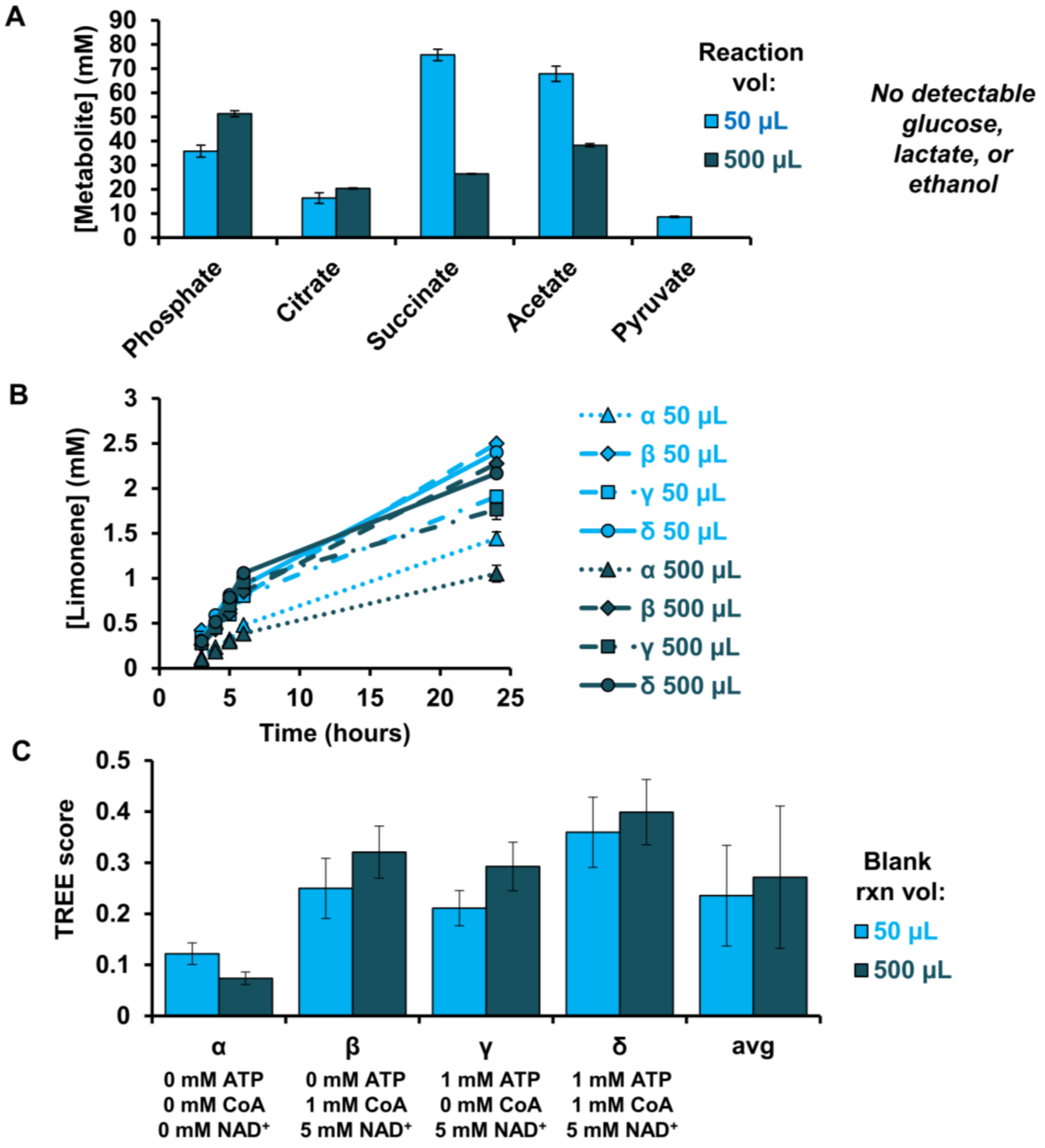


**Figure S4. Scaling volume of CFPS "blank" reactions (*i.e.,* CFPS without a plasmid substrate) from 50 to 500 µL changes the metabolic products.** (**A**) Analysis of metabolites in two CFPS reactions incubated for 20 hours with no DNA template. Values represent averages (n=3) and error bars represent 1 standard deviation. Phosphate is quantified by kit (Malachite Green Phosphate Assay POMG-25H, BioAssay Systems) while citrate, succinate, acetate, and pyruvate are measured on HPLC using methods previously described (Dudley et al., 2016). (**B-C**) Testing CFPS blank reaction volume using a glucose-to-limonene CFPS-ME reaction using enzyme set 3.1 with PgGPPS. Each 30 µL CFPS-ME reaction includes 11.2 µL "blank" CFPS reaction, 3.8 µL CFPS of pathway enzymes, 2.5 μL pre-enriched extracts, and 12.5 μL substrates/cofactors/water. Cofactor conditions α, β, γ, and δ are described in **Table S3**. (**B**) Limonene production where initial rate values are n=1 while the 24 hour value is an average (n=3) with error bars which represent 1 standard deviation. (**C**) Conversion of data from part B to a TREE score. Error bars represent propagated error (see **Appendix S1**) Due the metabolic differences between 50 μL and 500 μL "blank" reactions, all "blank" reactions used in this manuscript were made at the 50 μL scale for consistency.

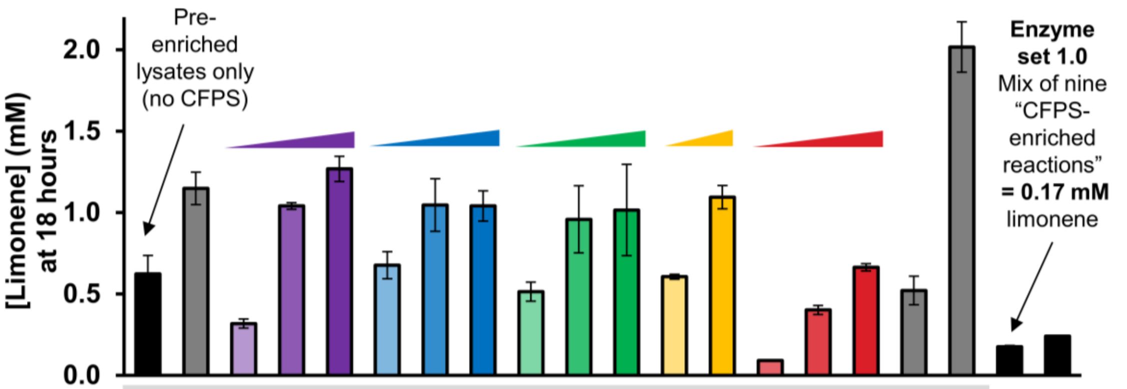


| | | | | | | | | | | | | | | | | | | | | | |
|---|---|---|---|---|---|---|---|---|---|---|---|---|---|---|---|---|---|---|---|---|---|
| *Ec*ACAT | 8.0* | 8.0* | 8.0* | 8.0* | 8.0* | 8.0* | 8.0* | 8.0* | 8.0* | 8.0* | 8.0* | 8.0* | 8.0* | 8.0* | 8.0* | 8.0* | 8.0* | 8.0* | 0.5 | 0.6 | μM |
| *Sc*HMGS | 6.0* | 6.0* | 6.0* | 6.0* | 6.0* | 6.0* | 6.0* | 6.0* | 6.0* | 6.0* | 6.0* | 6.0* | 6.0* | 6.0* | 6.0* | 6.0* | 6.0* | 6.0* | 0.4 | 0.6 | μM |
| *Pm*HMGR | 5.9* | 5.9* | 5.9* | 5.9* | 5.9* | 5.9* | 5.9* | 5.9* | 5.9* | 5.9* | 5.9* | 5.9* | 5.9* | 5.9* | 5.9* | 5.9* | 5.9* | 5.9* | 0.4 | 0.6 | μM |
| *Sc*MK | 2.1* | 2.1* | 0.2 | 0.8 | 1.8 | 2.1* | 2.1* | 2.1* | 2.1* | 2.1* | 2.1* | 2.1* | 2.1* | 2.1* | 2.1* | 2.1* | 0.4 | 0.4 | 0.4 | 0.4 | μM |
| *Sc*PMK | 3.2* | 3.2* | 3.2* | 3.2* | 3.2* | 0.2 | 2.0 | 3.9 | 3.2* | 3.2* | 3.2* | 3.2* | 3.2* | 3.2* | 3.2* | 3.2* | 0.4 | 0.4 | 0.4 | 0.4 | μM |
| *Sc*PMD | 0.8* | 0.8* | 0.8* | 0.8* | 0.8* | 0.8* | 0.8* | 0.8* | 0.2 | 2.2 | 4.5 | 0.8* | 0.8* | 0.8* | 0.8* | 0.8* | 0.4 | 0.4 | 0.4 | 0.4 | μM |
| *Ec*IDI | 8.7* | 8.7* | 8.7* | 8.7* | 8.7* | 8.7* | 8.7* | 8.7* | 8.7* | 8.7* | 8.7* | 0.5 | 4.8 | 8.7* | 8.7* | 8.7* | 1.0 | 1.0 | 1.0 | 2.0 | μM |
| *Pa*GPPS | - | 3.8 | 3.8 | 3.8 | 3.8 | 3.8 | 3.8 | 3.8 | 3.8 | 3.8 | 3.8 | 3.8 | 3.8 | 3.8 | 3.8 | 3.8 | 0.6 | 0.6 | 0.6 | 3.0 | μM |
| *Ms*LS | 2.5* | 2.5* | 2.5* | 2.5* | 2.5* | 2.5* | 2.5* | 2.5* | 2.5* | 2.5* | 2.5* | 2.5* | 2.5* | 0.2 | 1.0 | 2.2 | 2.2 | 2.5* | 1.6 | 2.5* | μM |

In the x-axis labeling table, a highlighted value with asterisk (**x.x***) indicates a pre-enriched lysate with the value indicates μM protein (estimated by densitometry). A non-highlighted value (**x.x**) indicates a CFPS reaction where the value indicates μM protein (measured via C14 leucine incorporation)

**Figure S5. Mixing of CFPS reactions with pre-enriched lysates indicates minimum enzyme concentrations needed for limonene production.** In order to compare the relative activity of pathway enzymes and determine appropriate concentrations for mixing CFPS-enriched reactions, we started with a mixture of 8 pre-enriched lysates capable of producing limonene (leftmost black bar, (Dudley et al., 2019)). We immediately observed that supplementing high concentrations of CFPS reaction expressing *Pa*GPPS increased limonene titer (leftmost gray bar) and used 3.8 μM in subsequent tests. We then removed the pre-enriched lysate encoding MK, PMK, PMD, IDI, and LS and added increasing concentrations of CFPS reaction expressing these reactions. These results informed the initial enzyme concentrations for Enzyme Set 1.0 used in **Figure 2** to be at 1.0 μM *Pm*HMGR, 0.4 μM ScMK, 0.4 μM ScPMK, 0.4 μM ScPMD, 2.0 EcIDI, 3.0 μM PaGPPS. Finally we show that the initial enzyme homologs, mixed as nine CFPS-enriched reactions, can support limonene production. Values represent averages (n=3) and error bars represent 1 standard deviation.

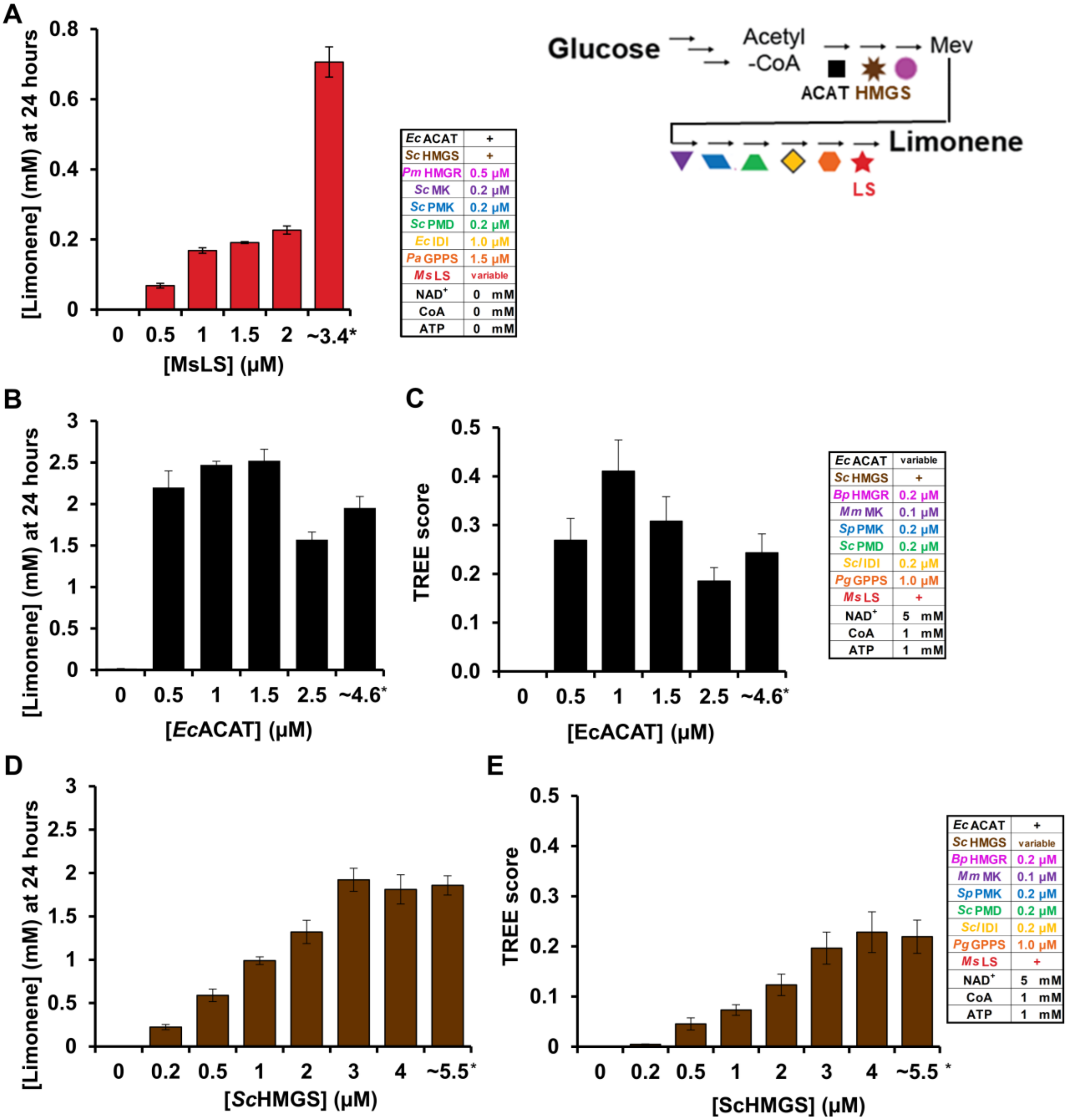


| | |
|---|---|
| *Ec* ACAT | + |
| *Sc* HMGS | + |
| *Pm* HMGR | 0.5 μM |
| *Sc* MK | 0.2 μM |
| *Sc* PMK | 0.2 μM |
| *Sc* PMD | 0.2 μM |
| *Ec* IDI | 1.0 μM |
| *Pa* GPPS | 1.5 μM |
| *Ms* LS | variable |
| NAD+ | 0 mM |
| CoA | 0 mM |
| ATP | 0 mM |

| | |
|---|---|
| *Ec* ACAT | variable |
| *Sc* HMGS | + |
| *Bp* HMGR | 0.2 μM |
| *Mm* MK | 0.1 μM |
| *Sp* PMK | 0.2 μM |
| *Sc* PMD | 0.2 μM |
| *Scl* IDI | 0.2 μM |
| *Pg* GPPS | 1.0 μM |
| *Ms* LS | + |
| NAD+ | 5 mM |
| CoA | 1 mM |
| ATP | 1 mM |

| | |
|---|---|
| *Ec* ACAT | + |
| *Sc* HMGS | variable |
| *Bp* HMGR | 0.2 μM |
| *Mm* MK | 0.1 μM |
| *Sp* PMK | 0.2 μM |
| *Sc* PMD | 0.2 μM |
| *Scl* IDI | 0.2 μM |
| *Pg* GPPS | 1.0 μM |
| *Ms* LS | + |
| NAD+ | 5 mM |
| CoA | 1 mM |
| ATP | 1 mM |

**Figure S6. Use of pre-enriched lysates for LS, ACAT, and HMGS improves limonene production and increases available volumetric space in the reaction for other CFPS-generated enzymes.** (**A**) Titration of limonene synthase from *Mentha spicate* (*Ms*LS) generated by CFPS (0.5-2 μM) or pre-enriched lysate (*, ~3.4 μM) suggests LS is a rate-limiting step for limonene production. (**B-C**) Titration of acetyl-CoA acetyltransferase from *Escherichia coli* (*Ec*ACAT) generated by CFPS (0.5-2.5 μM) or pre-enriched lysate (*, ~4.6 μM) suggests 0.5 μM ACAT is the minimum enzyme concentration sufficient to produce 2 mM of limonene. (**D-E**) Titration of 3-hydroxy-3-

methylglutaryl-CoA synthase from *Saccharomyces cerevisiae* (*Sc*HMGS) generated by CFPS (0.2-4 μM) or pre-enriched lysate (*, ~5.5 μM) suggests 3 μM HMGS is the minimum enzyme concentration sufficient to produce 2 mM of limonene. The standard iPROBE reaction used throughout this work dictates that only 15 μL of the 30 μL reaction volume is available for CFPS reactions; using pre-enriched lysates for ACAT, HMGS, LS not only maximized limonene titer (in the case of LS) but also frees up space for other pathway enzymes generated by CFPS to be included at a higher concentration in the limonene synthesis reaction. Concentrations of ACAT, HMGS, LS in pre-enriched lysates are estimated by densitometry (**Table S2**). The values for limonene at 24 hours represent averages (n=3) and error bars represent 1 standard deviation. The error bars associated with TREE scores represent propagated error (see **Appendix S1**).

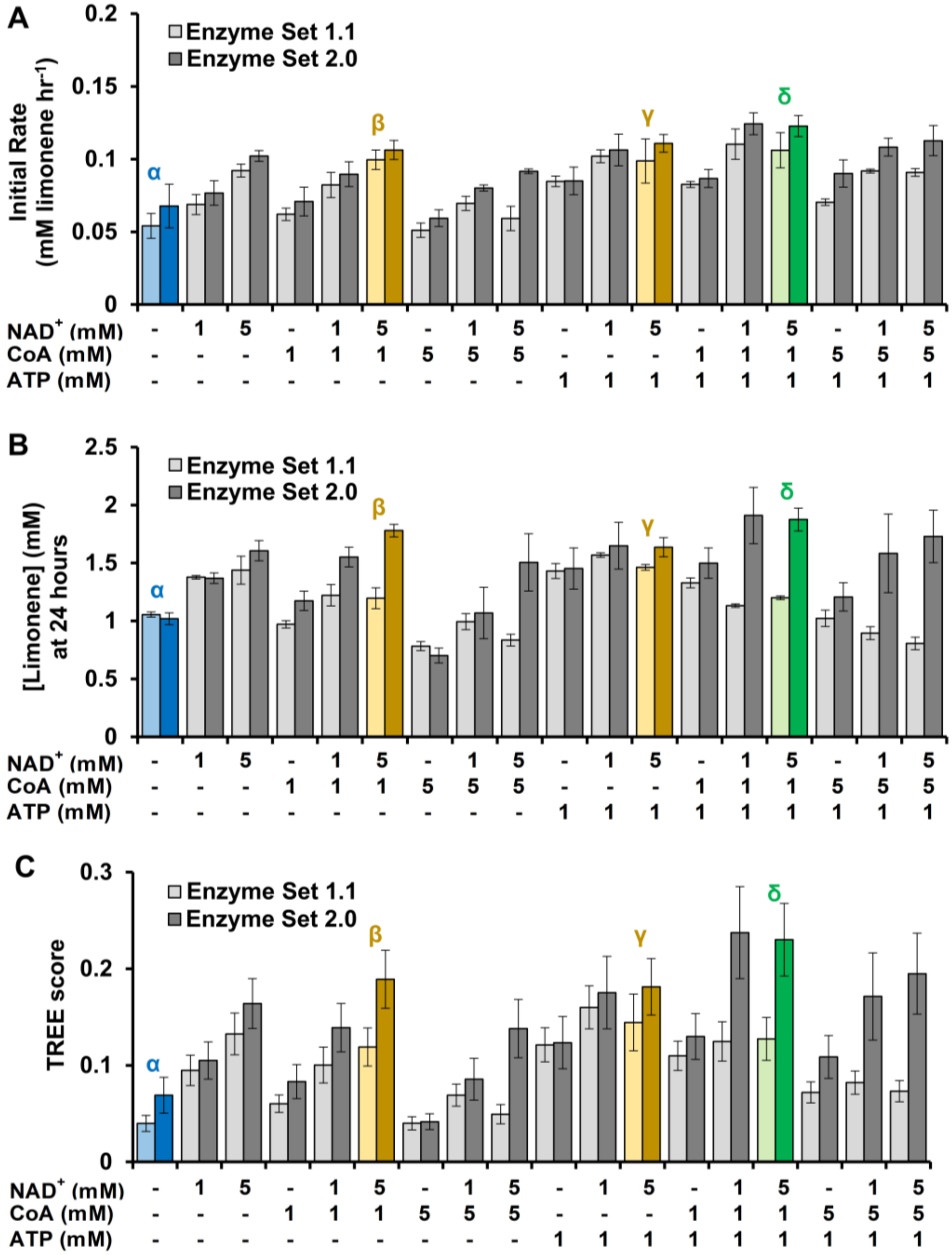


**Figure S7. Supplementation of cofactors NAD$^+$, CoA, and ATP improves limonene productivity (A), titer (B), and TREE score (C).** The improved set of enzyme homologs (enzyme set 2.0) has increased productivity and titer compared to the initial set of enzyme homologs (enzyme set 1.0) in nearly all cofactor conditions tested. Four reaction conditions (α, β, γ, δ) were selected as cofactor conditions for further experiments. Note that plot C is a bar chart representation of the data in **Figure 3B.** The initial rate value is the slope of a linear regression of four data points at 3, 4, 5, and 6 hours and error bars are the associated standard error (see **Appendix S1**). The values for limonene at 24 hours represent averages (n=3) and error bars represent 1 standard deviation. The error bars associated with TREE scores represent propagated error (see **Appendix S1**).

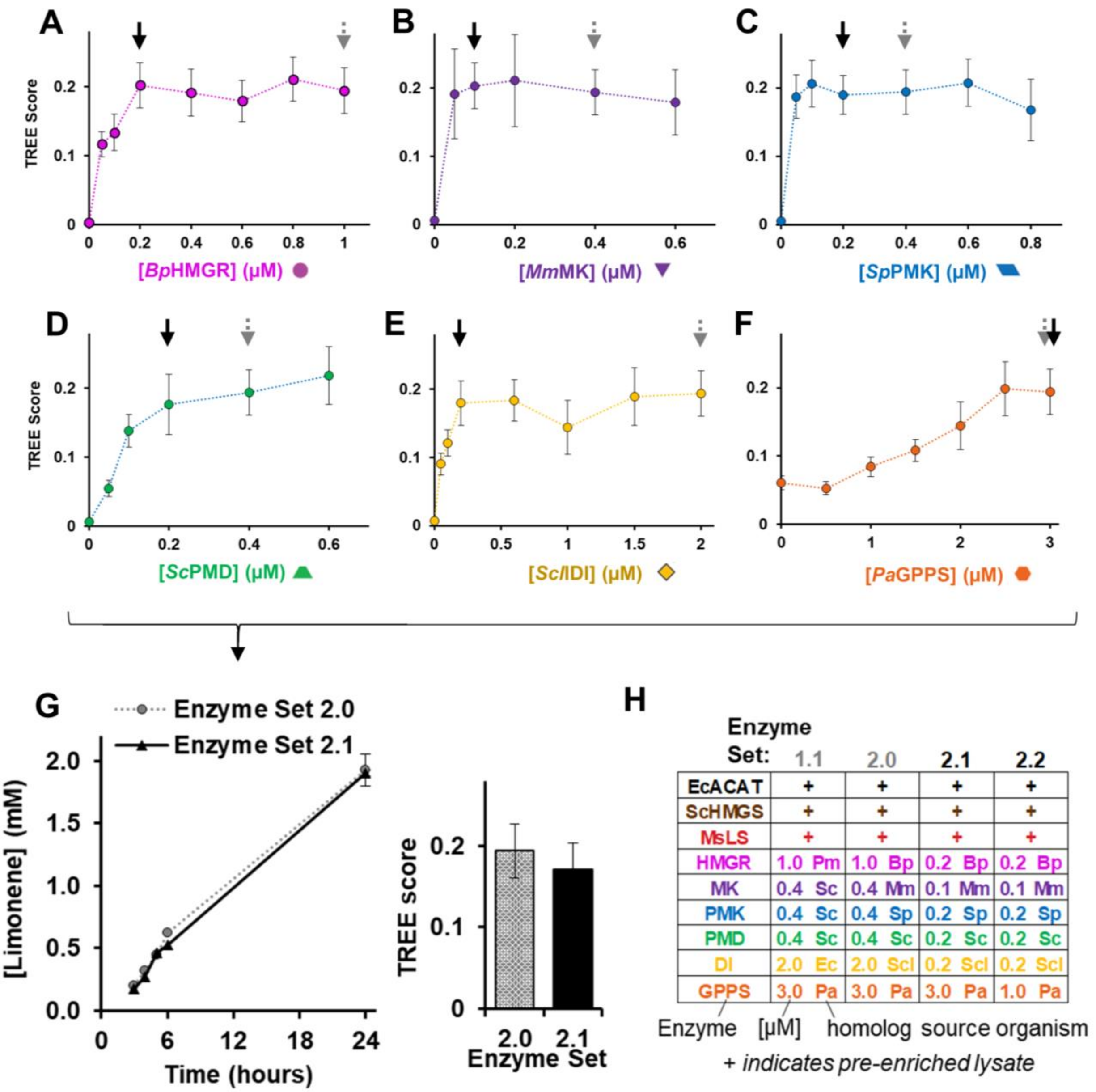


**Figure S8. Reducing enzyme concentrations to enable equivalent limonene production.** (**A-F**) Plots of limonene TREE score as a function of enzyme concentration using cofactor condition δ (5 mM $NAD^+$, 1 mM CoA, 1 mM ATP). Pathway enzymes not being adjusted are held at the same concentrations used in enzymes set 2.0. Enzyme set 2.1 uses the same enzyme homologs as set 2.0 but reduces the concentrations of *Bp*HMGR, *Mm*MK, *Sp*PMK, *Sc*PMD, and *Scl*IDI. Black arrows indicate the concentration of each enzyme in set 2.0 while gray arrows indicate concentration of each enzyme in set 2.1. Values represent TREE scores and error bars represent propagated error (see

**Appendix S1**). (**G**) After testing the effect of concentration on each enzyme individually, the concentrations of HMGR, MK, PMK, PMD, and IDI were reduced to create enzyme set 2.1 which has a similar limonene production and TREE score to enzyme set 2.0. Initial rate values are n=1 while the 24 hour value is an average (n=3) with error bars which represent 1 standard deviation. (**H**) Summary of enzyme set components used during cofactor and concentration optimization.

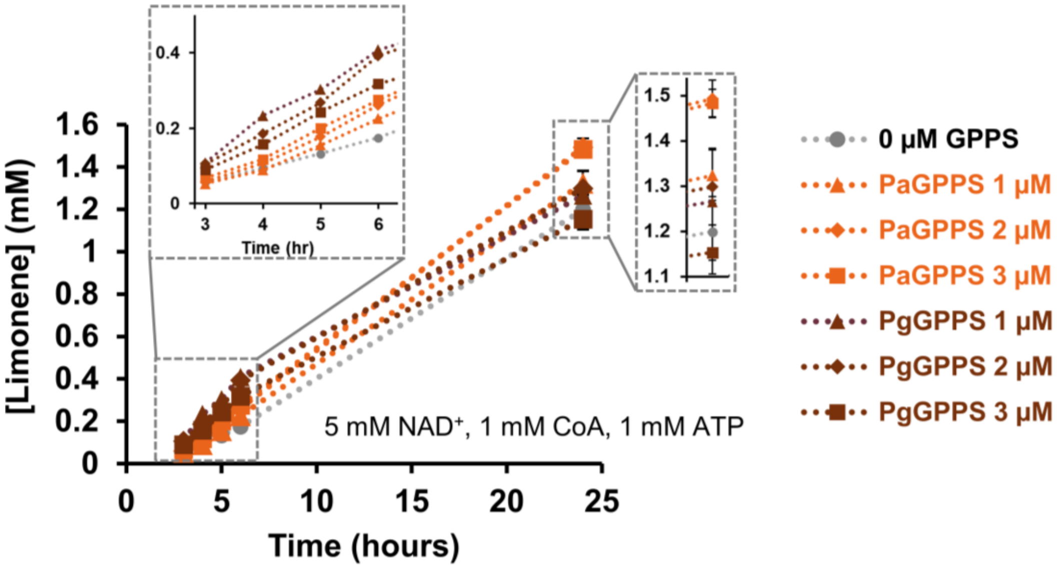


**Figure S9. Limonene production using two different GPPS enzymes at varying concentration**. The default homolog for GPPS throughout the study thus far has been *Pa*GPPS; increasing the concentration of PaGPPS maximizes limonene titer (3 µM is one of the largest concentrations possible without exceeding the 15 µL of CFPS reaction included in a 30 µL limonene synthesis reaction). However, comparison with *Pg*GPPS indicates that other GPPS homologs do not require such high enzyme concentrations to support the same limonene titer and that using 1 µM as the benchmark comparison is appropriate for future testing. Thus, enzyme set 2.2 (the starting point in **Figure 4**) reduces the concentration of GPPS from 3.0 µM to 1.0 µM in order to more accurately compare the specific activity of GPPS homologs in the next round of homolog screening. Initial rate values are n=1 while the 24 hour value is an average (n=3) with error bars which represent 1 standard deviation.

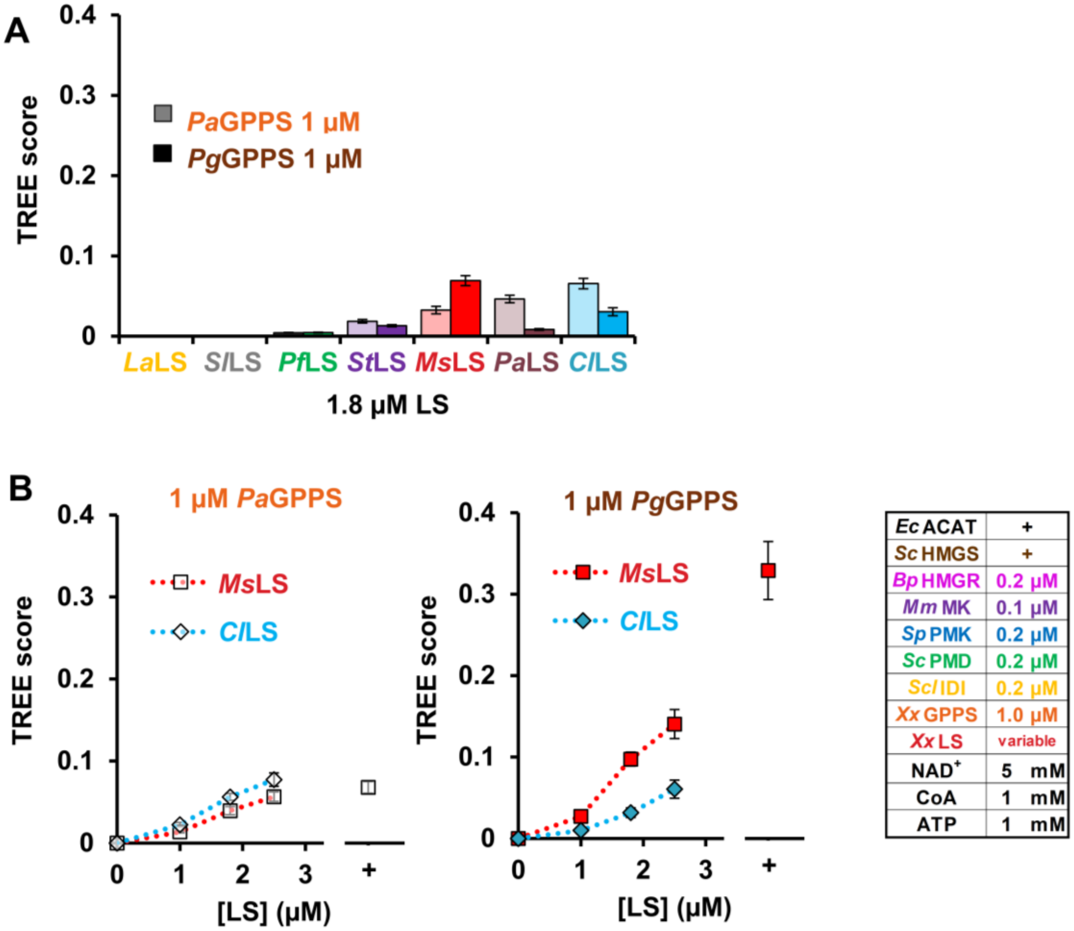


**Figure S10. Testing limonene synthases under improved cofactors and decreased enzyme concentrations.** (**A**) Comparison of seven limonene synthase homologs at 1.8 µM concentration. (**B**) Comparison of *Ms*LS (from *Mentha spicata*) and *Cl*LS (from *Citrus limon*) at different concentration and paired with two differed GPPS enzymes. *Ms*LS is known to produce (4S)-(-)-limonene (Colby et al., 1993) while *Cl*LS typically produces (4R)-(+)-limoenene.(Lücker et al., 2002). (+) indicates a pre-enriched lysate with an approximate enzyme concentration in the reaction of ~3.4 µM. Values represent TREE scores and error bars represent propagated error (see **Appendix S1**).

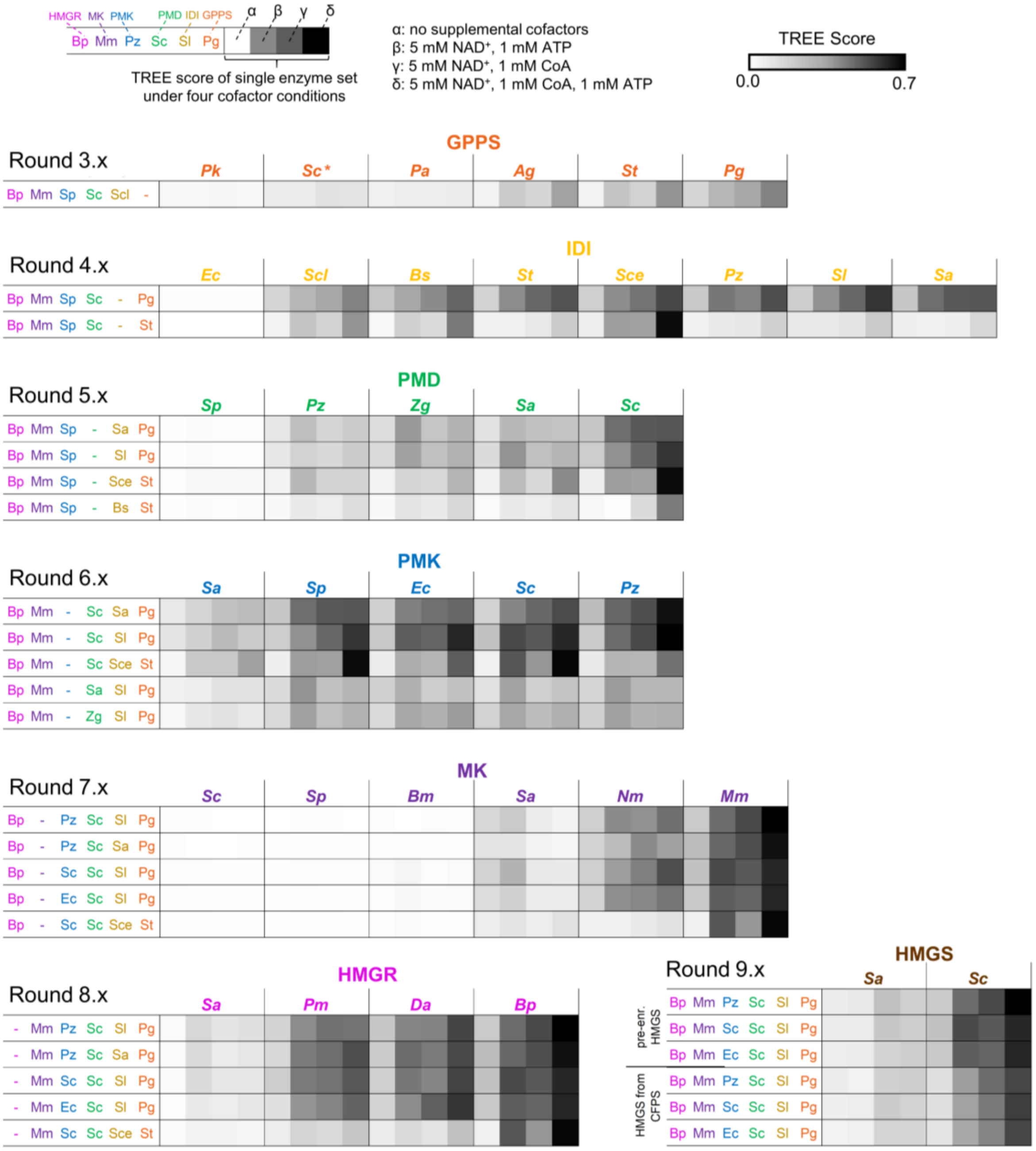


**Figure S11. TREE scores of 408 reactions where each shaded square is a unique reaction condition.** Each unique enzyme set is measured at four different cofactor concentrations (α, β, γ, δ). A duplicate representation of the same data presented in **Figure 4B**.

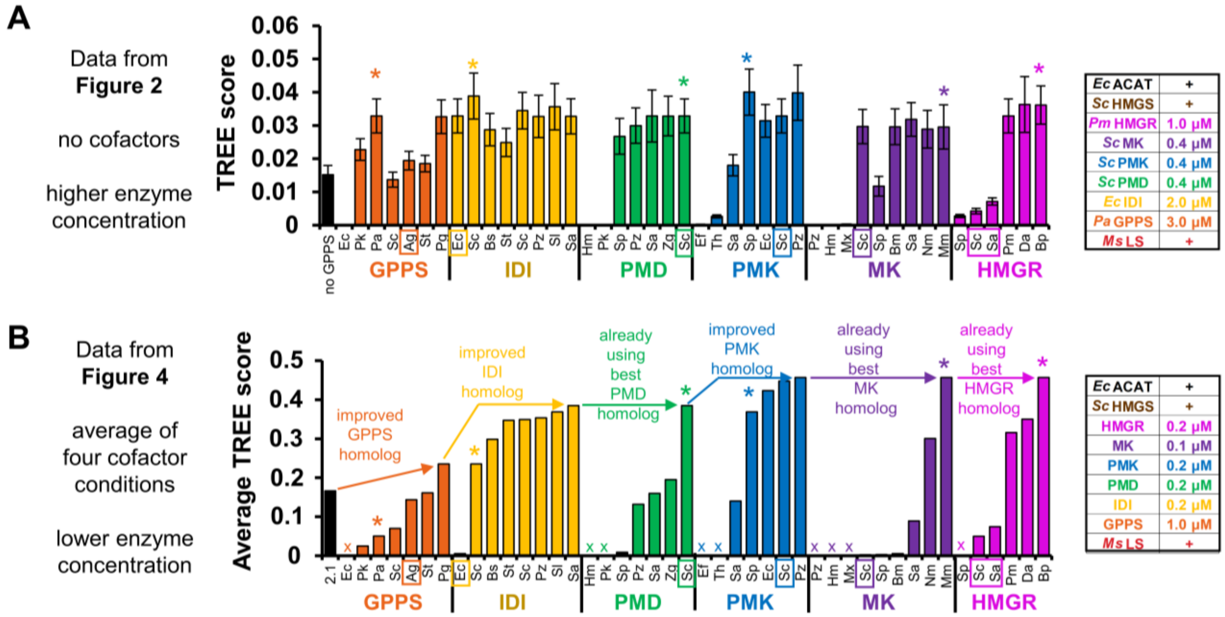


**Figure S12. A more stringent selection condition differentiates similar homologs.** (**A**) The TREE scores from **Figure 2E** are replotted; * denoted the enzyme selected as the best candidate when transitioning from enzyme set 1.0 to enzyme set 2.0. (**B**) The average TREE scores from **Figure 4C** are replotted**.** Homologs marked with "x" were inactive in **Figure 2** and were not retested in **Figure 4**. The boxed homologs (*Ag*GPPS, *Ec*IDI, *Sc*PMD, *Sc*PMK, *Sc*MK, and *Sa*HMGR/*Sc*HMGR) indicate some of the commonly used enzyme homologs utilize for metabolic engineering in living cells, for example the production of limonene (Alonso-Gutierrez et al., 2013; Alonso-Gutierrez et al., 2015) and pinene (Sarria et al., 2014). Values represent TREE scores and error bars (panel A) represent propagated error (see **Appendix S1**).

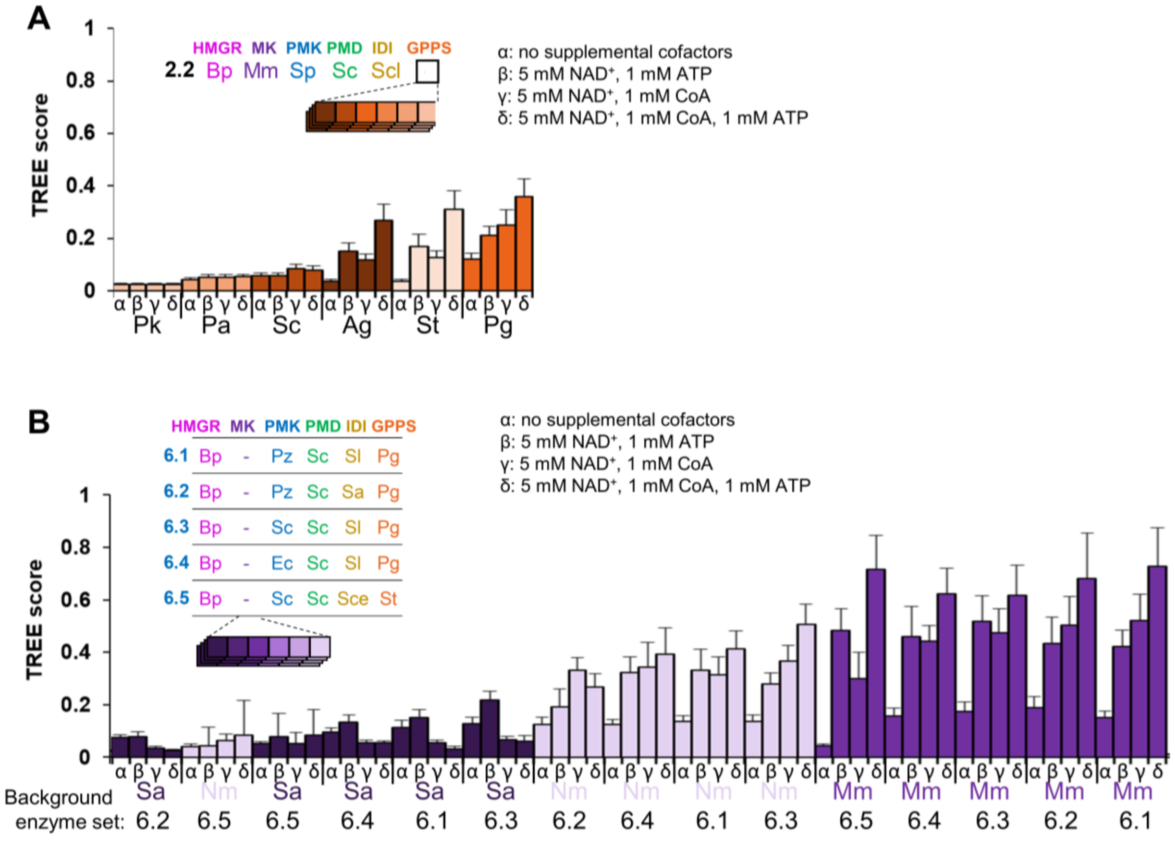


**Figure S13. TREE scores of selected reactions depicted in Figure 4B.** *Sa*MK is inhibited by higher CoA concentrations as compared to other MK homologs. Values represent TREE scores and error bars represent propagated error (see **Appendix S1**).

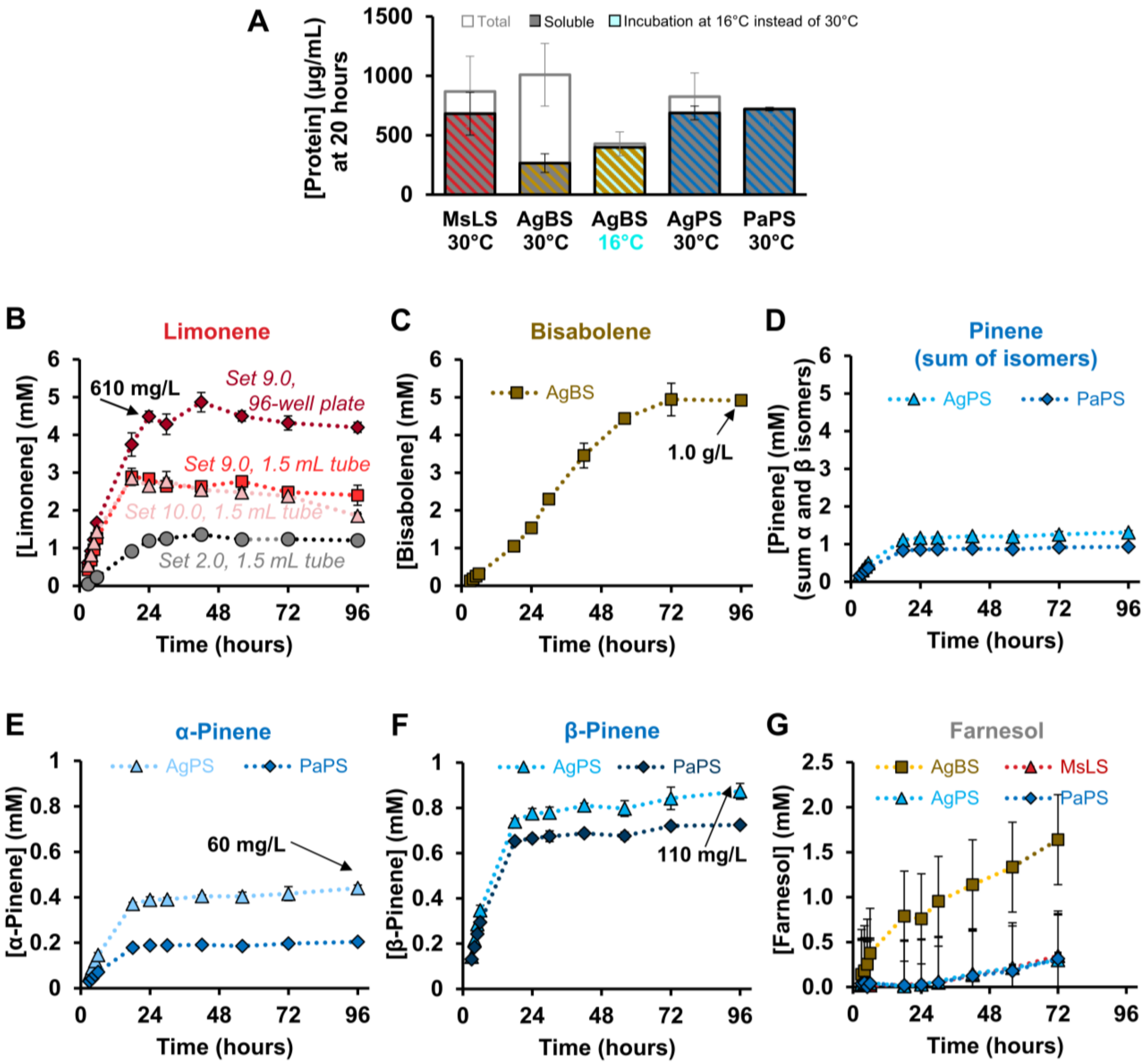


**H**

| Pinene Synthase | Purified enzymes *in vitro* | This study - lysates *in vitro* | (Sarria et al., 2014) *in vivo* |
|---|---|---|---|
| *Abies grandis* grand fir | 52% α-Pinene<br>48% β-Pinene | 33% α-Pinene<br>67% β-Pinene | 50% α-Pinene<br>50% β-Pinene |
| *Picea abies* Norway spruce | 57% α-Pinene<br>27% β-Pinene | 22% α-Pinene<br>78% β-Pinene | 47% α-Pinene<br>53% β-Pinene |

**Figure S14. Production of limonene, pinene, and bisabolene over 96 hours.** (**A**) Cell-free protein synthesis of terpenoid synthases; incubation at 16 °C improves expression of bisabolene synthase from *Abies grandis*. (**B**) Production of limonene from glucose using Enzymes Sets 2.0, 9.0, and 10.0. Incubation of reactions in a 96-well plate covered in foil rather than a 1.5 mL microcentrifuge tube further increased titer. Set 9.0 uses a pre-enriched MsLS lysate while set 10.0 generates MsLS using CFPS (3.8 μM). (**C**) Production of bisabolene from glucose by CFPS-enriched reactions containing farnesyl diphosphate synthase (EcFPPS encoded by *ispA*) from *E. coli* and bisabolene synthase (BS) from *Abies grandis*. (**D**) Production of pinene from glucose by CFPS-enriched reactions containing pinene synthases (PS) from *Abies grandis* (Grand fir) and *Picea abies* (Norway spruce). (**E-F**) Pinene exists as two isomers (α or β) that can be distinguished by GC-MS due to different fragmentation patterns and retention times. For biofuel applications, there is no preference for either isomer, however, β-pinene is a more valuable commodity chemical since it is less abundant than α-pinene in turpentine (Sarria et al., 2014). β-pinene is favored in this *in vitro* system. (**G**) Farnesol accumulates in all reactions conditions but especially in the bisabolene reaction. (**H**) The ratio of α or β pinene produced using purified pinene synthases (*A. grandis* (Bohlmann et al., 1997) and *P. abies* (Martin et al., 2004)), this study, and production in cells (Sarria et al., 2014). Unless noted, limonene, pinene, and bisabolene synthesis reactions use enzyme set 10.0 (see **Table S3**) and are supplemented with 5 mM $NAD^+$, 1 mM CoA, and 1 mM ATP. Values represent averages (n=3) and error bars represent 1 standard deviation.

#### Appendix S1. Equations

Equation 1:

$$\frac{\mu mol\ (14\text{C counts})\ leucine\ in\ protein}{\mu mol (14C\ counts)\ leucine\ in\ rxn} \times \frac{2010\ \mu mol\ leucine\ in\ rxn}{1\ L\ rxn} \times \frac{\mu mol\ protein}{\text{"\#"}\ \mu mol\ leucine} \times \frac{\text{"mw"}\ \mu g\ protein}{\mu mol\ protein} \times \frac{1\ L}{1000\ mL} = \frac{x\ \mu g\ protein}{mL}$$

Note that "#" is number of leucine amino acids in the expressed protein and "mw" is the protein molecular weight (g/mol).

Equation 2:

$$TREE\ Score = Titer\ \cdot Rate\ \cdot (Average\ enzyme\ solubility)$$

The TREE (Titer, Rate, and Enzyme Expression) score is an aggregate value of enzyme solubility, initial productivity, and final titer *(Karim et al., 2019).* Specifically, the TREE score is the product of initial productivity (mM/hr), final titer at 24 hr (mM), and average solubility of 9 overexpressed proteins (%). Error from each component value is propagated to obtain an estimate of TREE score uncertainty, see Equation 3.

Equation 3:

$$TREE\ Score\ error =\ TREE\ score \cdot \sqrt{\left(\frac{\sigma_{titer}}{Titer}\right)^2 + \left(\frac{\sigma_{rate}}{Rate}\right)^2 + \left(\frac{Average\ enzyme\ solubility\ error}{Average\ enzyme\ solubility}\right)^2}$$

The titer (mM) is the limonene concentration in the cell-free reaction at 24 hours and $\sigma_{titer}$ = standard deviation of the three experimental replicates. The rate (mM/hr) is an estimate of initial productivity and is the slope of a linear regression of four single replicate limonene concentrations measured at 3, 4, 5, and 6 hours. $\sigma_{rate}$ is the standard error for the $m_1$ slope term of the LINEST regression calculation and is determined using the following formula in Microsoft Excel: =INDEX(LINEST([3hr titer, 4hr titer, 5hr titer, 6hr titer],[3,4,5,6],1,1),2,1)

The average enzyme solubility is an average of the individual CFPS solubility values for the nine enzymes of the limonene pathway. Note that the values for ACAT, HMGS, and LS were included in the TREE score regardless of whether these enzymes were supplied as pre-enriched lysates or CFPS reactions. For each individual enzyme, the enzyme solubility and enzyme solubility error were calculated from (at minimum) six enzyme concentrations derived from $^{14}$C-leucine incorporation experiments. The average and standard deviation were calculated based on n ≥3 replicates for soluble protein ($\overline{soluble}, \sigma_{soluble}$) and n ≥3 replicates for total protein ($\overline{total}, \sigma_{total}$).

$$Enzyme\ solubility\ (\%)\ for\ a\ single\ enzyme = \mu_{solubility} = \frac{\overline{soluble}}{\overline{total}}$$

$$Enzyme\ solubility\ error\ (\%)\ for\ a\ single\ enzyme = \sigma_{solubility} = \frac{\overline{soluble}}{\overline{total}} \cdot \sqrt{\left(\frac{\sigma_{soluble}}{\overline{soluble}}\right)^2 + \left(\frac{\sigma_{total}}{\overline{total}}\right)^2}$$

The average enzyme solubility (and error) is then calculated as follows, where the n index describes the number of each enzyme in the pathway (i.e. ACAT = 1, HMGS = 2, etc.)

$$Average\ enzyme\ solubility\ (\%) = \sum_{n=1}^{n=9} \frac{(\mu_{solubility})_n}{9}$$

$$Average\ enzyme\ solubility\ error\ (\%) = \sqrt{\sum_{n=1}^{n=9} \frac{\left((\sigma_{solubility})_n\right)^2}{9}}$$

#### Appendix S2. Gene Sequences

Here is a color code for sequences used: XbaI, ribosome binding site (RBS) from pJL1, NdeI (containing ATG start codon), SalI, stop codon, and the N-terminal expression tag (MEKKI or MHMEKKI) sequence is underlined.

##### 354_pJL1-atoB (ACAT_Eco)

TCTAGAAATAATTTTGTTTAACTTTAAGAAGGAGATATACATATGAAAAATTGTGTCA
TCGTCAGTGCGGTACGTACTGCTATCGGTAGTTTTAACGGTTCACTCGCTTCCACC
AGCGCCATCGACCTGGGGGCGACAGTAATTAAAGCCGCCATTGAACGTGCAAAAA
TCGATTCACAACACGTTGATGAAGTGATTATGGGTAACGTGTTACAAGCCGGGCTG
GGGCAAAATCCGGCGCGTCAGGCACTGTTAAAAAGCGGGCTGGCAGAAACGGTG
TGCGGATTCACGGTCAATAAAGTATGTGGTTCGGGTCTTAAAAGTGTGGCGCTTGC
CGCCCAGGCCATTCAGGCAGGTCAGGCGCAGAGCATTGTGGCGGGGGGTATGGA
AAATATGAGTTTAGCCCCCTACTTACTCGATGCAAAAGCACGCTCTGGTTATCGTCT
TGGAGACGGACAGGTTTATGACGTAATCCTGCGCGATGGCCTGATGTGCGCCACC
CATGGTTATCATATGGGGATTACCGCCGAAAACGTGGCTAAAGAGTACGGAATTAC
CCGTGAAATGCAGGATGAACTGGCGCTACATTCACAGCGTAAAGCGGCAGCCGCA
ATTGAGTCCGGTGCTTTTACAGCCGAAATCGTCCCGGTAAATGTTGTCACTCGAAA
GAAAACCTTCGTCTTCAGTCAAGACGAATTCCCGAAAGCGAATTCAACGGCTGAAG
CGTTAGGTGCATTGCGCCCGGCCTTCGATAAAGCAGGAACAGTCACCGCTGGGAA
CGCGTCTGGTATTAACGACGGTGCTGCCGCTCTGGTGATTATGGAAGAATCTGCG
GCGCTGGCAGCAGGCCTTACCCCCCTGGCTCGCATTAAAAGTTATGCCAGCGGTG
GCGTGCCCCCCGCATTGATGGGTATGGGGCCAGTACCTGCCACGCAAAAAGCGTT
ACAACTGGCGGGGCTGCAACTGGCGGATATTGATCTCATTGAGGCTAATGAAGCA
TTTGCTGCACAGTTCCTTGCCGTTGGGAAAAACCTGGGCTTTGATTCTGAGAAAGT
GAATGTCAACGGCGGGGCCATCGCGCTCGGGCATCCTATCGGTGCCAGTGGTGC
TCGTATTCTGGTCACACTATTACATGCCATGCAGGCACGCGATAAAACGCTGGGG
CTGGCAACACTGTGCATTGGCGGCGGTCAGGGAATTGCGATGGTGATTGAACGGT
TGAATTAAGTCGAC

##### 310_pJL1-(CAT5aa)-HMGS_Sce

TCTAGAAATAATTTTGTTTAACTTTAAGAAGGAGATATACATATGGAGAAAAAAATCA
AACTGAGCACCAAGCTGTGCTGGTGTGGCATCAAGGGTCGCCTGCGCCCACAAAA
GCAGCAACAGCTGCACAACACGAACCTGCAAATGACCGAGCTGAAAAAGCAGAAG
ACGGCCGAGCAAAAGACCCGCCCGCAGAACGTTGGCATCAAGGGCATCCAGATTT
ATATCCCGACGCAGTGTGTCAACCAATCTGAGCTGGAGAAATTCGATGGCGTCAG
CCAGGGTAAGTACACCATCGGCCTGGGCCAGACCAACATGAGCTTCGTGAACGAC
CGTGAGGACATCTATTCTATGAGCCTGACGGTGCTGTCTAAGCTGATCAAGAGCTA

CAACATCGACACGAATAAGATCGGTCGTCTGGAGGTGGGTACGGAGACGCTGATT
GACAAGAGCAAAAGCGTGAAGTCTGTCTTAATGCAGCTGTTCGGCGAGAACACGG
ATGTCGAGGGTATCGACACCCTGAACGCGTGTTACGGCGGCACCAACGCACTGTT
CAATAGCCTGAACTGGATTGAGAGCAACGCCTGGGATGGCCGCGATGCGATCGTC
GTGTGCGGCGATATCGCCATCTATGACAAGGGTGCGGCACGTCCGACCGGCGGT
GCAGGCACCGTTGCGATGTGGATTGGCCCGGACGCACCAATTGTCTTCGATTCTG
TCCGCGCGTCTTACATGGAGCACGCCTACGACTTTTACAAGCCGGACTTCACGAG
CGAATACCCGTACGTGGACGGCCACTTCTCTCTGACCTGCTATGTGAAGGCGCTG
GACCAGGTTTATAAGTCTTATAGCAAAAAGGCGATTTCTAAGGGCCTGGTCAGCGA
CCCGGCAGGCAGCGACGCCCTGAACGTGCTGAAGTATTTCGACTACAACGTGTTC
CATGTCCCGACCTGCAAATTAGTGACCAAATCTTATGGCCGCCTGTTATATAATGAT
TTCCGTGCCAACCCGCAGCTGTTCCCGGAGGTTGACGCCGAGCTGGCGACGCGT
GATTACGACGAGAGCCTGACCGACAAGAACATCGAGAAGACCTTCGTCAACGTCG
CGAAGCCGTTCCACAAAGAGCGTGTGGCCCAAAGCCTGATCGTCCCGACCAACAC
GGGCAACATGTATACCGCGTCTGTCTACGCGGCATTCGCGAGCCTGCTGAATTAC
GTCGGTTCTGACGACCTGCAGGGCAAGCGCGTTGGCCTGTTCAGCTACGGTAGC
GGCTTAGCGGCCAGCCTGTATAGCTGCAAAATTGTCGGCGACGTCCAGCACATCA
TCAAGGAGCTGGACATCACCAACAAGCTGGCGAAGCGCATCACCGAGACGCCGA
AAGATTACGAGGCAGCGATCGAGTTACGCGAGAATGCGCATCTGAAGAAGAACTT
CAAGCCGCAAGGTAGCATCGAGCACCTGCAGAGCGGCGTCTACTACCTGACGAAC
ATTGACGACAAGTTCCGCCGTTCTTATGACGTCAAAAAGCTGGAGTGAGTCGAC

**311_pJL1-(CAT5aa)-HMGS_Sau**

TCTAGAAATAATTTTGTTTAACTTTAAGAAGGAGATATACATATGGAGAAAAAAATCA
CCCTGGGCATTGATAAAATCAACTTCTATGTGCCGAAATATTACGTGGATATGGCA
AAACTGGCAGAAGCACGTCAGGTTGATCCGAACAAATTTCTGATTGGTATTGGTCA
GACCGAAATGGCAGTTAGTCCGGTTAATCAGGATATTGTTAGCATGGGTGCAAATG
CAGCCAAAGATATTATCACCGATGAAGATAAAAAAAAAATCGGCATGGTTATCGTT
GCAACCGAAAGCGCAGTTGATGCAGCAAAAGCAGCAGCAGTTCAGATTCATAATCT
GCTGGGTATTCAGCCGTTTGCACGTTGTTTTGAAATGAAAGAAGCATGTTACGCAG
CAACACCGGCAATTCAGCTGGCAAAAGATTATCTGGCAACCCGTCCGAATGAAAAA
GTTCTGGTTATTGCCACCGATACCGCACGTTATGGTCTGAATAGCGGTGGTGAACC
GACCCAGGGTGCCGGTGCAGTTGCAATGGTTATTAGCCATAATCCGAGCATTCTG
GCACTGAATGAAGATGCAGTTGCCTATACCGAAGATGTGTATGATTTTTGGCGTCC
GACCGGTCATAAATATCCGCTGGTTGATGGTGCACTGAGCAAAGATGCATATATTC
GTAGCTTTCAGCAGAGCTGGAATGAATATGCAAAACGTCAGGGTAAAAGCCTGGC
AGATTTTGCAAGCCTGTGTTTTCATGTTCCGTTTACCAAAATGGGTAAAAAAGCCCT
GGAAAGCATTATTGATAATGCCGATGAAACCACCCAAGAACGTCTGCGTAGCGGTT
ATGAGGATGCCGTTGATTATAACCGTTATGTGGGTAACATTTATACCGGTAGCCTG
TATCTGAGCCTGATTAGCCTGCTGGAAAATCGTGATCTGCAGGCAGGCGAAACCA
TTGGTCTGTTTAGCTATGGTAGCGGTAGCGTTGGTGAGTTTTATAGCGCAACCCTG
GTTGAAGGTTATAAAGATCATCTGGATCAGGCAGCACATAAAGCACTGCTGAATAA

TCGTACCGAAGTTAGCGTTGATGCGTATGAAACCTTTTTCAAACGCTTCGATGATGT
GGATTTTGATGAACAGCAGGATGCAGTTCATGAAGATCGCCATATCTTTTATCTGA
GCAACATTGAAAACAATGTGCGCGAATATCATCGTCCGGAATAAGTCGAC

**312_pJL1-(CAT5aa)-HMGR_Sce**

TCTAGAAATAATTTTGTTTAACTTTAAGAAGGAGATATACATATGGAGAAAAAAATC
GTGCTGACGAACAAAACCGTCATTAGCGGCAGCAAGGTGAAGTCTCTGAGCAGCG
CCCAAAGCTCTAGCAGCGGCCCGTCTAGCAGCAGCGAGGAGGACGACAGCCGTG
ACATTGAGTCTCTGGACAAGAAGATCCGCCCGCTGGAGGAGTTAGAGGCCCTGCT
GAGCAGCGGCAACACCAAGCAGCTGAAGAACAAGGAAGTTGCAGCGCTGGTGAT
CCACGGTAAGCTGCCACTGTATGCGCTGGAAAAGAAACTGGGCGATACGACGCGT
GCGGTCGCGGTGCGTCGCAAAGCCTTAAGCATCTTAGCGGAGGCCCCGGTGTTA
GCCAGCGACCGCCTGCCGTACAAGAACTACGACTACGACCGCGTGTTTGGCGCG
TGCTGCGAGAATGTCATTGGCTACATGCCGTTACCGGTTGGTGTGATCGGCCCGC
TGGTCATTGATGGCACGAGCTATCACATTCCAATGGCGACCACGGAAGGTTGCTTA
GTCGCCAGCGCCATGCGTGGCTGTAAGGCGATTAACGCCGGCGGTGGCGCGACG
ACCGTGTTAACCAAGGATGGTATGACGCGCGGTCCGGTCGTCCGCTTCCCAACGC
TGAAGCGCAGCGGCGCGTGTAAGATTTGGCTGGATTCTGAGGAGGGCCAAAACG
CGATCAAGAAAGCCTTCAACTCTACGAGCCGTTTCGCGCGTTTACAGCATATCCAG
ACCTGCCTGGCCGGCGACCTGCTGTTCATGCGCTTCCGCACCACCACGGGCGAT
GCGATGGGCATGAACATGATCAGCAAGGGCGTCGAATATAGCCTGAAACAAATGG
TGGAAGAATATGGCTGGGAGGACATGGAGGTTGTCTCTGTGAGCGGCAACTATTG
CACCGACAAGAAGCCGGCAGCCATTAACTGGATTGAGGGTCGCGGCAAAAGCGT
CGTGGCAGAAGCGACCATCCCAGGCGACGTGGTCCGTAAGGTTCTGAAGAGCGA
CGTCAGCGCCCTGGTTGAGTTAAATATCGCGAAAAACCTGGTCGGCAGCGCGATG
GCGGGCAGCGTGGGTGGCTTTAACGCACATGCAGCGAATCTGGTTACGGCGGTTT
TCTTAGCCTTAGGTCAGGACCCAGCCCAAAATGTCGAGAGCAGCAACTGCATTAC
CTTAATGAAAGAGGTTGACGGTGACCTGCGCATCAGCGTTTCTATGCCGTCTATCG
AGGTCGGCACGATCGGCGGCGGCACCGTTTTAGAACCGCAAGGTGCGATGCTGG
ATCTGCTGGGCGTGCGCGGCCCACATGCAACGGCCCCAGGCACCAATGCCCGCC
AACTGGCCCGTATCGTGGCCTGCGCGGTTCTGGCGGGTGAGCTGAGCCTGTGCG
CCGCATTAGCCGCGGGCCATTTAGTTCAATCTCACATGACCCACAACCGCAAGCC
GGCAGAACCAACCAAGCCAAATAACCTGGACGCAACCGACATTAACCGTCTGAAG
GATGGCAGCGTCACGTGCATTAAAAGCTAAGTCGAC

**313_pJL1-(CAT5aa)-HMGR_Sau**

TCTAGAAATAATTTTGTTTAACTTTAAGAAGGAGATATACATATGGAGAAAAAAATC
CAGAGCCTGGATAAAAACTTTCGTCATCTGAGCCGTCAGCAGAAACTGCAGCAGC
TGGTTGATAAACAGTGGCTGAGCGAAGAACAGTTTAACATTCTGCTGAATCATCCG
CTGATTGATGAAGAAGTTGCAAACAGCCTGATTGAAAATGTTATTGCACAGGGTGC

ACTGCCGGTTGGTCTGCTGCCGAACATTATTGTTGATGATAAAGCATATGTGGTGC
CGATGATGGTTGAAGAACCGAGCGTTGTTGCAGCAGCAAGCTATGGTGCAAAACT
GGTTAATCAGACCGGTGGCTTTAAAACCGTTAGCAGCGAACGTATTATGATTGGCC
AGATTGTTTTTGATGGTGTGGATGATACCGAAAAACTGAGCGCAGATATTAAAGCC
CTGGAAAAACAAATTCATCAGATTGCCGATGAAGCCTATCCGAGCATTAAAGCACG
TGGTGGTGGTTATCAGCGTATTGCAATTGATACCTTTCCGGAACAGCAACTGCTGA
GCCTGAAAGTGTTTGTTGATACCAAAGATGCAATGGGTGCCAATATGCTGAATACC
ATTCTGGAAGCAATTACCGCCTTTCTGAAAAATGAATTTCCGCAGAGCGATATCCT
GATGAGCATTCTGAGCAATCATGCAACCGCAAGCGTTGTTAAAGTTCAGGGTGAAA
TTGATGTTAAAGACCTGGCACGCGGTGAACGTACCGGTGAAGAGGTTGCCAAACG
TATGGAACGTGCAAGCGTTCTGGCACAGGTTGATATTCATCGTGCAGCAACCCATA
ATAAAGGTGTGATGAACGGTATTCATGCAGTTGTTCTGGCAACCGGTAATGATACC
CGTGGTGCGGAAGCAAGCGCACATGCATATGCCAGCAAAGATGGTCAGTATCGTG
GTATTGCAACCTGGCGTTATGATCAAGAACGTCAGCGTCTGATTGGTACAATTGAA
GTTCCGATGACCCTGGCAATTGTTGGTGGTGGCACCAAAGTTCTGCCGATTGCAAA
AGCAAGCCTGGAACTGCTGAATGTTGAAAGCGCACAAGAACTGGGTCATGTTGTT
GCCGCAGTGGGTCTGGCCCAGAATTTTGCAGCATGTCGTGCACTGGTTAGCGAAG
GTATCCAGCAGGGTCATATGAGTCTGCAGTATAAAAGCCTGGCCATTGTGGTTGGT
GCCAAAGGTGATGAAATTGCGCAGGTTGCAGAAGCACTGAAACAAGAACCGCGTG
CAAATACCCAGGTTGCGGAACGTATTCTGCAGGATCTGCGTAGCCAGCAGTAAGT
CGAC

**314_pJL1-(CAT5aa)-HMGR_Pme**

TCTAGAAATAATTTTGTTTAACTTTAAGAAGGAGATATACATATGGAGAAAAAAATCA
GCTTGGACAGCCGCCTGCCAGCTTTTCGGAACCTGAGCCCCGCGGCGCGCCTGG
ATCATATTGGCCAGCTCTTGGGCCTGAGCCATGATGACGTGAGCCTGCTGGCGAA
CGCGGGAGCGCTGCCGATGGATATTGCGAACGGCATGATTGAGAACGTGATTGG
CACCTTTGAACTGCCATACGCGGTGGCGAGCAACTTTCAGATTAATGGCCGGGAC
GTGCTGGTGCCGCTGGTGGTGGAGGAACCAAGCATTGTGGCGGCTGCTAGCTATA
TGGCGAAACTGGCGCGGGCGAACGGCGGCTTTACCACCAGCAGCAGCGCGCCG
CTGATGCACGCGCAGGTACAGATTGTGGGCATACAaGATCCGTTGAATGCTCGCC
TGAGCCTGCTGCGCCGCAAGGATGAGATTATAGAGCTGGCGAACCGCAAAGATCA
GCTCTTAAACAGCTTAGGCGGCGGCTGCCGCGATATTGAGGTGCATACCTTTGCG
GACACCCCGCGGGGCCCGATGCTGGTGGCGCATCTGATTGTGGACGTACGCGAC
GCGATGGGCGCGAACACCGTGAATACCATGGCGGAAGCGGTAGCGCCGCTGATG
GAGGCGATTACCGGAGGCCAGGTACGCCTGCGCATACTGAGCAACCTGGCGGAT
CTGCGCCTGGCGAGGGCGCAGGTtCGGATAACACCGCAGCAACTGGAGACAGCG
GAGTTTTCAGGCGAAGCTGTGATTGAGGGCATTTTGGATGCGTATGCGTTTGCTGC
GGTtGATCCCTATCGCGCGGCGACCCATAACAAAGGCATTATGAATGGCATTGATC
CCCTGATTGTGGCGACAGGCAACGATTGGCGGGCTGTGGAGGCGGGCGCGCAC
GCGTACGCGTGCCGCTCAGGACATTATGGCAGCCTGACCACCTGGGAGAAAGATA
ACAACGGCCACCTTGTGGGCACCCTGGAGATGCCGATGCCAGTAGGCCTGGTGG

GCGGCGCGACCAAGACCCACCCGCTGGCGCAACTGAGCCTGCGCATTTTAGGCG
TGAAGACAGCGCAGGCGTTGGCTGAAATAGCGGTGGCGGTAGGCCTGGCGCAAA
ACTTAGGAGCGATGCGCGCGCTGGCGACCGAGGGCATTCAGCGCGGCCATATGG
CGCTGCACGCGCGCAATATAGCGGTGGTGGCGGGCGCGAGGGGCGACGAAGTG
GATTGGGTAGCGCGGCAGCTTGTGGAGTATCATGATGTGCGCGCGGATCGCGCG
GTAGCTCTGCTGAAGCAAAAACGCGGCCAATTAAGTCGAC

**315_pJL1-(CAT5aa)-HMGR_Spn**

TCTAGAAATAATTTTGTTTAACTTTAAGAAGGAGATATACATATGGAGAAAAAAATCA
AGATCAGCTGGAACGGTTTTAGCAAAAAAGCTACCAGGAGCGGCTTGAGCTCTT
GAAAGCGCAAGCTCTCTTAAGCCCAGAGCGGCAGGCGAGCCTTGAGAAGGATGA
ACAAATGAGCGTAACGGTAGCCGACCAACTTAGCGAGAACGTCGTAGGTACGTTT
AGCCTCCCATATTCACTTGTGCCaGAGGTCTTAGTCAATGGCCAAGAATACACTGT
GCCTTATGTAACGGAAGAACCtAGCGTAGTGGCTGCTGCATCATATGCATCAAAAA
TCATCAAGCGCGCCGGCGGCTTTACGGCCCAGGTCCATCAACGGCAAATGATTGG
GCAAGTCGCATTGTATCAGGTGGCGAACCCAAAATTGGCTCAGGAGAAGATTGCA
TCAAAGAAAGCTGAGCTCTTGGAGTTTGCAAACCAGGCATATCCAAGCATCGTGAA
ACGCGGTGGCGGGGCTCGCGATCTTCATGTCGAGCAAATcAAAGGGGAACCGGA
CTTTCTTGTGGTGTATATTCATGTCGATACTCAAGAAGCAATGGGCGCAAACATGC
TTAATACTATGCTTGAAGCATTAAAACCGGTCTTAGAAGAACTCAGCCAAGGTCAAA
GCCTTATcGGTATCCTCTCAAATTACGCTACTGATAGCCTTGTAACGGCCTCATGCC
GGATGGCATTTCGGTACTTGTCACGGCAGAAGGATCAGGGTCGGGAGATTGCTGA
GAAAATTGCTTTGGCGAGCCAATTTGCTCAAGCcGATCCATACCGGGCGGCGACG
CATAACAAAGGTATTTTTAACGGCATTGATGCTATTTTGATTGCAACGGGCAACGAC
TGGCGCGCAATCGAAGCGGGGGCACATGCATTTGCAAGCCGGGATGGTCGGTAT
CAGGGCTTAAGCTGTTGGACACTCGACTTGGAACGGGAAGAATTGGTCGGCGAGA
TGACTCTTCCtATGCCAGTCGCTACGAAGGGCGGGAGCATCGGGCTCAATCCGCG
CGTCGCGTTCAGCCATGATTTGTTAGGTAACCCAAGCGCACGGGAATTGGCACAA
ATTATCGTATCAATCGGCTTGGCACAGAACTTTGCCGCACTTAAAGCGTTAGTCAG
CACAGGAATCCAACAGGGGCACATGAAATTACAAGCAAAATCATTAGCGCTTTTAG
CGGGGGCGAGCGAAAGCGAAGTGGCGCCtTTAGTCGAGCGGCTCATCAGCGATAA
AACTTTTAATTTGGAGACGGCACAACGGTATCTCGAAAACTTACGGAGCTAAGTCG
AC

**316_pJL1-(CAT5aa)-HMGR_Bpe**

TCTAGAAATAATTTTGTTTAACTTTAAGAAGGAGATATACATATGGAGAAAAAAATC
AGCACCGATGCAAAAAATAGCCGTATTAGCGGCTTTCACAAAGATGATATTCCGAC
CCGTCTGGCACGTGTTGCAGCATTTGCAGGTCTGGATGATGAAACCGTTCAGCAT
CTGGCAAATATGGGTAATCTGGACCCGCAGCTGGCAGATCGTCTGATTGAAAATGT
TGTTGCAACCCTGAATGTGCCGATTGGTATTGCAACCAATATGAAAGTTGATGGCG

AAGATGTTCTGGTTCCGATGGCAACCGAAGAAAGCAGCGTTGTTGCAGCCGTTTG
TAATGCAGCACGTCAGTGTTATGATCAGGGTGGTTTTACCACCAGTATGAGCGGTA
GCCTGATGATTGCACAGGTTCAGCTGGTTGATGTTCCGGATGCAGCACATGCACG
TATGCGTATTCTGGAACATAAAGCCGAAGTTAAAGCACTGTGTGATGATTGTGATC
CGCTGCTGGTTAAACTGGGTGGTGGTCTGCAGGATGTTGAAGTTCGTATTGTTGAT
GCAGCCGGTGGTCCGATGGTTGTTACCCATCTGATTGTTGATACCCGTGATGCAAT
GGGTGCAAATGCAGTTAATAGCATGGCAGAAAAACTGGCACCGCATATTGAAAGCT
GGACCGGTGGTCGTGTTTATCTGCGCATTCTGAGCAATCTGGCCGATCGTCGCCT
GGCACGCGCACGTGCAGTTTGGACCTGTGATGCCATTGGTGGTGCAAGCGTTCGT
GATGGTATTATTAGCGCATATCGTTTTGCAGCAGCAGATCCGTATCGTGCAGCAAC
CCATAACAAAGGTATTATGAATGGTGTTAGCGCAGTTGTTCTGGCAACCGGTAATG
ATACACGTGCCGTTGAAGCCGGTGCACATGCATATGCCGCACGTAAAGGTTGGTA
TAGCAGCCTGACCGATTGGGAAGTTACCGCAGAAGGTCATCTGGCAGGCACCCTG
GAAATGCCGATGGCAGTTGGTCTGGTGGGTGGTGCCACAAAACTGCATCCGACCG
CACGTGCCTGTCTGAAAATTCTGGGTGTTAGCACCGCAGAACGGCTGGCACGCCT
GATTGCAGCAGTGGGTCTGGCACAGAATTTTAGCGCACTGAAAGCACTGGCAACC
ACCGGTATTCAGAAAGGTCATATGAGCCTGCATGCACAGAATATTGCAATGATGGC
AGGCGCAGTTGGTGATGAAATTGAACCGGTTGCAAAAGCCCTGGTTGCACAGGGT
GCAGTTCGTGTTGATGTTGCAGAAGCAGAACTGGCACGTCTGCGTGGTCAGGGTT
AAGTCGAC

**317_pJL1-(CAT5aa)-HMGR_Dac**

TCTAGAAATAATTTTGTTTAACTTTAAGAAGGAGATATACATATGGAGAAAAAAATC
GTCGCCGATTCCCGCCTGCCGAACTTCCGCGCCCTGACTCCGGCCCAGCGTCGT
GATTTTCTGGCAGATGCATGTGGTCTGAGTGATGCAGAACGTGCACTGCTGGCAG
CACCGGGTGCACTGCCTCTGGCACTGGCCGATGGTATGATTGAAAATGTTTTTGG
CTCATTTGAACTGCCGCTGGGTGTTGCAGGTAATTTTCGCGTTAATGGTCGTGATG
TGCTGGTTCCGATGGCAGTTGAAGAACCGAGCGTTGTTGCAGCAGCAAGCTATAT
GGCAAAACTGGCACGTGAAGATGGTGGTTTTCAGACCAGCAGCACCCTGCCGCTG
ATGCGTGCACAGGTTCAGGTTCTGGGTGTTACCGATCCGCATGGTGCACGTCTGG
CAGTTCTGCAGGCACGTGCACAGATTATTGAACGTGCAAATAGCCGTGATAAAGTG
CTGATTGGTCTGGGTGGTGGTTGTAAAGATATTGAAGTTCATGTGTTTCCGGATAC
ACCGCGTGGTCCGATGCTGGTTGTTCATCTGATTGTTGATGTTCGTGATGCAATGG
GTGCCAATACCGTTAATACCATGGCAGAAAGCGTTGCACCGCTGGTTGAGCAGAT
TACCGGTGGTAGCGTTCGTCTGCGTATTCTGAGCAATCTGGCCGATCTGCGTCTG
GCACGCGCACGTGTTCGTCTGACACCGCAGACCCTGGCAACCCAAGAACGTAGC
GGTGAAGAAATTATTGAAGGTGTTCTGGATGCATATACCTTTGCAGCAATTGATCC
GTATCGTGCAGCAACCCATAATAAAGGTATTATGAATGGTATCGATCCGGTTATTGT
TGCGACCGGTAATGATTGGCGTGCCGTTGAAGCCGGTGCACATGCCTATGCAAGC
CGTAGCGGTAGCTATACCAGCCTGACCCGTTGGGAAAAAGATGCCGGTGGTGCAC
TGGTTGGTAGCATCGAACTGCCGATGCCGGTTGGTCTGGTTGGCGGTGCCACCAA
AACCCATCCGCTGGCACGCCTGGCACTGAAAATTATGGATCTGCAGAGCGCACAG

CAGCTGGGTGAAATTGCAGCCGCAGTGGGTCTGGCACAGAATCTGGGTGCCCTG
CGTGCACTGGCAACCGAAGGTATTCAGCGTGGTCATATGGCACTGCATGCACGTA
ATATTGCCCTGGTTGCGGGTGCAACCGGTGATGAAGTTGATGCAGTTGCACGTCA
GCTGGCAGCCGAACATGATGTGCGTACCGATCGTGCCCTGGAAGTTCTGGCAGCA
CTGCGTGCCCGTGCATAAGTCGAC

**281_pJL1-(CAT5aa)-MK1_Sce**

TCTAGAAATAATTTTGTTTAACTTTAAGAAGGAGATATACATATGGAGAAAAAAATCA
GTTTACCCTTCTTAACGAGCGCCCCCGGGAAGGTGATCATCTTTGGCGAACACAG
CGCGGTCTACAATAAGCCAGCAGTCGCGGCGTCGGTCAGCGCTTTGCGCACTTAC
CTCTTAATCTCAGAGTCCAGCGCCCCGGATACGATCGAATTAGACTTCCCCGACAT
CTCATTTAACCATAAGTGGTCCATCAACGATTTCAACGCAATCACTGAGGATCAGG
TCAATTCCCAGAAATTGGCAAAGGCGCAGCAGGCAACTGATGGGTTAAGCCAAGA
ACTAGTGAGTTTATTAGATCCCTTATTGGCGCAGTTATCCGAATCCTTCCACTACCA
TGCCGCTTTTTGCTTCCTCTATATGTTTGTGTGTTTATGTCCTCATGCAAAGAACAT
CAAGTTTAGCTTGAAGAGCACGTTGCCTATCGGCGCGGGATTAGGGTCTTCAGCA
AGCATCAGCGTCAGTCTTGCATTGGCGATGGCATACTTAGGAGGATTAATCGGGA
GCAACGACTTGGAAAAGCTCAGTGAAAATGATAAGCATATCGTCAACCAGTGGGCA
TTCATCGGCGAAAAGTGCATCCACGGCACTCCATCTGGGATCGATAATGCGGTCG
CAACGTATGGCAACGCACTCTTGTTTGAAAAAGACTCCCATAACGGGACGATCAAT
ACGAATAACTTTAAGTTCTTAGATGATTTCCCGGCAATCCCGATGATCTTAACTTAT
ACGCGCATCCCGCGGAGCACGAAAGATTTAGTGGCGCGAGTGCGGGTCTTAGTC
ACTGAGAAATTTCCAGAAGTGATGAAGCCGATCTTAGATGCAATGGGCGAATGCG
CATTACAGGGGTTAGAGATCATGACGAAGTTGAGTAAATGCAAAGGGACTGATGAC
GAGGCGGTCGAAACGAACAACGAACTTTATGAACAGTTGTTGGAATTGATCCGCAT
CAACCATGGGCTCTTGGTTTCGATCGGCGTGAGCCATCCAGGGTTAGAATTAATCA
AAAACCTCTCAGATGATTTACGCATCGGGTCTACGAAATTAACTGGCGCGGGCGG
TGGGGGCTGTAGCTTGACGTTGTTACGACGCGACATCACGCAGGAGCAGATCGAC
TCATTCAAAAAGAAATTGCAGGATGATTTTAGTTACGAGACGTTTGAAACGGACTTG
GGCGGAACGGGGTGTTGCTTATTATCAGCCAAAAACTTAAACAAAGATTTAAAAAT
CAAATCGTTGGTCTTCCAGTTATTTGAAAACAAAACGACTACGAAGCAGCAGATCG
ACGATTTGTTATTACCGGGGAATACAAACTTACCGTGGACGTCTTGAGGATCCGCA
CTCGAGCACCACCACCACCACCACTGAGATCGTCGAC

**321_pJL1-(CAT7aa)-MK_Sau**

TCTAGAAATAATTTTGTTTAACTTTAAGAAGGAGATATACATATGCATATGGAGAAAA
AAATCACCCGTAAAGGTTATGGTGAAAGCACCGGTAAAATCATTCTGATTGGTGAA
CATGCAGTGACCTTTGGTGAACCGGCAATTGCAGTTCCGTTTAATGCAGGCAAAAT
CAAAGTTCTGATTGAAGCACTGGAAAGCGGTAACTATAGCAGCATTAAATCCGATG
TGTATGATGGCATGCTGTATGATGCACCGGATCATCTGAAAAGCCTGGTTAATCGT

TTTGTGGAACTGAACAACATTACCGAACCGCTGGCAGTTACCATTCAGACCAATCT
GCCTCCGAGCCGTGGTCTGGGTAGCAGCGCAGCAGTTGCAGTTGCATTTGTTCGT
GCAAGCTATGATTTTCTGGGTAAAAGCCTGACCAAAGAAGAACTGATTGAAAAAGC
AAATTGGGCAGAGCAGATTGCACATGGTAAACCGAGCGGTATTGATACCCAGACA
ATTGTTAGCGGTAAACCGGTGTGGTTTCAGAAAGGTCATGCAGAAACCCTGAAAAC
CCTGTCACTGGATGGTTATATGGTTGTGATTGATACCGGTGTTAAAGGTAGCACCC
GTCAGGCAGTTGAAGATGTTCATAAACTGTGTGAAGATCCGCAGTATATGAGCCAT
GTTAAACATATTGGTAAACTGGTTCTGCGTGCCAGTGATGTTATTGAACATCATAAT
TTTGAAGCCCTGGCCGATATCTTTAATGAATGTCATGCCGATCTGAAAGCACTGAC
CGTTAGCCATGATAAAATTGAGCAGCTGATGAAGATCGGCAAAGAAAATGGTGCCA
TTGCAGGTAAACTGACCGGTGCAGGTCGTGGTGGTAGCATGCTGCTGCTGGCAAA
AGACCTGCCGACCGCAAAAAACATTGTTAAAGCAGTGGAAAAAGCCGGTGCAGCA
CATACCTGGATTGAAAATTTAGGTGGTTAAGTCGACGTCGAC

**322_pJL1-(CAT7aa)-MK_Spn**

TCTAGAAATAATTTTGTTTAACTTTAAGAAGGAGATATACATATGCATATGGAGAAAA
AAATCACCAAAAAAGTTGGTGTTGGTCAGGCACATAGCAAAATTATCCTGATTGGT
GAACATGCCGTGGTTTATGGTTATCCGGCAATTAGCCTGCCGCTGCTGGAAGTTGA
AGTTACCTGTAAAGTTGTTAGCGCAGAAAGCCCGTGGCGTCTGTATGAAGAGGATA
CCCTGAGCATGGCAGTTTATGCAAGCCTGGAATATCTGGATATCACCGAAGCATGT
GTTCGTTGTGAAATTGATAGCGCAATTCCGGAAAAACGTGGTATGGGTAGCAGCG
CAGCAATTAGCATTGCAGCAATTCGTGCAGTGTTCGATTATTATCAGGCCGATCTG
CCGCATGATGTTCTGGAAATTCTGGTTAATCGTGCAGAAATGATTGCACACATGAA
TCCGAGCGGTCTGGATGCAAAAACCTGTCTGAGCGATCAGCCGATTCGTTTTATCA
AAAATGTGGGTTTTACCGAACTGGAAATGGATCTGAGCGCATATCTGGTTATTGCA
GATACCGGTGTTTATGGTCATACCCGTGAAGCAATTCAGGTTGTTCAGAATAAAGG
TAAAGATGCACTGCCGTTTCTGCATGCACTGGGTGAACTGACCCAGCAGGCAGAA
GTTGCCATTAGCCAGAAAGATGCAGAAGGTCTGGGTCAGATTCTGAGCCAGGCAC
ATCTGCATCTGAAAGAAATTGGTGTTAGCAGTCCGGAAGCAGATTTTCTGGTTGAA
ACCACACTGAGCCATGGTGCCCTGGGTGCAAAAATGAGCGGTGGTGGTTTAGGTG
GTTGTATTATTGCACTGGTTACCAATCTGACACATGCACAAGAACTGGCAGAACGT
CTGGAAGAAAAAGGTGCCGTTCAGACCTGGATTGAAAGCCTGTAAGTCGACGTCG
AC

**323_pJL1-(CAT7aa)-MK_Mma**

TCTAGAAATAATTTTGTTTAACTTTAAGAAGGAGATATACATATGCATATGGAGAAAA
AAATCGTTAGCTGTAGCGCACCGGGTAAAATCTACCTGTTTGGTGAACATGCAGTT
GTGTATGGTGAAACCGCAATTGCATGTGCAGTTGAACTGCGTACCCGTGTTCGTGC
AGAACTGAATGATAGCATTACCATTCAGAGCCAGATTGGTCGTACCGGTCTGGATT
TTGAAAAACATCCGTATGTTAGCGCAGTGATCGAAAAAATGCGTAAAAGCATTCCG

ATTAACGGTGTTTTTCTGACCGTTGATAGCGATATTCCGGTTGGTAGCGGTCTGGG
TAGCAGCGCAGCAGTTACCATTGCAAGCATTGGTGCACTGAATGAACTGTTTGGTT
TTGGTCTGAGCCTGCAAGAAATTGCAAAACTGGGTCATGAAATCGAGATTAAAGTT
CAGGGTGCAGCAAGCCCGACCGATACCTATGTTAGCACCTTTGGTGGTGTTGTTA
CCATTCCGGAACGTCGTAAACTGAAAACACCGGATTGTGGTATTGTTATTGGTGAT
ACCGGTGTGTTTAGCAGCACCAAAGAACTGGTTGCAAATGTTCGTCAGCTGCGTG
AAAGCTATCCGGATCTGATTGAACCGCTGATGACCAGCATTGGTAAAATTAGTCGT
ATTGGCGAACAGCTGGTTCTGAGCGGTGATTATGCGAGCATTGGTCGTCTGATGA
ATGTTAATCAGGGTCTGCTGGATGCACTGGGTGTTAATATTCTGGAACTGAGCCAG
CTGATTTATAGCGCACGTGCAGCCGGTGCATTTGGTGCAAAAATTACCGGTGCCG
GTGGTGGTGGTTGTATGGTTGCACTGACCGCACCGGAAAAATGTAATCAGGTTGC
CGAAGCAGTTGCCGGTGCAGGCGGTAAAGTGACCATTACCAAACCGACCGAACAG
GGTCTGAAAGTTGATTAAGTCGACGTCGAC

**324_pJL1-(CAT7aa)-MK_Pze**

TCTAGAAATAATTTTGTTTAACTTTAAGAAGGAGATATACATATGCATATGGAGAAAA
AAATCAGCACCGGTCGTCCGGAAGCGGGTGCACATGCACCGGGTAAACTGATTCT
GAGCGGTGAACATAGCGTTCTGTATGGTGCACCGGCACTGGCAATGGCAATTGCA
CGTTATACCGAAGTTTGGTTTACCCCGTTAGGTATTGGTGAAGGTATTCGTACCAC
CTTTGCAAATCTGAGTGGTGGTGCAACCTATAGCCTGAAACTGCTGAGCGGTTTTA
AAAGCCGTCTGGATCGTCGTTTTGAACAGTTTCTGAATGGTGATCTGAAAGTGCAT
AAAGTTCTGACCCATCCTGATGATCTGGCAGTTTATGCCCTGGCAAGCCTGCTGCA
TGATAAACCGCCTGGCACCGCAGCAATGCCTGGTATTGGCGCAATGCATCATCTG
CCTCGTCCGGGTGAACTGGGTAGCCGTACCGAACTGCCGATTGGTGCCGGTATG
GGTAGCAGCGCAGCAATTGTTGCAGCAACCACCGTTCTGTTTGAAACCCTGCTGG
ATCGCCCTAAAACACCGGAACAGCGTTTTGATCGTGTTCGTTTTTGTGAACGTCTG
AAACATGGTAAAGCAGGTCCGATTGATGCAGCAAGCGTTGTTCGTGGTGGTCTGG
TTCGTGTTGGTGGTAATGGTCCGGGTAGCATTAGCTCATTTGATCTGCCTGAAGAT
CACGATCTGGTTGCAGGTCGTGGTTGGTATTGGGTTCTGCATGGCCGTCCGGTTA
GCGGCACCGGTGAATGTGTTAGCGCAGTTGCAGCAGCACATGGTCGTGATGCAG
CCCTGTGGGATGCATTTGCAGTTTGTACCCGTGCACTGGAAGCAGCACTGCTGTC
AGGTGGTAGTCCGGATGCAGCCATTACCGAAAATCAGCGTCTGCTGGAACGTATT
GGTGTTGTTCCGGCAGCAACCCAGGCACTGGTTGCACAGATTGAAGAAGCAGGCG
GAGCAGCAAAAATTTGTGGTGCAGGTAGCGTGCGTGGTGATCATGGTGGTGCCGT
TCTGGTGCGTATTGATGATGCACAGGCCATGGCAAGCGTTATGGCACGTCATCCG
GATCTGGATTGGGCACCGCTGCGTATGAGTCGTACCGGTGCAGCACCTGGTCCG
GCACCGCGTGCACAGCCGTTACCTGGTCAGGGTTAAGTCGACGTCGAC

**325_pJL1-(CAT7aa)-MK_Hme**

TCTAGAAATAATTTTGTTTAACTTTAAGAAGGAGATATACATATGCATATGGAGAAAA
AAATCACCGTTAGCAGCGCACCGGGTAAAGTTTACCTGTTTGGTGAACATGCAGTT
GTTTATGGTGAACCGGCAGTTCCGTGTGCAGTTGAACGTCGTGCAACCGTTAGCG
TTAGCGCACGTGATGATGATCATGTTCGTGTTCGTGCAGAGGATCTGAGCCTGAAT
GGTTTTACCGTTGAATATAGCGGTAGCACCGGTAATCATCCTGATGTTGATGTTCC
GACACCGCTGGTTGAAGCAGCAATGGGTTATATTGATGCAGCAGTTGCACAGGCT
CGTGATGCAGCCGATGCACCGGATGCAGGTTTTGATATTACCGTTAAAAGCGATAT
TCCGCTTGGTGCAGGTCTGGGTAGCAGTGCAGCCGTTGTTGTTGCAGGTATTGAT
GCGGCAACCCGTGAACTGGGTGTTGAACTGAGTCCGCGTGAAATTGCAGATCGTG
CATATCGTGCAGAACATGAAGTTCAGGATGGTCAGGCAAGCCGTGCAGATACCTTT
TGTAGCGCAATGGGTGGTGCAGTTCGTGTTGAAGGTGATGATTGTCGTACCATTGA
TGCACCGCCTCTGCCGTTTGTTATTGGTTTTGATGGTGGTGCCGGTGATACCGGTG
CACTGGTTAGCGGTGTGCGTGCACTGCGTGAAGAATATGATTTTGCAGCAGATAC
CGTGAGCACCATTGGTGATATTGTTCGTCGTGGTGAGGATCTGCTGGCAGATGCA
GATCCGGAAGAACCGAGCGAAGCACTGCTGAGCGAACTGGGTCGTTTTATGAATT
TTAACCATGGTCTGCTGGAAGCCCTGGGTGTTAGCAGCCGTAGCCTGGATAGCAT
GGTTTGGGCAGCACGTGAAGCCGGTGCCTATGGTGCAAAACTGACCGGTGCAGG
CGGTGGTGGTTGTATTGTTGCACTGGACCCGACACCGGAAACACAGACCGCACTG
CGTTTTACACCGGGTTGTGAAGATGCATTTCGTGCCGAACTGGCAACCGAAGGTG
TTCGTGTGGAAGAACCTCCGGCAAGCGCAGCCAGCGCAGAAAGCAATGTTGGTGA
TGATCAGAGTCCGGAAGGTAGCGCATAAGTCGACGTCGAC

**326_pJL1-(CAT7aa)-MK_Nma**

TCTAGAAATAATTTTGTTTAACTTTAAGAAGGAGATATACATATGCATATGGAGAAAA
AAATCAAAAGCAAAGCAAGCGCACCGGGTAAAGTTATTCTGTTTGGTGAACATTTT
GTGGTGTATGGCGTTAAAGCAATTCTGTGCGCAATTAACAAACGTATTGCAGTGAC
CGCAGAAAAAATCGATGAACGCAAAATCAGCATCAAAAGCAATATTGGTCATCTGG
AACTGGAACCGAATAAACCGATTAGCGAAATTAATAGTCCGCTGAAACCGTTCTAT
TATCTGGCCAATAAAATCATCCAGGACAAGAACTTTGGCATCAAAATTGATGTGGA
AAGCGAAATTCCGTTAGGTGTTGGTCTGGGTAGCAGCAGCGCATGTTGTGTTGCC
GGTGCAGCAGCAATTAGCAACCTGTTTGAAAATAACAGCAAAGAAGAGATCCTGAA
ACTGGCAATTGAAGCCGAAAAAACCATTTTTCAGAATACCAGCGGTGCAGATTGTA
CCGTTTGTACCTTTGGTGGTCTGATGGAATATGATAAAGAAAACGGCTTCAGCAAA
ATCGAAAGCGAACCGAATTTTCATCTGGTGATTGCCAATAGCAATGTGGAACATAG
CACCGAAAGCGTTGTTGCGGGTGTTCGTAAATTCAAAAAAAACAACGAAGCCGAGT
TCAGCAAACTGTGTAAAGATGAAAGCCATCTGATTGAGAATGTGCTGGAACTGCTG
AAAGAAAACAATATTCGTGAACTGGGTGAACGCGTGATCAAAAATCAAGAATATCT
GGAACGCATTGGCATCAGCAATGCAAAACTGCGTGAAATGATTCAGACAGGTCAG
AATAGCAGCTTTGGTGCAAAAATTACCGGTGCCGGTGGTGGTGGTTGTATTTTTGC
CCTGACCGATGAAAGTAATCTGGAAAACACCATCAAAGAATTTAAAGAAAAAAACC
ACGAGTGCTTTAGCGTGAAAATCGACTTTAAAGGTCTGGACACCTTTTAAGTCGAC
GTCGAC

**327_pJL1-(CAT7aa)-MK_Mxa**

TCTAGAAATAATTTTGTTTAACTTTAAGAAGGAGATATACATATGCATATGGAGAAAA
AAATCGCACCGCGTCCGGAAAGCCTGAGCGCATTTGGTGCAGGTAAAGTTATTCT
GCTGGGTGAACATAGCGTTGTTTATGGTCATCCGGCACTGGCAGGTCCGCTGAGC
CAGGGTGTTACCGCACGTGCAGTTCCGGCAAAAGCATGTCAGCTGGCACTGCCGA
GCACACTGAGCCGTCCGCAGCGTGCACAGCTGACCGCAGCATTTGCCCGTGCAG
CCGAAGTTACCGGTGCACCTCCGGTTAAAGTTAGCCTGGAAGCCGATCTGCCGCT
GGCAGTTGGTCTGGGTAGCAGCGCAGCACTGAGCGTTGCATGTGCACGTCTGCTG
CTGCAGGCAGCCGGTAAAGTTCCGACACCGAAAGATGCAGCACGTGTTGCCTGG
GCAATGGAACAAGAATTTCATGGCACCCCGAGCGGTGTTGATCATACCACCAGTG
CAGCAGAACAGCTGGTGCTGTATTGGCGTAAACCGGGTGCAGCAAAAGGCACCG
GTCAGGTTGTTGAAAGTCCGCGTCCGCTGCATGTTGTTGTTACCCTGGCAGGCGA
ACGTAGCCCGACCAAAAAAACCGTTGGTGCACTGCGTGAACGTCAGGCACGTTGG
CCGAGCCGTTATGAACGTCTGTTTGCAGAAATTGGTCGTGTTAGCAGCGAAGGTG
CAAAAGCAGTTGCAGCCGGTGATCTGGAAGCACTGGGTGATGCAATGAATGTTAA
TCAGGGTCTGCTGGCAGCCCTGGGTCTGAGCAGCCCTCCGCTGGAAGAAATGGTT
TATCGTCTGCGCGAACTGGGTGCCCTGGGTGCAAAACTGACAGGTGCCGGTGGT
GATGGTGGTGCAGTTATTGGTCTGTTTCTGGAACCGAAACCGGTTGTTACCAAACT
GACCCGTATGGGTGTTCGTTGTTTTAGCTCACAGCTGGCTGGTCCGCGTGCAAGC
TAAGTCGACGTCGAC

**328_pJL1-(CAT7aa)-MK_Bmo**

TCTAGAAATAATTTTGTTTAACTTTAAGAAGGAGATATACATATGCATATGGAGAAAA
AAATCGAAGTTCGTGCCCGTGCACCGGGTAAAATCATTCTGGCAGGCGAACATGC
AGTTGTTCATGGTAGCACCGCAGTTGCAGCAGCAATTGATCTGTATACCTATATCA
GCCTGCATTTTCCGACACCGGCAGAAAATGATGATGCACTGAAACTGCATCTGAAA
GATATGGGCTTAGAATTTAGCTGGCCTGTGGGTCGTATTAAAGATGTTCTGCCGGA
AGTTAGCAGCCATGATGTGAGCAGCCCGAGCAGCTGTAGCCTGGAAACCCTGAAA
GCAATTGCAGCACTGGTTGAAGAACAGAATATTCCGGAAGCAAATGTTGGTCTGGC
AAGCGGTGTTAGCACCTTTCTGTGGATGTATAGCAGCATTCATGGTTACAAACCGG
CAAAAGTTGTTGTTACCAGCGAACTGCCGTTAGGTAGCGGTCTGGGTAGCAGCGC
AGCATTTTGTGTTAGCCTGAGCGCAGCACTGCTGGCACTGAGCGATAGCGTTAAA
CTGGATTTTAGCAATCAAGGCTGGCAGATGTTTGCAGAAACCGAACTGGAACTGGT
GAATAAATGGGCATTTGAAGGCGAAAAAATCATCCATGGTAAACCGAGCGGTATTG
ATAATACAGTGAGCACCTATGGCAACATGATCAAATTCAAAAGCGGTGAAATGGTG
CGCATCAAAACCAATATGCCGCTGAAAATGCTGATCACCAATACCAAAGTTGGCCG
TAATACAAAAGCCCTGGTTGCGGGTGTTAGCGAACGTACCGTTCGTCATAGCAATG
CAATGAGCAGCGTTTTTAATGCCGTTGATTGCATTAGCAATGAACTGGCAGCAATT
ATTCAGAGTCCGGTTAGTGATGATCTGGCCATTACCGAAAAAGAAGAAAAACTGGG

CGAACTGATGGAAATGAATCAGGGTCTGCTGCAGTGTATGGGTGTGAGCCATGCA
AGCATTGAAACCGTTATTCGTACCACGCTGAAATACAAACTGGCAACCAAACTGAC
CGGTGCCGGTGGTGGTGGTTGTGTTCTGAGCCTGCTGCCGACACTGCTGAGCGG
CACCGTTGTTGATATTGTTATTAGTGAACTGGAAGCCTGTGGTTTTCAGTGTCTGAT
TGCAGGTATTGGCGGTAATGGTGTTGAAATTAGCTTTAGCCCGAGCTAAGTCGACG
TCGAC

**282_pJL1-(CAT5aa)-PMK1_Sce**

TCTAGAAATAATTTTGTTTAACTTTAAGAAGGAGATATACATATGGAGAAAAAAATCA
GTGAGTTGCGCGCATTCTCGGCACCGGGAAAAGCTTTATTAGCGGGCGGGTATTT
AGTGTTGGATACGAAATATGAAGCCTTTGTCGTCGGGTTATCAGCCCGCATGCATG
CGGTCGCACATCCATACGGCTCTTTACAGGGAAGCGATAAGTTTGAAGTCCGGGT
CAAATCTAAACAGTTTAAAGATGGAGAGTGGCTTTACCATATCAGTCCAAAAAGTG
GGTTCATCCCAGTGTCAATCGGTGGGAGCAAGAATCCATTCATCGAAAAAGTGATC
GCGAATGTCTTTTCATACTTTAAACCAAATATGGACGACTACTGTAACCGCAATTTA
TTCGTGATCGATATCTTCAGCGATGATGCATACCATAGCCAAGAGGATTCAGTGAC
TGAACATCGGGGGAATCGCCGCTTGTCTTTTCATTCACACCGCATCGAAGAAGTGC
CTAAAACGGGACTTGGGTCGTCAGCCGGCTTGGTCACGGTGTTGACGACGGCGTT
AGCATCCTTTTTTGTCTCAGACCTCGAAAACAACGTCGACAAATATCGCGAAGTGA
TCCATAACTTGGCCCAGGTGGCGCATTGCCAGGCGCAAGGCAAAATCGGGTCAG
GATTTGATGTCGCTGCTGCCGCCTATGGGAGCATCCGCTATCGCCGCTTCCCGCC
TGCCTTGATCAGCAACTTGCCGGATATCGGGTCGGCGACGTACGGGTCTAAACTT
GCTCATTTAGTGGATGAAGAAGACTGGAACATCACAATCAAATCGAATCATTTACCA
TCAGGGTTGACGTTATGGATGGGGGATATCAAGAACGGCAGTGAAACGGTCAAAC
TTGTCCAAAAGGTCAAAAACTGGTATGATTCACATATGCCGGAATCATTAAAAATCT
ATACGGAACTAGATCATGCCAACAGCCGCTTTATGGATGGGTTGAGCAAATTAGAT
CGATTGCACGAGACGCATGACGATTACTCAGATCAAATCTTTGAGAGCTTGGAGCG
GAACGACTGCACTTGCCAGAAGTATCCAGAAATCACGGAAGTGCGCGATGCCGTG
GCAACGATCCGCCGGTCGTTTCGCAAAATCACGAAAGAAAGCGGCGCAGATATCG
AACCACCTGTCCAGACGTCATTATTGGATGATTGTCAAACTTTGAAAGGGGTGTTG
ACGTGTTTGATCCCAGGCGCGGGCGGCTATGACGCAATCGCCGTCATCACGAAGC
AGGATGTGGATTTGCGGGCGCAGACTGCGAACGACAAACGCTTTAGCAAGGTGCA
GTGGCTCGATGTCACGCAAGCGGACTGGGGCGTGCGGAAAGAAAAAGATCCCGA
AACGTATTTGGATAAATGAGGATCCTGACTCGAGCACCACCACCACCACCACTGAG
ATCGTCGAC

**329_pJL1-(CAT7aa)-PMK_Sau**

TCTAGAAATAATTTTGTTTAACTTTAAGAAGGAGATATACATATGCATATGGAGAAAA
AAATCATCCAGGTTAAAGCACCGGGTAAACTGTATATTGCCGGTGAATATGCAGTT
ACCGAACCGGGTTATAAAAGCGTTCTGATTGCACTGGATCGTTTTGTTACCGCAAC

CATTGAAGAAGCCGATCAGTATAAAGGCACCATTCATAGCAAAGCCCTGCATCACA
ATCCGGTTACCTTTAGCCGTGATGAAGATAGCATTGTTATTAGCGATCCGCATGCA
GCAAAACAGCTGAATTATGTTGTGACCGCCATCGAAATCTTTGAGCAGTATGCAAA
AAGCTGCGACATTGCCATGAAACATTTTCATCTGACCATCGATAGCAACCTGGATG
ATAGCAATGGTCATAAATATGGTCTGGGTAGCAGCGCAGCAGTTCTGGTTAGCGTT
ATTAAAGTGCTGAACGAGTTCTACGATATGAAACTGAGCAACCTGTACATCTATAAA
CTGGCCGTTATTGCCAACATGAAACTGCAGAGCCTGAGCAGCTGTGGTGATATTG
CAGTTAGCGTTTATAGCGGTTGGCTGGCATATAGCACCTTTGATCATGAATGGGTG
AAACACCAGATTGAAGATACCACCGTTGAAGAAGTGCTGATTAAAAACTGGCCTGG
TCTGCATATTGAACCGCTGCAGGCACCGGAAAATATGGAAGTTCTGATCGGTTGGA
CCGGTAGTCCGGCAAGCAGTCCGCATTTTGTTAGCGAAGTTAAACGTCTGAAAAG
CGATCCGAGCTTTTATGGTGATTTTCTGGAAGATAGCCATCGGTGTGTTGAAAAAC
TGATCCATGCCTTTAAAACCAACAACATTAAAGGCGTGCAGAAAATGGTTCGTCAG
AATCGTACCATTATTCAGCGCATGGATAAAGAAGCAACCGTTGATATTGAAACCGA
GAAACTGAAATACCTGTGCGATATTGCCGAAAAATATCATGGTGCAAGCAAAACCA
GCGGTGCCGGTGGTGGTGATTGTGGTATTACCATTATCAACAAAGATGTGGACAAA
GAGAAAATCTACGACGAATGGACCAAACATGGTATCAAACCGCTGAAATTCAACAT
CTATCATGGCCAGTAAGTCGACGTCGAC

**330_pJL1-(CAT7aa)-PMK_Spn**

TCTAGAAATAATTTTGTTTAACTTTAAGAAGGAGATATACATATGCATATGGAGAAAA
AAATCATTGCCGTTAAAACCTGCGGTAAACTGTATTGGGCAGGCGAATATGCAATT
CTGGAACCGGGTCAGCTGGCACTGATTAAAGATATTCCGATTTATATGCGTGCCGA
AATCGCATTTAGCGATAGCTACCGTATTTATAGCGATATGTTTGATTTTGCCGTTGA
CCTGCGTCCGAATCCTGATTATAGCCTGATTCAAGAAACCATTGCACTGATGGGTG
ATTTTCTGGCAGTTCGTGGTCAGAATCTGCGTCCGTTTAGCCTGGCCATTTATGGT
AAAATGGAACGCGAAGGCAAAAAATTCGGTCTGGGTAGCAGCGGTAGCGTTGTTG
TTCTGGTTGTTAAAGCACTGCTGGCCCTGTATAATCTGAGCGTTGATCAGAACCTG
CTGTTTAAACTGACCAGCGCAGTTCTGCTGAAACGTGGTGATAATGGTAGCATGGG
TGATCTGGCATGTATTGCAGCAGAGGATCTGGTTCTGTATCAGAGCTTTGATCGTC
AGAAAGTTGCAGCATGGCTGGAAGAAGAAAATCTGGCAACCGTTCTGGAACGTGA
TTGGGGTTTTAGCATTAGCCAGGTTAAACCGACACTGGAATGTGACTTTCTGGTTG
GTTGGACCAAAGAAGTTGCAGTTAGCAGCCACATGGTTCAGCAGATTAAACAGAAT
ATTAACCAGAACTTCCTGACCAGCAGCAAAGAAACCGTTGTTAGCCTGGTTGAAGC
ACTGGAACAGGGTAAAAGCGAAAAAATCATTGAACAGGTTGAAGTGGCAAGCAAA
CTGCTGGAAGGTCTGAGCACCGATATCTATACACCGCTGCTGCGTCAGCTGAAAG
AAGCAAGCCAGGATCTGCAGGCAGTTGCAAAAAGCAGTGGTGCCGGTGGTGGTG
ATTGTGGTATTGCACTGTCATTTGATGCACAGAGCACCAAAACACTGAAAAATCGTT
GGGCTGATCTGGGTATTGAACTGCTGTATCAAGAACGTATTGGCCACGATGATAAA
AGCTAAGTCGACGTCGAC

**331_pJL1-(CAT7aa)-PMK_Efa**

TCTAGAAATAATTTTGTTTAACTTTAAGAAGGAGATATACATATGCATATGGAGAAAA
AAATCATCGAAGTTACCACACCGGGTAAACTGTTTATTGCCGGTGAATATGCAGTT
GTTGAACCGGGTCATCCGGCAATTATTGTTGCAGTTGATCAGTTTGTTACCGTGAC
CGTTGAAGAAACCACCGATGAAGGTAGCATTCAGAGCGCACAGTATAGCAGCCTG
CCGATTCGTTGGACCCGTCGTAATGGTGAACTGGTTCTGGATATTCGTGAAAACCC
GTTTCATTATGTTCTGGCAGCAATTCATCTGACCGAAAAATATGCACAAGAGCAGA
ATAAAGAGCTGAGCTTCTATCATCTGAAAGTTACCAGCGAACTGGATAGCAGCAAT
GGTCGTAAATATGGTCTGGGTAGCAGCGGTGCAGTTACCGTTGGCACCGTTAAAG
CACTGAATATCTTTTATGATCTGGGCCTTGAAAACGAGGAAATCTTTAAACTGAGCG
CACTGGCACATCTGGCAGTTCAAGGTAATGGTAGCTGTGGTGATATTGCAGCAAG
CTGTTATGGTGGTTGGATTGCATTTAGCACCTTTGATCATGATTGGGTGAATCAGAA
AGTTGCAACCGAAACACTGACCGATCTGCTGGCAATGGATTGGCCTGAACTGATG
ATTTTTCCGCTGAAAGTTCCGAAACAGCTGCGTCTGCTGATTGGCTGGACCGGTAG
TCCGGCAAGCACCAGCGATCTGGTTGATCGTGTTCATCAGAGCAAAGAAGAAAAA
CAGGCAGCCTATGAACAGTTCCTGATGAAAAGCCGTCTGTGTGTTGAAACCATGAT
CAATGGCTTTAACACCGGTAAAATTAGCGTGATTCAGAAGCAGATTACCAAAAATC
GTCAGCTGCTGGCAGAACTGAGCAGCCTGACCGGTGTTGTTATTGAAACCGAAGC
GCTGAAAAATCTGTGTGATCTGGCAGAAAGCTATACCGGTGCAGCAAAAAGCAGT
GGTGCCGGTGGTGGTGATTGTGGTATTGTGATTTTTCGCCAGAAAAGCGGTATTCT
GCCGCTGATGACCGCATGGGAAAAAGATGGTATTACACCGCTGCCGCTGCATGTT
TATACCTATGGTCAGAAAGAATGCAAAGAGAAACACGAAAGCAAACGTTAAGTCGA
CGTCGAC

**332_pJL1-(CAT7aa)-PMK_Pze**

TCTAGAAATAATTTTGTTTAACTTTAAGAAGGAGATATACATATGCATATGGAGAAAA
AAATCGATCAGGTTATTCGTGCAAGCGCACCGGGTAGCGTTATGATTACCGGTGAA
CATGCAGTTGTTTATGGTCATCGTGCAATTGTTGCAGGTATTGAACAGCGTGCACA
TGTTACCATTGTTCCGCGTGCAGATCGTATGTTTCGTATTACCAGCCAGATTGGTG
CACCGCAGCAGGGTAGCCTGGATGATCTGCCTGCCGGTGGCACCTATCGTTTTGT
TCTGGCAGCAATTGCCCGTCATGCACCGGATCTGCCGTGTGGTTTTGATATGGATA
TTACCAGTGGTATTGATCCGCGTTTAGGTCTGGGTAGCAGCGCAGCAGTTACCGTT
GCATGTCTGGGTGCACTGAGCCGTCTGGCAGGTCGTGGCACCGAAGGTCTGCAT
GATGATGCACTGCGTATTGTTCGTGCCATTCAAGGTCGTGGTAGCGGTGCCGATC
TGGCAGCCAGCCTGCATGGTGGTTTTGTTGCATATCGCGCACCGGATGGTGGTGC
AGCACAGATTGAAGCACTGCCGGTTCCGCCTGGTCCGTTTGGTCTGCGTTATGCA
GGTTATAAAACCCCGACCGCAGAAGTGCTGCGTCTGGTTGCCGATCGTATGGCAG
GTAATGAAGCAGCATTTGATGCCCTGTATAGCCGTATGGGTGCCAGCGCAGATGC
AGCAATTCGTGCAGCCCAAGGTCTGGATTGGGCAGCATTTCATGATGCGCTGAAT
GAATATCAGCGTCTGATGGAACAGCTGGGTGTTAGTGATGATACCCTGGATGCAAT
TATTCGCGAAGCACGTGATGCCGGTGCAGCAGTTGCAAAAATTTCAGGTAGCGGT

CTGGGTGATTGTGTTCTGGCCCTGGGTGATCAGCCGAAAGGCTTTGTTCCGGCAA
GCATTGCAGAAAAAGGTCTGGTTTTTGATGATTAAGTCGACGTCGAC

**333_pJL1-(CAT7aa)-PMK_Tha**

TCTAGAAATAATTTTGTTTAACTTTAAGAAGGAGATATACATATGCATATGGAGAAAA
AAATCATCGAAGTTAGCACACCGGGTAAACTGTATATTGCCGGTGAATATGCAGTT
GTTGAACCGGGTCATCTGGCAATTATTGCAGCAGTTGATCAGTTTATCAACGTGAC
CATTGAAAGCGCAACCGAAAATGGTAGCATTCAGAGCCAGCAGTATAGCGATCTG
CCGATTCGTTGGACCCGTCGTGAAGGTGAACTGGTTCTGGATCATCGTGAAAATCC
GTTTCATTATATTCTGGCAGCAATTCGTCTGACCGAACGTTATGCAAAAGAACAGG
GCACCCTGCTGAGCTTTTATCATCTGAAAGTTACCAGCGAACTGGATAATAGCAGC
GGTCGTAAATATGGTCTGGGTAGCAGTGGTGCAGTTACCGTTGGCACCGTTAAAG
CACTGAACCTGTTTTATGATCTGCAAATGGACCCGCTGATGCAGTTTAAAATCGCA
GCACTGGCACATCTGGCAGTTCAAGGTAATGGTAGCTGTGGTGATATTGCAGCCA
GCTGTTTTGGTGGTTGGCTGGCATTTAGCACCTTTGATCATCAGTGGGTTAAAAAA
CGTCAAGAAACCTGGAAAATCAGCGACCTGCTGAAAAGCGATTGGCCGAAACTGA
GCATTCAGCCGCTGCAGAGCCCGAAAAATATGCGTCTGCTGATTGGCTGGACCGG
TAGTCCGGCAAGCACCAGCGATCTGGTTGATCAGGTTAATCAGAGCAAAGAGGAT
AAAGACGACATCCAGAAAAACTATGAACAGTTTCTGACCGATAGCCATCATTGTGT
TGAGGATCTGATGGATGGTTTCGTTAAAGATGATGTGACCAAAATCAAGAAAATGA
TCCGCAAAAATCGTACCCTGCTGCAGAATCTGGCAAAAGCAACCAATGTTGTTATT
GAAACACCGGCACTGAAACAGCTGTGTGATCTGGCAGAAAATTGTGGTGGTGCAG
CAAAAAGCAGCGGTGCCGGTGGTGGTGATTGTGGTATTGTTATTGCCGATCAGAA
AACCGGTATCCTGCCGCTGATGAGCAAATGGGAAAAAGCAGATATTATTCCGCTGC
CGCTGCATGTTTATCATTATCGTGGTGGTCCGAAATAAGTCGACGTCGAC

**334_pJL1-(CAT7aa)-PMK_Eco**

TCTAGAAATAATTTTGTTTAACTTTAAGAAGGAGATATACATATGCATATGGAGAAAA
AAATCCTGAAGAAAATCAAACATGATCCGACGCTGAATAGCCAAGGTCAGGGTTTT
GCACCGGGTAAACTGTATCTGGCAGGCGAATATGCAGTTCTGGCACCGCGTCAGC
CTGCAATTCTGCTGGCACTGAATCGTTATGTTAAAGTGACCATTAAACCGAGCAGC
ACCCTGAATCAGGGCATTCTGAGCCAGGCAAAAGGTCAGGCAGATTATCATTATCA
GCGTCAGAATGGTAGCATTCCGCAGAAAGAAGCATATTGGACCTATTGTCTGGCA
GCCATTCAGATTGTTGAAGTTCTGTTTCGTCAGAAAGGCCAGGTTATTGCCGATTAT
CATCTGGAAACCATGAGCGATCTGGTTGAAGAAGTTAGCGGCAAAAAATTCGGTCT
GGGTAGCAGCGGTGCAATTACCGTTGCAACCATTCGTGCACTGCTGGATTTTTATG
GTTATCAGGCCGATAGTCCGCTGGATGTGTATAAACTGGCCGTTCTGGCCCTGGTT
AATCTGGGTAATAATGGTAGCTTTGGTGATCTGGCAGCAGCAGCATTTGGTGGTTG
GGTTTATTATCAGGCACCGGATCGTCAGTGGCTGGCAGATCAGGTTAGCCAGAAT
CAGACCATTGATTTCTTTCTGGAAAATAGCTGGCCGAACCTGCAGATTGAAAGCCT

GCCGGTTCCGAGCAAAATTGATTTCCTGGTTGCATGGACCCAGAGTCCGGCAAGC
AGCGATCATTTTGTTGCAAACTTTAAAGAAGCCAGCCAGCAAGAACCGCAGCGTTA
TCAAGAATTTCTGGCCGAAAATAAAGATGCAGTGCTGGCCCTGAAAACCGCACTGA
TCCAGGATGATGTTGGTCAGAGCCAGAGCCTGATTAACAAAATTGGTCAGCAGCT
GGATAATCTGAGCCATCATCTGAAACTGGGTATTCTGACACCGCAGCTGGAAACAA
TGATTAGCCTGGCACAGGTGTATGGTTATGCAGCAAAAAGCAGTGGTGCCGGTGG
TGGTGATTGTGGTATTGCATTAGGTGGTCTGGAAGCACGTAAAACCGATCTGATTC
GTGCATGGGAAAAACAAGAAATCACCTATCTGGATCTGCAAATCAGCCAGACCTTT
TAAGTCGACGTCGAC

**283_pJL1-(CAT5aa)-PMD1_Sce**

TCTAGAAATAATTTTGTTTAACTTTAAGAAGGAGATATACATATGGAGAAAAAAATCA
CTGTGTACACGGCCTCTGTGACTGCCCCTGTCAATATCGCCACTTTGAAGTATTGG
GGAAAACGGGACACAAAGTTGAACCTTCCTACTAACTCATCTATCTCGGTCACGTT
ATCACAGGATGACCTACGCACATTAACTAGCGCTGCGACGGCCCCAGAGTTTGAA
CGAGACACGTTATGGTTAAACGGGGAACCGCACTCAATCGACAACGAACGCACGC
AGAACTGCCTTCGAGACTTGCGACAGTTACGCAAGGAAATGGAATCAAAGGACGC
AAGTTTGCCTACGTTAAGCCAGTGGAAACTACACATCGTCTCCGAAAACAATTTTC
CAACGGCCGCGGGCTTGGCGTCTTCTGCGGCGGGGTTTGCGGCCTTGGTCAGCG
CCATCGCGAAGTTGTACCAGTTGCCGCAATCCACGTCCGAAATCAGCCGCATCGC
CCGCAAGGGAAGCGGCTCTGCGTGCCGCTCATTGTTTGGTGGGTACGTCGCATG
GGAAATGGGGAAAGCGGAAGATGGCCATGATTCCATGGCCGTCCAGATCGCCGA
CTCAAGCGACTGGCCACAAATGAAAGCGTGCGTCTTAGTGGTTTCAGATATCAAAA
AGGATGTCTCCTCTACGCAAGGCATGCAGTTGACTGTCGCCACTTCTGAATTATTT
AAAGAACGCATCGAACATGTCGTCCCGAAGCGCTTTGAAGTCATGCGGAAAGCAA
TCGTGGAAAAAGATTTCGCAACTTTTGCCAAGGAAACGATGATGGATTCTAATAGC
TTCCATGCAACGTGCTTAGACAGCTTCCCACCGATCTTCTACATGAACGACACGAG
TAAGCGGATCATCTCTTGGTGTCACACTATCAACCAATTTTACGGGGAAACGATCG
TGGCCTACACATTTGATGCCGGCCCGAACGCGGTCTTATACTACTTAGCGGAAAAC
GAGTCAAAACTATTTGCCTTTATCTATAAATTATTTGGGAGCGTGCCAGGGTGGGA
CAAGAAATTTACGACGGAGCAATTGGAGGCGTTCAATCATCAGTTTGAATCGAGCA
ATTTTACGGCCCGGGAATTGGATTTAGAGTTACAGAAGGATGTGGCACGCGTCATC
TTGACGCAGGTCGGCTCCGGGCCGCAGGAAACGAATGAAAGCTTAATCGACGCCA
AGACGGGCTTGCCGAAGGAATAAGGATCCTGACTCGAGCACCACCACCACCACCA
CTGAGATCGTCGAC

**335_pJL1-(CAT7aa)-PMD_Sau**

TCTAGAAATAATTTTGTTTAACTTTAAGAAGGAGATATACATATGCATATGGAGAAAA
AAATCATCAAAAGCGGTAAAGCACGTGCCCATACCAATATTGCACTGATCAAATATT
GGGGCAAAAAAGATGAAGCCCTGATTATTCCGATGAACAATAGCATTAGCGTTACC

CTGGAAAAGTTCTACACCGAAACCAAAGTGACCTTTAATGATCAGCTGACCCAGGA
TCAGTTTTGGCTGAATGGTGAAAAAGTTAGCGGCAAAGAACTGGAAAAAATCAGCA
AATATATGGATATCGTGCGTAATCGTGCAGGCATTGATTGGTATGCAGAAATTGAA
AGCGATAACTTTGTTCCGACCGCAGCAGGTCTGGCAAGCAGCGCAAGCGCCTATG
CAGCACTGGCAGCAGCATGTAATCAGGCACTGGATATGCAGCTGAGCGATAAAGA
CCTGAGCCGTCTGGCACGTATTGGTAGCGGTAGCGCAAGCCGTAGCATTTATGGT
GGTTTTGCAGAATGGGAGAAAGGTTATAGTGATGAAACCAGCTATGCAGTTCCGCT
GGAAAGCAATCATTTTGAAGATGATCTGGCCATGATCTTCGTTGTGATTAACCAGC
ATAGCAAAAAAGTTCCGAGCCGTTATGGTATGAGTCTGACCCGTAATACCAGCCGT
TTTTATCAGTACTGGCTGGATCATATTGATGAGGACCTGGCAGAAGCAAAAGCAGC
AATTCAGGATAAAGATTTCAAACGTCTGGGCGAAGTGATTGAAGAAAATGGTCTGC
GTATGCATGCAACCAATCTGGGTAGCACCCCTCCGTTTACCTATCTGGTTCAAGAA
AGCTATGACGTTATGGCACTGGTTCATGAATGTCGTGAAGCAGGTTATCCGTGTTA
TTTTACCATGGATGCAGGTCCGAATGTGAAAATTCTGGTGGAAAAAAAGAACAAAC
AGCAGATCATCGATAAACTGCTGACGCAGTTTGATAACAACCAGATTATTGACAGC
GATATCATTGCCACCGGCATTGAAATTATCGAATTAAGTCGACGTCGAC

**336_pJL1-(CAT7aa)-PMD_Spn**

TCTAGAAATAATTTTGTTTAACTTTAAGAAGGAGATATACATATGCATATGGAGAAAA
AAATCGATCGTGAACCGGTTACCGTTCGTAGCTATGCAAATATTGCCATCATCAAAT
ACTGGGGCAAAAAGAAAGAAAAAGAGATGGTTCCGGCAACCAGCAGCATTAGTCT
GACCCTGGAAAATATGTATACCGAAACCACACTGAGTCCGCTGCCTGCAAATGTTA
CCGCAGATGAATTTTACATTAATGGCCAGCTGCAGAACGAAGTTGAACATGCAAAA
ATGAGCAAAATCATCGATCGTTATCGTCCGGCAGGCGAAGGTTTTGTTCGTATTGA
TACCCAGAATAATATGCCGACCGCAGCAGGTCTGAGCAGCAGCAGTAGCGGTCTG
AGCGCACTGATTAAAGCATGTAATGCCTATTTCAAACTGGGCTTAGATCGTAGTCA
GCTGGCACAAGAAGCAAAATTTGCAAGCGGTAGCAGCAGCCGTTCATTTTATGGTC
CGTTAGGTGCATGGGATAAAGATAGCGGTGAAATTTATCCGGTTGAAACCGATCTG
AAACTGGCAATGATTATGCTGGTTCTGGAAGATAAGAAAAAACCGATTAGCAGCCG
TGATGGTATGAAACTGTGTGTTGAAACCAGCACCACCTTTGATGATTGGGTTCGTC
AGAGCGAAAAAGATTATCAGGATATGCTGATCTATCTGAAAGAGAACGATTTCGCC
AAAATTGGTGAACTGACCGAAAAAAATGCCCTGGCAATGCATGCAACCACCAAAAC
CGCAAGTCCGGCATTTAGCTATCTGACCGATGCAAGCTATGAAGCAATGGCATTTG
TTCGTCAGCTGCGTGAAAAAGGTGAAGCATGTTATTTTACCATGGATGCAGGTCCG
AATGTGAAAGTTTTTTGCCAAGAAAAGGATCTGGAACACCTGAGCGAAATTTTTGG
TCAGCGTTATCGCCTGATTGTTAGCAAAACCAAAGACCTGAGCCAGGATGATTGTT
GTTAAGTCGACGTCGAC

**337_pJL1-(CAT7aa)-PMD_Pku**

TCTAGAAATAATTTTGTTTAACTTTAAGAAGGAGATATACATATGCATATGGAGAAAA
AAATCGCCAATGCAGAAAAATGGGTTCTGACCGTTACCGCACAGACCCCGACCAA
TATTGCAGTTATCAAATATTGGGGTAAAGGCGACGAGGATCTGATTCTGCCGATTA
ATGATAGCATTAGCGTTACCTTAGATCCGGACCATCTGTGTACCACCACCAGTGTT
GCAGTTAGTCCGGCATTTACCCATGATCGTATGTGGCTGAATGGTAAAGAAGTTAG
CCTGTCAGGCGGTCGTTTTCAGAATTGTCTGCGTGAAGTTCGTAGCTGTGCAAATG
ATGTTGAAGATGAAAAAGAAGGCGTTCTGAAAAGCCTGAAAGGTCTGGGTGATCTG
CATGTTCACATGTGTCCGACCATTGATTTTCCGACCGCAGCAGGTCTGGCAAGCAG
CGCAGCTGGCCTGGCATGTCTGGTTTTTAGCCTGGCAAAACTGATGAACGTGAAA
GAAGATCATAGCCGTCTGAGCGCAATTGCACGTCAAGGTAGCGGTAGCGCATGTC
GTAGCCTGTATGGTGGTTTTGTTAATGGTAGCTTTCTGAAAGAGGAAAACGGTAGC
GATAGCATTGCAGTTCAGCTGGCCGATGAAAAACATTGGGATGATCTGGTTATTAT
CATTGCCGTTGTTAGCAGCCGTCAGAAAGAAACCAGCAGCACCAGCGGTATGCGT
GAAACCGTTGAAACCAGTATGCTGCTGCAACATCGTGCAAAAGAAGTTGTGCCGG
AACGTATTATTCAGATGGAAGAGGCCATTAAAAACCGCGATTTTCCAGCATTTGCA
CGTCTGACCTGTGCAGATAGCAATCAGTTTCATGCAGTTTGTCTGGATACCCTGCC
TCCGATCTTTTATATGAATGATACCAGCCATCGTATCATCAGCTGTGTGGAAAAATG
GAATTGTAGCGAAGGTACACCGCAGGTTGCATATACCTTTGATGCAGGTCCGAAAT
GCCGTATTAACTTTACCCAGACCGAAAAACTGCTGCCGAATTTTTTCAAAGGTTGC
AGCTTTCATTTTCCGCCTAATAGCGATACCGATCTGAATAGCTATGTTATTGGTGAT
CAGACCATTCTGCAGGATGCAGGTATTAAAGACCTGAAAGATATTGAAGCACTGAG
CACCCCTCCGGAAACCAAAGAAAATCTGAGTGCACAGAAATATCGTGGTGATGTGA
GCTATTTTATCTGCACCAAACCTGGTCGTGGTCCGGCAGTTGTTAATGATGAAAGC
CGTAGCCTGATTAATCCGGAATTTGGTCTGCCGAAATAAGTCGACGTCGAC

**338_pJL1-(CAT7aa)-PMD_Pze**

TCTAGAAATAATTTTGTTTAACTTTAAGAAGGAGATATACATATGCATATGGAGAAAA
AAATCACCGATGCAGTTCGTGACATGATTGCCCGTGCAATGGCAGGCGCAACCGA
TATTCGTGCAGCCGAAGCCTATGCACCGAGTAATATTGCACTGAGCAAATATTGGG
GTAAACGTGATGCAGCACGTAATCTGCCGCTGAATAGCAGCGTTAGCATTAGCCT
GGCAAATTGGGGTAGTCATACCCGTGTTGAAGGTAGCGGCACCGGTCATGATGAA
GTGCATCATAATGGCACCCTGTTAGATCCGGGTGATGCATTTGCACGTCGTGCACT
GGCATTTGCAGACCTGTTTCGTGGTGGTCGTCATCTGCCTCTGCGTATTACCACAC
AGAATAGCATTCCGACCGCAGCAGGTCTGGCAAGCAGCGCAAGCGGTTTTGCAGC
ACTGACCCGTGCATTAGCCGGTGCATTTGGTCTGGATCTGGATGATACCGATCTGA
GCCGTATTGCACGTATTGGTAGCGGTAGCGCAGCCCGTAGCATTTGGCATGGTTT
TGTTCGTTGGAATCGTGGTGAAGCCGAAGATGGTCATGATAGCCATGGTGTTCCG
CTGGATCTGCGTTGGCCTGGTTTTCGTATTGCAATTGTTGCAGTTGATAAAGGTCC
GAAACCGTTTAGCAGCCGTGATGGCATGAATCATACCGTTGAAACCAGTCCGCTGT
TTCCGCCTTGGCCTGCACAGGCCGAAGCAGATTGTCGTGTTATTGAAGATGCAATT
GCCGCACGTGATATGGCAGCACTGGGTCCGCGTGTGGAAGCAAATGCCCTGGCA
ATGCATGCAACCATGATGGCAGCCCGTCCGCCTCTGTGTTATCTGACCGGTGGTA

GCTGGCAGGTTCTGGAACGTCTGTGGCAGGCACGTGCAGATGGTCTGGCAGCATT
TGCCACCATGGATGCAGGTCCGAATGTTAAACTGATTTTTGAAGAAAGCAGTGCCG
CAGATGTTCTGTACCTGTTTCCGGATGCAAGCCTGATTGCACCGTTTGAAGGTCGT
TAAGTCGACGTCGAC

**339_pJL1-(CAT7aa)-PMD_Hme**

TCTAGAAATAATTTTGTTTAACTTTAAGAAGGAGATATACATATGCATATGGAGAAAA
AAATCAAAGCAACCGCAAAAGCACATCCGATTCAAGGTCTGGTTAAATATCATGGT
ATGCGCGATCCGGAAATTCGTCTGCCGTATCATGATTCAATTAGCGTTTGTACCGC
ACCGAGCCATACCAAAACCACCGTTGAATTTCTGCCGGATGCAGATGAAGATGTTT
ATGTTATTGGTGGTGAAGAGGTTGAAGGTCGTGGTGCAGAACGTATTCGTGATGTT
GTTGAACATGTGCGTGATCTGGCAGATTTTGATCATCGTGTTCGTCTGGAAAGCGA
AAATAGCTTTCCGAGCAATATTGGTTTTGGCAGCAGCAGCTCAGGTTTTGCCGCAG
CCGCAATGGCACTGGCAGAAGCAGCCGATCTGGATCTGACCCGTCCGGAAATCAG
CACCATTGCACGTCGTGGTAGCAGCAGCGCAGCACGTGCAGTTACCGGTGCATTT
AGCCATCTGTATAGCGGTATGAATGATACCGATTGTCGTAGCGAACGTATTGAAAC
GGATCTGGAAGATGATCTGCGTATTGTTGCAGCCCATGTTCCGGCATATAAAGAAA
CCGAACAGGCACATGCAGAAGCCGCAGATAGCCACATGTTTCAGGCACGTATGGC
ACACATGCATAAGCAGATTGATGATATGCGTGATGCACTGTATGAAGCCGATTTTG
ATGCAGCATTTGAACTGGCCGAACATGATAGCCTGAGCCTGGCAGCAACCACCAT
GACCGGTCCGGCAGGTTGGGTTTATTGGCAGCCTCGTACCATTGCAGTTTTTAATG
CAATTCGTGAACTGCGTGCCGAAGAAGATATTCCTGCATATTTTAGCACCGATACC
GGTGCCAGCGTTTATATCAATACCACCACCGAATATGTTGACCGTGTTGAAAAAGT
TGTTGCCGATTGTAATGTTGAAACCGATGTTTGGGAAGTTGGTGGTCCTGCCGAAA
TTCTGGATGAAAGTGATGCCCTGTTTTAAGTCGACGTCGAC

**340_pJL1-(CAT7aa)-PMD_Zga**

TCTAGAAATAATTTTGTTTAACTTTAAGAAGGAGATATACATATGCATATGGAGAAAA
AAATCACCGTGAAAGAATTTATCCCGAGTCCGTATACCAAACCGGTTGCAAGCGGT
AATACCCGTTATAAAAGCCCGAGTAATATTGCCCTGGTGAAATATTGGGGCAAAAA
AGAAAATCAGATTCCGGCAAATCCGAGCATTAGCTTTACCCTGAATGAATGTGCAA
CCGTTACCACACTGAGCTATCGTAAAGCAGATCGTCCGAATGATGCATTTAGCTTT
GAAATTAGCCTGGACGGCAAGAAAGAAGAAGGTTTTAAACCGAAAATCAAAACCTT
TTTCGAACGCGTGTATCCGTATCTGCCGTTTCTGAAAGAATATCACTTTGAGATTGA
AACCAGCAACAGCTTTCCGCATAGCAGCGGTATTGCAAGCAGCGCCAGCGGTATG
AGCGCACTGGCACTGTGTCTGATGGAAATTGAACGTAATTTAGATCCGGGTATGAG
TGCCGATTTTTTCAACCGTAAAGCAAGCTTTCTGGCACGTTTAGGTAGCGGTAGCG
CAGCACGTAGCATTAAAGGTAGCCTGGTTCAGTGGGGTGAACATGCAGGCACCGA
AGGTAGCAGCGATCTGTATGGTATTGAATATCCGTATAAAGTGCACAGCGTGTTCA
ACGATTATTGCGATACCATTCTGCTGGTTGATAAAGGTCAGAAACAGGTTAGCAGC

ACCGTTGGTCATGATCTGATGCATAATCATCCGTTTAGCAAACAGCGTTTTGATCAG
GCACATGAAAATCTGAGCAAACTGCGTAGCATTTTTGAAAGCGGCAATCTGGATGA
ATTTATTGGTCTGGTTGAAAGCGAAGCACTGACCCTGCATGCAATGATGATGACCA
GCCGTCCGTATTTTATCCTGATGAAACCGAATACGCTGGAAATTATCAATCGCATTT
GGGCATATCGTGAAGCCACCAAAACACATGTTTGTTTTACCCTGGATGCCGGTGCA
AATGTTCATGTTCTGTATCCGAAAAATGAAAAGGCACTGGTGGAACGTTTTATTGCA
GATGAACTGGCAGGTTATTGTCAGAATGGTCAGTTTATTCATGATCATGTTGGTCGT
GGTGCCCGTAAAATCAATTAAGTCGACGTCGAC

**284_pJL1-(CAT5aa)-IDI_Eco**

TCTAGAAATAATTTTGTTTAACTTTAAGAAGGAGATATACATATGGAGAAAAAAATC
CAAACGGAACACGTCATTTTATTGAATGCACAGGGAGTTCCCACGGGTACGCTGG
AAAAGTATGCCGCACACACGGCAGACACCCGCTTACATCTCGCGTTCTCCAGTTG
GCTGTTTAATGCCAAAGGACAATTATTAGTTACCCGCCGCGCACTGAGCAAAAAAG
CATGGCCTGGCGTGTGGACTAACTCGGTTTGTGGGCACCCACAACTGGGAGAAAG
CAACGAAGACGCAGTGATCCGCCGTTGCCGTTATGAGCTTGGCGTGGAAATTACG
CCTCCTGAATCTATCTATCCTGACTTTCGCTACCGCGCCACCGATCCGAGTGGCAT
TGTGGAAAATGAAGTGTGTCCGGTATTTGCCGCACGCACCACTAGTGCGTTACAG
ATCAATGATGATGAAGTGATGGATTATCAATGGTGTGATTTAGCAGATGTATTACAC
GGTATTGATGCCACGCCGTGGGCGTTCAGTCCGTGGATGGTGATGCAGGCGACA
AATCGCGAAGCCAGAAAACGATTATCTGCATTTACCCAGCTTAAATAAGGATCCTA
ACTCGAGCACCACCACCACCACCACTGAGATCGTCGAC

**285_pJL1-(CAT5aa)-IDI_Bsu**

TCTAGAAATAATTTTGTTTAACTTTAAGAAGGAGATATACATATGGAGAAAAAAATCA
CCCGTGCAGAACGTAAACGTCAGCATATCAATCATGCACTGAGCATTGGTCAGAAA
CGTGAAACCGGTCTGGATGATATTACCTTTGTTCATGTTAGCCTGCCGGATCTGGC
ACTGGAACAGGTTGATATTAGCACCAAAATTGGTGAACTGAGCAGCAGCAGCCCG
ATTTTTATCAATGCAATGACCGGTGGTGGTGGTAAACTGACCTATGAAATTAACAAA
AGCCTGGCACGTGCAGCAAGCCAGGCAGGTATTCCGCTGGCAGTTGGTAGCCAG
ATGAGCGCACTGAAAGATCCGTCAGAGCGTCTGTCTTATGAGATCGTGCGGAAAG
AAAATCCGAATGGACTCATTTTTGCGAATTTAGGTAGCGAGGCTACCGCCGCGCA
GGCTAAAGAGGCGGTCGAAATGATTGGCGCGAACGCGCTGCAGATCCATCTGAAC
GTAATTCAGGAGATCGTGATGCCGGAAGGCGATCGTTCGTTTAGTGGTGCCCTCA
AGCGTATTGAGCAGATTTGTTCTCGCGTCTCAGTGCCGGTTATCGTGAAAGAGGTA
GGATTTGGAATGTCGAAAGCTTCCGCGGGTAAGCTGTACGAGGCAGGGGCCGCT
GCAGTTGATATTGGCGGCTATGGGGGTACGAACTTCAGCAAAATTGAAAACCTCC
GTCGCCAGCGCCAAATCTCCTTCTTCAACAGCTGGGGCATCTCTACGGCTGCATC
ATTAGCAGAGATCCGTTCTGAATTTCCGGCAAGCACCATGATTGCAAGCGGTGGTC
TGCAGGATGCACTGGATGTTGCAAAAGCAATTGCACTGGGTGCAAGCTGTACCGG

TATGGCAGGTCATTTTCTGAAAGCACTGACCGATAGCGGTGAAGAAGGTCTGCTG
GAAGAAATTCAGCTGATTCTGGAAGAACTGAAACTGATTATGACCGTTCTGGGTGC
ACGTACCATTGCCGATCTGCAGAAAGCACCGCTGGTTATTAAAGGTGAAACCCATC
ATTGGCTGACAGAACGTGGTGTTAATACCAGCAGCTATAGCGTTCGTTAAGGATCC
TGACTCGAGCACCACCACCACCACCACTGAGATCGTCGAC

**341_pJL1-(CAT7aa)-IDI_Sce**

TCTAGAAATAATTTTGTTTAACTTTAAGAAGGAGATATACATATGCATATGGAGAAAA
AAATCACCGCAGATAATAACAGCATGCCGCATGGTGCAGTTTCAAGCTATGCAAAA
CTGGTTCAGAATCAGACACCGGAAGATATCCTGGAAGAATTTCCTGAAATTATTCC
GCTGCAGCAGCGTCCGAATACACGTAGCAGCGAAACCAGCAATGATGAAAGCGGT
GAAACCTGTTTTAGCGGTCATGATGAAGAACAAATCAAGCTGATGAACGAAAACTG
CATTGTTCTGGATTGGGATGATAATGCAATTGGTGCAGGCACCAAAAAAGTTTGTC
ATCTGATGGAAAACATCGAGAAAGGTCTGCTGCATCGTGCATTTAGCGTGTTTATC
TTTAATGAACAGGGTGAACTGCTGCTGCAACAGCGTGCAACCGAAAAAATCACCTT
TCCGGATCTGTGGACCAATACCTGTTGTAGCCATCCGCTGTGTATTGATGATGAAC
TGGGTCTGAAAGGTAAACTGGACGATAAAATCAAAGGTGCAATTACCGCAGCCGTT
CGCAAACTGGATCACGAACTGGGTATTCCGGAAGATGAAACCAAAACACGTGGCA
AATTTCATTTTCTGAACCGCATCCATTATATGGCACCGAGCAATGAACCGTGGGGT
GAACATGAAATTGATTACATCCTGTTCTACAAAATCAACGCCAAAGAAAACCTGACC
GTTAATCCGAATGTTAATGAAGTGCGTGATTTCAAATGGGTGAGCCCGAATGATCT
GAAAACCATGTTTGCCGATCCGAGCTATAAATTCACCCCGTGGTTTAAAATCATCT
GCGAAAACTACCTGTTTAACTGGTGGGAACAGCTGGATGATCTGAGCGAAGTTGA
AAATGATCGTCAGATTCATCGTATGCTGTAAGTCGACGTCGAC

**342_pJL1-(CAT7aa)-IDI_Sly**

TCTAGAAATAATTTTGTTTAACTTTAAGAAGGAGATATACATATGCATATGGAGAAAA
AAATCGTTGATGTTATTGCCGATGCAAATATGGATGCAGTTCAGCGTCGTCTGATG
TTTGATGATGAATGTATTCTGGTGGACGTGAACGATAAAGTTGTTGGTCATGAGAG
CAAATATAACTGCCACCTGATGGAAAAAATCGAAAGCGAAAATCTGCTGCATCGTG
CCTTTAGCGTTTTTCTGTTTAACAGCAAATATGAGCTGCTGCTGCAACAGCGTAGC
GCAACCAAAGTTACCTTTCCGCTGGTTTGGACCAATACCTGTTGTAGCCATCCGCT
GTATCGTGAAAGCGAACTGATTGAAGAAAATGCACTGGGTGTTCGTAATGCAGCAC
AGCGTAAACTGCTGGATGAACTGGGTATTCCGGCAGAAGATGTTCCGGTTGATCA
GTTTACACCGCTGGGTCGTATGCTGTATAAAGCACCGAGTGATGGTAAATGGGGT
GAACATGAACTGGATTATCTGCTGTTTATTGTGCGTGATGTTAACGTTCATCCGAAT
CCTGATGAAGTTGCCGATATCAAATACGTGAATCAAGAACAGCTGAAAGAACTGCT
GCGTAAAGCAGATGCCGGTGAAGAAGGTCTGAAACTGAGCCCGTGGTTTCGTCTG
GTTGTTGATAATTTTCTGTTCAAATGGTGGGACCATGTTGAAAAAGGCACCATTCAA
GAGGCAGCAGATATGAAAACCATTCACAAACTGACCTAAGTCGACGTCGAC

**343_pJL1-(CAT7aa)-IDI_Str**

TCTAGAAATAATTTTGTTTAACTTTAAGAAGGAGATATACATATGCATATGGAGAAAA
AAATCACCAGCGCACAGCGTAAAGATGATCATGTTCGTCTGGCAATTGAACAGCAT
AATGCACATAGCGGTCGTAACCAGTTTGATGATGTTAGCTTTGTTCATCATGCACTG
GCAGGTATTGATCGTCCGGATGTTAGCCTGGCAACCAGCTTTGCAGGTATTAGCTG
GCAGGTTCCGATTTATATCAATGCAATGACCGGTGGTAGCGAAAAAACCGGTCTGA
TTAATCGTGATCTGGCAACCGCAGCACGTGAAACCGGTGTTCCGATTGCAAGCGG
TAGCATGAATGCATATATCAAAGATCCGAGCTGTGCAGATACCTTTCGTGTTCTGC
GTGATGAAAATCCGAATGGTTTTGTGATTGCCAATATTAACGCAACCACCACCGTT
GATAATGCCCAGCGTGCAATTGATCTGATTGAAGCAAATGCACTGCAGATCCATAT
TAACACCGCACAAGAAACCCCGATGCCGGAAGGTGATCGTAGCTTTGCAAGCTGG
GTTCCGCAGATTGAAAAAATTGCAGCAGCAGTTGATATTCCGGTGATCGTTAAAGA
AGTTGGTAACGGTCTGAGCCGTCAGACCATTCTGCTGCTGGCCGATCTGGGTGTT
CAGGCAGCAGATGTGAGCGGTCGTGGTGGCACCGATTTTGCACGTATTGAAAATG
GTCGTCGTGAACTGGGTGATTATGCATTTCTGCATGGTTGGGGTCAGAGCACCGC
AGCATGTCTGCTGGATGCACAGGATATTAGCCTGCCGGTTCTGGCAAGCGGTGGT
GTTCGTCATCCGCTGGATGTTGTTCGTGCACTGGCCCTGGGTGCACGTGCCGTTG
GTAGCAGCGCAGGTTTTCTGCGTACCCTGATGGATGATGGTGTTGATGCACTGATT
ACCAAACTGACCACCTGGCTGGATCAGCTGGCAGCCCTGCAGACCATGCTGGGA
GCACGTACCCCTGCCGATCTGACCCGTTGTGATGTTCTGCTGCATGGTGAACTGC
GCGATTTTTGTGCAGATCGTGGTATTGATACCCGTCGTCTGGCCCAGCGTAGCAG
CAGTATTGAAGCGCTGCAGACAACCGGTAGCACCCGTTAAGTCGACGTCGAC

**344_pJL1-(CAT7aa)-IDI_Pze**

TCTAGAAATAATTTTGTTTAACTTTAAGAAGGAGATATACATATGCATATGGAGAAAA
AAATCACCGATAGCAAAGATCATCATGTTGCAGGTCGTAAACTGGATCATCTGCGT
GCACTGGATGATGATGCAGATATTGATCGTGGTGATAGCGGTTTTGATCGTATTGC
ACTGACCCATCGTGCACTGCCGGAAGTTGATTTTGATGCAATTGATACCGCAACCA
GCTTTCTGGGTCGTGAACTGAGCTTTCCGCTGCTGATTAGCAGCATGACCGGTGG
TACAGGTGAAGAAATTGAACGTATTAATCGTAATCTGGCAGCCGGTGCCGAAGAG
GCACGTGTTGCAATGGCAGTTGGTAGCCAGCGTGTTATGTTTACCGATCCGAGCG
CACGTGCATCATTTGACCTGCGTGCCCATGCACCGACCGTGCCGCTGCTGGCAAA
TATTGGTGCAGTGCAGCTGAATATGGGTCTGGGTCTGAAAGAATGTCTGGCAGCA
ATTGAAGTTCTGCAGGCAGATGGTCTGTATCTGCACCTGAATCCGCTGCAAGAAGC
AGTTCAGCCGGAAGGTGATCGTGATTTTGCCGATCTGGGTAGCAAAATTGCAGCC
ATTGCACGTGATGTTCCGGTTCCGGTGCTGCTGAAAGAAGTTGGCTGCGGTCTGA
GCGCAGCCGATATTGCAATTGGTCTGCGTGCCGGTATTCGTCATTTTGATGTTGCC
GGTCGCGGTGGCACCAGTTGGAGCCGTATTGAATATCGTCGTCGTCAGCGTGCAG
ATGATGATTTAGGTCTGGTTTTTCAGGATTGGGGACTGCAGACCGTTGATGCACTG

CGTGAAGCACGTCCGGCACTGGCAGCACATGATGGCACCAGCGTTCTGATTGCAA
GCGGTGGTATTCGTAATGGTGTTGATATGGCCAAATGTGTTATTCTGGGTGCCGAT
ATGTGTGGTGTTGCAGCACCGCTGTTAAAAGCAGCACAGAATAGCCGTGAAGCAG
TTGTTAGCGCAATTCGCAAACTGCATCTGGAATTTCGTACCGCAATGTTTCTGTTAG
GTTGTGGCACCCTGGCCGATCTGAAAGATAATAGCAGCCTGATTCGTCAGTAAGTC
GACGTCGAC

**345_pJL1-(CAT7aa)-IDI_Sau**

TCTAGAAATAATTTTGTTTAACTTTAAGAAGGAGATATACATATGCATATGGAGAAAA
AAATCAGCGATTTTCAGCGTGAACAGCGCAAAAATGAACATGTTGAAATTGCAATG
GCACAGAGTGATGCAATGCATAGCGATTTTGATAAAATGCGCTTTGTGCATCATTC
CATTCCGAGCATTAATGTGAACGATATTGATCTGACCAGCCAGACACCGGATCTGA
CAATGACCTATCCGGTTTATATCAATGCAATGACCGGTGGTAGCGAATGGACCAAA
AACATTAATGAAAAACTGGCAGTTGTTGCCCGTGAAACCGGTCTGGCAATGGCAGT
TGGTAGCACCCATGCAGCACTGCGTAATCCGCGTATGGCAGAAACCTTTACCATTG
CACGTAAAATGAATCCGGAAGGCATGATTTTTAGCAATGTTGGTGCAGATGTTCCG
GTTGAAAAAGCACTGGAAGCCGTTGAACTGCTGGAAGCACAGGCACTGCAGATTC
ATGTTAATAGTCCGCAAGAACTGGTTATGCCGGAAGGTAATCGTGAATTTGTTACC
TGGCTGGATAATATTGCAAGCATTGTTAGCCGTGTTTCAGTTCCGGTTATCATTAAA
GAAGTTGGTTTCGGCATGAGCAAAGAACTGATGCATGATCTGCAGCAGATTGGTGT
TAAATATGTTGATGTTAGCGGTAAAGGTGGCACCAACTTTGTGGATATTGAAAATGA
ACGTCGTGCCAACAAAGATATGGATTATCTGAGCAGCTGGGGTCAGAGCACCGTT
GAAAGCCTGCTGGAAACCACCGCATATCAGAGCGAAATTAGCGTTTTTGCAAGCG
GTGGTCTGCGTACACCGCTGGATGCAATTAAAGCCTGGCACTGGGTGCAAAAGC
AACCGGTATGAGCCGTCCGTTTCTGAATCAGGTTGAAAATAATGGTATTGCCCATA
CCGTTGCCTATGTGGAAAGCTTTATTGAACACATGAAAAGCATCATGACCATGCTG
GATGCGAAAAATATCGATGATCTGACACAGAAGCAGATCGTTTTTAGTCCGGAAAT
TCTGAGCTGGATTGAACAGCGTAATCTGAACATTCATCGTGGCTAAGTCGACGTCG
AC

346_pJL1-(CAT7aa)-IDI_Scl

TCTAGAAATAATTTTGTTTAACTTTAAGAAGGAGATATACATATGCATATGGAGAAAA
AAATCGTTATGCCGACCAGTCCGACCTCACCGACCGCAGCAAATAGCGTTAGCAA
TGGCACCAGCAATGATGTTCCGGATGGTGCAGCACGTGAAATTCTGCTGGAACTG
GTTGATGAACATGGCACCACCATTGGCACCGCAGAAAAACTGGCAGCACATCAGC
CTCCGGGTCTGCTGCATCGTGCATTTAGCGTTTTTCTGTTTGATGATCGTGGTCGT
CTGCTGCTGCAACAGCGTGCACTGGGTAAATATCATAGCCCTGGTGTTTGGAGCA
ATACCTGTTGTGGTCATCCGTATCCGGGTGAAGCACCGTTTGCAGCCGCAGCACG
TCGTACCCATGAAGAACTGGGTATTAGTCCGGCACTGCTGGCAGAAGCAGGCACC
GTTCGTTATAATCATCCTGATCCGGATAGCGGTCTGGTTGAACAAGAATATAATCA

CCTGTTTGTTGGTCTGGTTCAGGCAAGTCCGGAACCAGATCCGGAAGAAGTTGGC
GGTACAGTTTTTGTTACACCGGGTGAACTGGCAGAACGTCATGCAGCGGCACCGT
TTAGCTCATGGTTTATGACCGTTCTGGATGCAGCCCGTCCGGCAATTCGTGAACTG
ACCGGTCCGAGCGGTGGTTGGTAAGTCGACGTCGAC

**286_pJL1-(CAT5aa)-GPPS_Agr_F3F**

TCTAGAAATAATTTTGTTTAACTTTAAGAAGGAGATATACATATGGAGAAAAAAATC
GAATTCGACTTCAACAAATACATGGATAGCAAAGCCATGACCGTTAATGAAGCACT
GAATAAAGCAATTCCGCTGCGTTATCCGCAGAAAATCTATGAAAGCATGCGTTATA
GCCTGCTGGCAGGCGGTAAACGTGTTCGTCCGGTTCTGTGTATTGCAGCATGTGA
ACTGGTTGGTGGCACCGAAGAACTGGCAATTCCGACCGCATGTGCAATTGAAATG
ATTCATACCATGAGCCTGATGCATGATGATCTGCCGTGTATTGATAATGATGACCT
GCGTCGTGGTAAACCGACCAATCATAAAATCTTTGGTGAAGATACCGCAGTGACCG
CAGGTAATGCACTGCATAGTTATGCATTTGAACATATTGCAGTGAGCACCAGCAAA
ACCGTTGGTGCAGATCGTATTCTGCGTATGGTTAGCGAACTGGGTCGTGCAACCG
GTAGCGAAGGTGTTATGGGTGGTCAGATGGTTGATATTGCAAGTGAAGGTGATCC
GAGCATTGATCTGCAGACCCTGGAATGGATTCATATTCATAAAACCGCAATGCTGC
TGGAATGTAGCGTTGTTTGTGGTGCAATTATTGGTGGTGCAAGCGAAATTGTTATT
GAACGTGCCCGTCGTTATGCACGTTGTGTTGGTCTGCTGTTTCAGGTTGTTGATGA
TATTCTGGATGTGACCAAAAGCAGTGATGAACTGGGCAAAACCGCAGGCAAAGAC
CTGATTAGCGATAAAGCAACCTATCCGAAACTGATGGGTCTGGAAAAAGCCAAAGA
ATTTTCAGATGAACTGCTGAATCGTGCCAAAGGTGAACTGAGCTGTTTTGATCCGG
TTAAAGCAGCACCGCTGCTGGGTCTGGCAGATTATGTTGCATTTCGTCAGAACTAA
GGATCCTGACTCGAGCACCACCACCACCACCACTGAGATCGTCGAC

**287_pJL1-(CAT5aa)-GPPS2_Pab**

TCTAGAAATAATTTTGTTTAACTTTAAGAAGGAGATATACATATGGAGAAAAAAATC
GAGTTCGACTTTGACAAGTATATGCACAGTAAAGCAATCGCCGTAAACGAGGCTTT
GGATAAAGTGATCCCGCCCCGCTACCCCCAAAAGATTTATGAATCAATGCGCTATA
GTTTACTGGCAGGTGGAAAACGTGTGCGCCCGATCCTGTGTATTGCGGCTTGCGA
GCTGATGGGAGGCACTGAGGAGTTAGCTATGCCGACCGCATGCGCAATTGAGATG
ATTCATACCATGTCTCTTATCCACGACGACTTGCCGTATATCGATAATGATGACTTA
CGTCGGGGAAAACCGACCAACCATAAAGTTTTCGGCGAAGACACCGCTATTATCG
CAGGAGATGCTCTGCTGAGCTTAGCATTCGAGCATGTTGCAGTGAGTACTAGCCG
GACTCTGGGTACTGATATCATCCTGCGCCTGCTGAGCGAAATCGGTCGTGCAACG
GGCTCGGAGGGCGTCATGGGCGGCAAGTAGTTGATATTGAATCGGAGGGAGAT
CCCTCTATCGACCTGGAAACCCTGGAGTGGGTACATATCCATAAAACGGCAGTTTT
GCTCGAATGCTCGGTCGTTTGCGGTGCAATTATGGGGGGTGCCTCAGAGGATGAT
ATTGAACGTGCCCGTCGTTACGCGCGCTGCGTCGGCTTGCTGTTTCAGGTCGTCG
ATGATATCCTTGATGTCAGCCAGTCTAGTGAAGAACTGGGAAAAACCGCTGGCAAG

GATCTGATTAGCGACAAAGCCACTTACCCAAAATTAATGGGTCTGGAAAAAGCTAA
AGAATTTGCCGACGAATTATTGAACCGTGGTAAGCAGGAACTGTCATGTTTTGACC
CCACGAAAGCAGCACCTCTGTTTGCGCTTGCAGACTATATTGCCAGCCGGCAGAA
CTAAGGATCCTGACTCGAGGTCGAC

**288_pJL1-(CAT5aa)-GPPS_Str**

TCTAGAAATAATTTTGTTTAACTTTAAGAAGGAGATATACATATGGAGAAAAAAATCA
CCACCGATACCGGTCAGGATGCAGTTGGTCTGGCACGTCGTACCGGTGCAGACCT
GCTGCATCGTGTTGAAGATCGTCTGCGTAGCCTGCTGGCAGTCGAACGTGATGCA
TGGGCGGCCGTGCACGAGCAGGCAGTTGTGCCGGTTGACGCCTTATCAGAGCTG
ATCGCAAGTGGCGGCAAACGCATCCGCCCGGCATTCTGTATCACGGGTTATTTAG
CTGCCGGTGGCGATCCAGCCGAACCAGGTATTGTGGCTGCGGGCGCGGCTCTGG
AAATGTTGCATCTTTCAGCCCTGGTGCACGACGACGTTTTAGATGACTCCAGCTCA
CGCCGTGGGGTACCGACCGTACATACCCAAGCTATGGCCCTGCATGAATCTAGCG
GTTGGCAGGGCGAACCGCGCCGCTACGGGGAAGGCGTGGCTATCCTGGTAGGC
GACCTGGCCTTAGTATACTCAGAAGAGTTAATGGCGGAAGCGCCTCGCCGCGTCC
TGCCAGAATGGAATAAACTGCGTTCGGAAGTTATGATCGGTCAATACATGGACGTA
CATGCAGCCGCTGAATTTAGCGTGGATCCGCGTAGCTCCCGCCTTATTGCGCGCA
TTAAATCTGGGCGTTATACTATCCATCGTCCATTAGTAGTAGGCGCCAAACGGGGC
CGCGGTCGCGGTGATCTGGCGCCAGCATTAGAAGAGTACGGTGAAGCCGTGGGC
GAGGCCTTCCAACTGCGCGATGACTTACTGGATGCGTCTGCAACGCCGGCCGAAA
CCGGGAAGCCAACTGGCCTGGATTTCACACAGCATAAAATGACCTTACTGCTGGG
CTGGGCGATGCAGCGCGATGAGCATATCCATACGCTGGTAACGGAACCAGGGCA
CACGCCCGATGAGGTGCGCCGGCGTCTGTTAGATACGGGCGTTCCGGCCGATGT
AGAACGGCATATCGCGGGCTTAGTGGAACGTGGCTGCAAAGCCATCGCTGATGCA
CCTGTCGATCAGGTTTGGCGGGGCGAGCTGGCAGCCATGGCCGGCCGTGTCGCG
TATCGTACGGCATAAGGATCCTGACTCGAGGTCGAC

**347_pJL1-(CAT7aa)-GPPS_Pgl**

TCTAGAAATAATTTTGTTTAACTTTAAGAAGGAGATATACATATGCATATGGAGAAAA
AAATCGAATTTGACTTCAAAGAATACATGCGCAGCAAAGCCATGAGCGTTAATGAA
GCACTGGATCGTGCAGTTCCGCTGCGTTATCCGGAAAAAATTCATGAAGCAATGC
GTTATAGCCTGCTGGCAGGCGGTAAACGTGTTCGTCCGATTCTGTGTATTGCAGCA
TGTGAACTGGTTGGTGGTAGCGAAGAACTGGCAATGCCGACCGCATGTGCAATGG
AAATTATTCATACCATGAGCCTGATCCATGATGATCTGCCTCCGATGGATAATGATG
ACCTGCGTCGTGGTAAACCGACCAATCATAAAGTTTTTGGTGAAGGCACCGCAGTT
TTAGCCGGTGATGCACTGCTGAGCTTTGCATTTGAACATATTGCAGTTAGCACCAG
CAAAACCGTTGAAAGCGATCGTGTTCTGCGTGTTGTTAGCGAACTGGGTCGTGCA
ATTGGTAGTGAAGGTGTTGCCGGTGGTCAGGTTGCAGATATTACCAGCCAGGGTA
ATCCGAGCGTTGGTCTGGAAACCCTGGAATGGATTCATATTCATAAAACCGCAGTT

CTGCTGGAATGTAGCGTTGCAAGCGGTGCAATTATTGGTGGTGCAAGCGAAGATG
AAATTGAACGTGTGCGTAAATATGCACGTTGTGTTGGTCTGCTGTTTCAGGTTGTT
GATGATATTCTGGATGTTACCAAAAGCAGCGAGGAACTGGGTAAAACAGCAGCAAA
AGACCTGCTGAGCGATAAAGCAACCTATCCGAAACTGATGGGTCTTGAAAAAGCAA
AAGAATTTGCAGATGAACTGCTGGGCAAAGCCAAAGAAGAACTGAGCTTTTTTAAC
CCGACCAAAGCAGCACCGCTGCTGGGTTTAGCAGATTATATTGCACAGCGTCAGA
ACTAAGTCGACGTCGAC

**348_pJL1-(CAT7aa)-GPPS_Pku**

TCTAGAAATAATTTTGTTTAACTTTAAGAAGGAGATATACATATGCATATGGAGAAAA
AAATCAGCCTGGTTAATAGCATTACCTGGTCACAGACCAGCAGCATTCTGAACATT
CAGAGCAACATTAGCAAAAAACTGACCCCGTTTAGCATTCTGCCGCATCCGCTGAC
CAATAATCTGCCGATTAGCCTGTTTCCGAATCCGAAAAGCAATATCAGCAATAGCA
ATACACCGCTGAGCGCAATTCTGACCAAAGATCAGAAACCGCAGAATCCGCCTAC
CACACCGACCTTTGATTTCAAAAGCTATATGCTGCAGAAAGCCGATAGCGTTAATA
AAGCACTGGATGATAGCATTCCGCTGACAGAACCGCTGAAAATTCAAGAAAGCATG
CGTTATAGCCTGCTGGCAGGCGGTAAACGTATTCGTCCGATGCTGTGTATTGCAG
CATGTGAACTGGTTGGTGGTGATGAAAGCACCGCAATGCCTGCAGCCTGTGCAGT
TGAAATGGTTCATACCATGAGCCTGATGCATGATGATCTGCCGTGTATGGATAATG
ATGACCTGCGTCGTGGTAAACCGACCAATCATAAAGTTTTTACCGAAGATGTTGCC
GTGTTAGCCGGTGATGCAATGCTGGCATTTAGCTTTGAACATGTTGCAAGCCTGAC
AAAAGGTGTTTGTAGCGAACGTATTGTGCGCGTTATTTATGAACTGGCAAAATGTG
TTGGTTGCGAAGGTCTGGTTGCAGGTCAGGTTGTTGATATTTGCAGCGAAGGTATG
GATGAAGTTGGTCTGGAACATCTGGAATTTATCCATCTGAATAAAACCGCAGCACT
GCTGGAAGGTAGCGTTGTTCTGGGTGCAATTTTAGGTGGTGGTAGTGATGAAGAA
GTTGAAAAACTGCGTAATTTTGCCCGTTGTATTGGTCTGCTGTTTCAGGTTGTGGAT
GATATTCTGGATGTTACCAAAAGCAGCAAAGAACTGGGTAAAACAGCAGGTAAAGA
CCTGGTTGCCGATAAAACCACCTATCCGAAACTGATTGGTATCGAGAAAAGCAAAG
AATTTGCCGAACGTCTGAATCGTGAAGCAAAAGAACACCTGGCAGGTTTTGATCAG
AATAAAGCAGCACCGCTGATTGCACTGGCCGATTATATTGCATATCGTGACAATTA
AGTCGACGTCGAC

**349_pJL1-(CAT7aa)-GPPS*_Sce**

TCTAGAAATAATTTTGTTTAACTTTAAGAAGGAGATATACATATGCATATGGAGAAAA
AAATCGCCAGCGAAAAAGAAATTCGTCGTGAACGTTTTCTGAACGTGTTTCCGAAA
CTGGTTGAAGAACTGAATGCAAGCCTGCTGGCCTATGGTATGCCGAAAGAAGCAT
GCGATTGGTACGCACATAGCCTGAATTATAACACCCCTGGTGGTAAACTGAATCGT
GGTCTGAGCGTTGTTGATACCTATGCAATTCTGAGCAATAAAACCGTGGAACAGCT
GGGTCAAGAGGAATATGAAAAAGTTGCAATTCTTGGCTGGTGCATTGAACTGCTGC
AGGCATATTTTCTGGTTGCAGATGATATGATGGACAAAAGCATTACCCGTCGTGGT

CAGCCGTGTTGGTATAAAGTTCCGGAAGTTGGTGAAATTGCCATCAATGATGCATT
TATGCTGGAAGCAGCAATCTACAAACTGCTGAAAAGCCATTTTCGCAACGAGAAAT
ATTACATCGATATCACCGAACTGTTTCACGAAGTTACCTTTCAGACCGAACTGGGT
CAGCTGATGGATCTGATTACCGCACCGGAAGATAAAGTTGATCTGAGCAAATTCAG
CCTGAAAAAGCATAGCTTTATCGTGACCTTTGAAACCGCCTATTATAGCTTTTATCT
GCCGGTTGCACTGGCAATGTATGTTGCAGGTATTACCGATGAAAAAGACCTGAAAC
AGGCACGTGATGTTCTGATTCCGCTGGGTGAATATTTTCAGATCCAGGATGATTAT
CTGGATTGCTTTGGTACACCGGAACAAATTGGTAAAATTGGCACCGATATCCAGGA
TAACAAATGTAGCTGGGTTATTAACAAAGCACTGGAACTGGCAAGCGCAGAACAGC
GTAAAACCCTGGATGAAAATTATGGCAAAAAAGATAGCGTTGCCGAGGCCAAATGC
AAGAAAATCTTTAACGACCTGAAAATCGAGCAGCTGTATCACGAATATGAAGAATC
AATTGCCAAGGACCTGAAAGCCAAAATTAGCCAGGTTGATGAAAGCCGTGGTTTTA
AAGCAGATGTGCTGACCGCATTTCTGAACAAAGTGTATAAACGCAGCAAATAAGTC
GACGTCGAC

**318_pJL1-(CAT5aa)-ispA_Ec**

TCTAGAAATAATTTTGTTTAACTTTAAGAAGGAGATATACATATGGAGAAAAAAATC
GACTTTCCGCAGCAACTCGAAGCCTGCGTTAAGCAGGCCAACCAGGCGCTGAGCC
GTTTTATCGCCCCACTGCCCTTTCAGAACACTCCCGTGGTCGAAACCATGCAGTAT
GGCGCATTATTAGGTGGTAAGCGCCTGCGACCTTTCCTGGTTTATGCCACCGGTC
ATATGTTTGGCGTTAGCACAAACACGCTGGACGCACCCGCTGCTGCCGTAGAGTG
TATCCACGCTTACTCATTAATTCATGATGATTTACCGGCGATGGATGATGACGATCT
GCGCCGCGGTTTGCCGACCTGCCATGTGAAGTTTGGCGAAGCAAACGCGATTCTC
GCTGGCGACGCTTTACAAACGCTGGCGTTCTCGATTCTAAGCGATGCCGATATGC
CGGAAGTGTCGGATCGCGACAGAATTTCGATGATTTCTGAACTGGCGAGCGCCAG
CGGTATTGCCGGAATGTGCGGTGGTCAGGCACTAGATTTAGACGCGGAAGGCAAA
CACGTACCTCTGGACGCGCTTGAGCGTATTCATCGTCATAAAACCGGCGCATTGAT
TCGCGCCGCCGTTCGCCTTGGTGCATTAAGCGCCGGAGATAAAGGGCGTCGTGCT
CTGCCAGTACTCGACAAGTACGCAGAGAGCATCGGCCTTGCCTTCCAGGTTCAAG
ATGACATCCTGGATGTGGTAGGAGATACTGCAACGTTGGGAAAACGCCAGGGTGC
CGACCAGCAACTTGGTAAAAGTACCTACCCTGCACTTCTGGGTCTTGAGCAAGCC
CGGAAGAAAGCCCGGGATCTGATCGACGATGCCCGTCAGTCGCTGAAACAACTGG
CTGAACAGTCACTCGATACCTCGGCACTGGAAGCGCTAGCGGACTACATCATCCA
GCGTAATAAATAAGTCGAC

**246_pJL1-(CAT5aa)-LS_Msp**

TCTAGAAATAATTTTGTTTAACTTTAAGAAGGAGATATACATATGGAGAAAAAAATCC
GTCGTTCGGGGAACTATAACCCGTCACGTTGGGATGTGAACTTCATCCAGTCTTTA
CTGAGCGATTATAAGGAAGACAAACATGTCATTCGTGCATCCGAACTGGTTACGTT
AGTCAAAATGGAATTAGAGAAAGAAACAGACCAGATTCGGCAACTGGAATTGATCG

ACGATTTGCAACGCATGGGCTTATCTGATCACTTCCAAAACGAATTTAAGGAGATT
CTGAGCTCCATCTATCTGGATCATCATTACTATAAAAATCCTTTTCCCAAAGAAGAA
CGCGACCTGTACAGTACCTCTCTGGCATTCCGGCTGCTGCGTGAACATGGCTTCC
AGGTAGCACAAGAGGTTTTCGATAGTTTCAAAAATGAAGAAGGTGAATTTAAAGAA
AGTCTGTCCGATGATACTCGGGGGCTCCTGCAACTTTATGAAGCCTCCTTTCTGCT
GACCGAGGGCGAAACCACCCTGGAGTCCGCCCGCGAGTTCGCAACCAAGTTTCT
GGAAGAAAAAGTGAATGAAGGGGGCGTAGATGGCGACCTGCTCACCCGCATTGC
GTATAGCCTGGACATCCCGCTCCACTGGCGCATTAAACGCCCGAACGCTCCGGTT
TGGATTGAGTGGTACCGTAAGCGCCCTGATATGAACCCGGTTGTTTTAGAGCTGG
CCATCTTAGACCTGAACATCGTCCAGGCCCAGTTTCAAGAGGAACTGAAAGAAAGC
TTTCGCTGGTGGCGTAACACCGGTTTTGTGGAAAAGTTACCATTTGCACGCGACCG
TCTTGTCGAGTGCTACTTTTGGAATACAGGCATTATCGAACCGCGTCAGCACGCCA
GTGCCCGTATCATGATGGGAAAAGTTAATGCGCTGATCACCGTGATCGACGACAT
CTATGATGTGTACGGCACCTTAGAAGAGCTCGAACAATTCACCGACCTTATCCGGC
GCTGGGATATTAATTCGATCGATCAGCTTCCGGATTATATGCAGCTGTGTTTTCTG
GCGCTTAATAATTTTGTTGATGACACGAGTTACGACGTTATGAAAGAAAAAGGTGT
GAACGTGATCCCGTACTTACGCCAATCATGGGTGGACCTGGCGGACAAGTATATG
GTTGAAGCCCGGTGGTTTTATGGTGGACATAAACCTTCGTTGGAAGAATATTTAGA
AAATTCTTGGCAGTCTATCTCCGGCCCCTGTATGTTGACCCATATTTTCTTCCGTGT
CACCGACAGCTTTACGAAAGAAACGGTAGACTCTCTCTATAAGTACCATGACCTGG
TTCGTTGGTCCTCCTTTGTACTGCGCCTTGCAGATGATCTGGGCACAAGTGTAGAA
GAAGTTTCTCGTGGTGATGTCCCTAAAAGCCTCCAGTGTTATATGTCTGACTACAAT
GCATCGGAAGCGGAGGCGCGCAAGCATGTAAAATGGCTTATCGCTGAGGTGTGG
AAGAAAATGAACGCAGAGCGCGTAAGCAAGGATTCGCCATTCGGTAAGGATTTCAT
TGGATGTGCGGTGGACTTAGGGCGGATGGCTCAGTTAATGTACCACAACGGCGAC
GGTCATGGCACGCAGCACCCAATTATTCATCAGCAGATGACGCGGACGCTGTTCG
AGCCGTTCGCGTAAGGATCCTGACTCGAGGTCGAC

**247_pJL1-(CAT5aa)-LS_Cli**

TCTAGAAATAATTTTGTTTAACTTTAAGAAGGAGATATACATATGGAGAAAAAAATCC
GTCGGAGCGCTAACTATCAACCGTCAATTTGGGACCATGATTTTCTGCAATCGCTG
AACTCAAACTATACAGACGAGGCGTACAAACGGCGCGCCGAGGAACTTCGCGGTA
AGGTCAAAATTGCCATTAAAGACGTGATTGAACCGCTGGACCAGCTCGAACTGATT
GACAACCTGCAACGTCTCGGTCTGGCGCATCGCTTCGAAACGGAAATTCGTAATAT
CTTAAATAACATCTATAATAACAACAAAGATTATAACTGGCGCAAAGAAAATCTGTA
TGCAACTAGCCTCGAATTTCGGCTGCTGCGTCAGCATGGCTACCCGGTGTCGCAG
GAGGTCTTCAATGGTTTCAAAGATGATCAGGGTGGTTTCATTTGTGATGACTTTAAA
GGAATCCTTTCGCTCCACGAGGCCAGCTATTATTCACTGGAAGGTGAATCAATTAT
GGAGGAAGCATGGCAGTTTACGTCAAAACACCTGAAGGAAGTGATGATCTCTAAAA
ACATGGAAGAAGATGTGTTCGTCGCGGAACAAGCTAAACGTGCTCTGGAACTGCC
GCTGCATTGGAAAGTCCCGATGCTGGAAGCACGCTGGTTTATCCATATTTACGAAC
GCCGCGAAGATAAAAATCACCTGCTGCTGGAATTAGCGAAAATGGAATTTAACACC
CTGCAGGCCATTTACCAGGAAGAATTAAAGGAAATCTCGGGTTGGTGGAAGGATA

CGGGGCTTGGAGAGAAACTGTCCTTTGCTCGCAACCGGTTAGTGGCGTCGTTCTT
GTGGTCTATGGGGATCGCGTTCGAACCACAATTCGCCTATTGTCGCCGTGTGTTAA
CTATCTCTATTGCACTGATTACGGTGATCGATGACATTTACGACGTATACGGTACCC
TGGACGAGTTAGAAATCTTTACGGACGCCGTCGAACGGTGGGATATCAATTATGC
GCTCAAACACTTACCTGGTTATATGAAAATGTGCTTTCTGGCGCTGTATAATTTTGT
AAACGAATTTGCATATTATGTGCTGAAGCAACAGGACTTTGATCTGCTGTTGTCTAT
CAAAAACGCCTGGCTGGGATTGATTCAAGCTTACCTTGTTGAAGCGAAGTGGTACC
ACAGCAAATACACCCCGAAACTGGAAGAGTATCTGGAGAACGGCCTGGTTAGCAT
CACCGGGCCCCTGATTATTACCATCTCTTATCTGTCAGGGACCAACCCAATTATCA
AAAAAGAACTGGAATTTCTTGAAAGTAACCCGGATATTGTTCACTGGTCTTCCAAAA
TCTTCCGCCTGCAGGATGATCTGGGCACCAGCTCAGACGAAATTCAACGGGGCGA
CGTACCAAAATCAATTCAGTGCTATATGCATGAAACCGGTGCGAGTGAGGAGGTA
GCCCGTCAGCACATCAAGGATATGATGCGCCAAATGTGGAAAAAAGTCAATGCTTA
TACCGCAGACAAGGACAGTCCGCTGACCGGCACTACAACGGAATTTCTGCTGAAC
TTAGTCCGTATGAGCCATTTTATGTATTTGCATGGCGACGGACACGGCGTTCAGAA
CCAGGAAACCATTGATGTCGGCTTCACCTTGCTGTTTCAGCCAATCCCTTTGGAAG
ATAAGCACATGGCGTTTACCGCAAGCCCGGGCACCAAAGGCTAAGGATCCTGACT
CGAGGTCGAC

**248_pJL1-(CAT5aa)-LS_Pfr**

TCTAGAAATAATTTTGTTTAACTTTAAGAAGGAGATATACATATGGAGAAAAAAATCC
GTCGTAGCGGTAATTATAGCCCGAGCTTTTGGAATGCAGATTATATTCTGAGCCTG
AACAACCATTATAAAGAAGAAAGCCGTCATATGAAACGTGCCGGTGAACTGATTGT
TCAGGTTAAAATGGTTATGGGCAAAGAAACCGATCCGGTTGTGCAGCTGGAACTG
ATCGATGATCTGCATAAACTGGCACTGAGCCATCATTTTGAGAAAGAAATTAAAGA
GATCCTGTTCAACATCAGCATCTACGACCATAAAATCATGGTTGAACGTGATCTGTA
TAGCACCGCACTGGCATTCCGGTTGCTCCGTCAGTATGGCTTCAAGGTGCCGCAG
GAGGTTTTCGATTGTTTTAAAAACGACAATGGAGAATTTAAACGGTCTTTAAGCTCC
GATACGAAAGGTCTGCTGCAGTTGTATGAAGCCTCCTTTCTGCTGACGGAAGGTG
AAATGACACTGGAACTGGCGCGTGAGTTCGCGACCATCTTTCTTCAGGAGAAACT
GAACGATAAAACAATCGACGACGACGATGATGCGGATACAAACCTCATTTCTTGCG
TGCGTCATTCACTCGATATCCCGATCCATTGGCGCATCCAACGTCCGAACGCCAGT
TGGTGGATCGATGCCTACAAACGCCGCAGTCACATGAATCCTCTGCTTGAGTTAGC
AAAGTTAGACCTGAATATTTTTCAGGCACAGTTTCAACAGGAACTGAAACAGGACC
TGGGTTGGTGGAAAAATACATGTCTCGCAGAGAAGCTGCCGTTTACCCGTGACCG
CCTGGTGGAATGCTACTTCTGGTGCACCGGTATCATTCAGCCGCTGCAGCATGAG
AACGCTCGCGTTACTCTGGCAAAAGTTAACGCCTTGATCACCACGCTGGACGACAT
TTACGATGTATACGGCACCCTGGAGGAACTGGAACTGTTCACGGAAGCGATTCGG
CGTTGGGACGTTAGTAGCATTGACCACTTACCGAACTATATGCAGCTGTGCTTCCT
CGCCCTGAACAATTTTGTCGATGACACCGCGTACGATGTTATGAAAGAAAAGGACA
TTAACATTATCCCGTATCTGCGGAAATCGTGGTTGGACCTCGCCGAGACATACCTG
GTGGAAGCTAAATGGTTCTATTCAGGCCATAAACCGAATATGGAAGAATATTTAAAT
AATGCGTGGATCTCGATTAGCGGTCCGGTTATGTTGTGCCATGTATTCTTCCGCGT

GACTGATTCCATTACCCGCGAAACCGTAGAATCTCTGTTCAAATATCACGACCTTAT
TCGTTACTCCTCTACCATCCTTCGCCTGGCGGATGACTTAGGCACAAGCCTGGAAG
AGGTTAGCCGTGGTGATGTTCCGAAAAGCATTCAGTGTTATATGAATGATAACAAC
GCCAGCGAAGAAGAAGCACGTCGTCACGTTCGTTGGCTGATTGCAGAAACCTGGA
AAAAAATCAACGAAGAAGTTTGGAGCGCAGATAGCCCGTTTTGCAAAGATTTTATT
GCATGTGCAGCAGATATGGGTCGTATGGCGCAGTTTATGTATCATAATGGTGATGG
TCATGGCATTCAGAATCCGCAGATTCATCAGCAGATGACCGATATTCTGTTTGAAC
AGTGGCTGTAAGGATGTCGAC

**350_pJL1-(CAT7aa)-LS_Ste**

TCTAGAAATAATTTTGTTTAACTTTAAGAAGGAGATATACATATGCATATGGAGAAAA
AAATCCGTCGTAGCGGTAACTATAAACCGAGCCGTTGGGATGTTGATTTTATGCAG
AGCCTGAATAGCGATTATCAAGAAGAACGTCATCGTACCAAAGCAAGCGAACTGAT
TACCCAGGTTAAAAACCTGCTGGAAAAAGAAACCAGTGATGATCCGATTCGTCAGC
TGGAACTGATTGATGATCTGCAGCGTCTGGGTCTGAGCGATCATTTTGAACATGAA
TTTAAAGAGGTGCTGAACAGCATCTACCTGGACAACAAATATTACAACATCAACATC
ATGAAAGAAACGACCAGCAGCCGTGATCTGTATAGCACCGCACTGGCATTTCGTCT
GCTGCGTGAACATGGTTTTCAGGTTGCACAAGAAGTTTTCGACTGCTTCAAAAATG
AAGAGGGTGAGTTTAAAGCAAGCCTGTCAGATGATCCGCGTGGTCTGCTGCAGCT
GTATGAAGCAAGCTTTCTGTTTAAAGAAGGCGAAAACACCCTGGAAATTGCCCGTG
AATTTGCAACCAAACTGCTGCAAGAAAAAGTGAACAGCTCCGATGAAATTGATGAT
AATCTGCTGAGCAGCATTCGTTATAGTCTGGAAATTCCGACCTATTGGAGCGTTATT
CGTCCGAATGTTAGCGTTTGGATTGATGCATATCGTAAACGTCCGGATATGAATCC
GGTTGTTCTGGAACTGGCAATTCTGGATGCCAATATTATGCAGGCACAGCTGCAAC
AAGAACTGAAAGAAGCATTAGGTTGGTGGCGTAATACCTGGTTTGTTGAAAAACTG
CCGTTTGCACGTGATCGTCTGGTTGAAAGCTATTTTTGGAGCACCGGTATGGTTCC
GCGTCGTCAGCATAAAACCGCACGTCAGCTGATGGCAAAAGTTATTGCCCTGATTA
CCGTGATGGATGATATCTATGACGTTTATGGTACACTGGAAGAACTGGAACTGTTT
ACCGATGCCTTTCGTCGCTGGGATGTTAGCAGCATTGATCATCTGCCGACCTATAT
GCAACTGTGTTTTCTGAGCATTAACAACTTTGTTGTTGACACCGCCTACAACATTCT
TAAAGAAACCGGTGTTAATGTGACCACCTATCTGGAAAAAAGCTGGGTTGATCAGG
CAGAAAATTATCTGATGGAAAGCAAATGGTTCTACAGCGGTCATAAACCGTCACTG
GATGAATACCTGGAAAATAGTTGGATTAGCGTTAGCGGTCCGTGTGTTCTGACCCA
TGAATTTTTTGGTGTTACCGATAGCCTGGCAAAAGATACCCTGGATAGCCTGTATG
AATATCACGATATTGTTCGTTGGAGCAGCTATCTGCTGCGCCTGGCCGATGATCTG
GGCACCAGCGTTGAAGAGGTTAGCCGTGGTGATGTTCCGAAAAGCATTCAGTGTT
ATATGAACGATAATAACGCCAGCGAAGAAGAAGCACGCGAACACGTTAAAGGTCT
GATTCGTGTTATGTGGAAAAAAATGAATGCCGAACGTGTTAGCGAAGATAGCCCGT
TTTGTAAAGATTTTATTCGTTGTTGTGAGGACCTGGGTCGTATGGCACAGTTTATGT
ATCATTATGGTGATGGTCATGGCACCCAGCATGCCAAAATTCATCAGCAGATCACC
GATTGTCTGTTTCAGCCGTTTGCCTAAGTCGACGTCGAC

**351_pJL1-(CAT7aa)-LS_Lan**

TCTAGAAATAATTTTGTTTAACTTTAAGAAGGAGATATACATATGCATATGGAGAAAA
AAATCCGTCGTAGCGGTAACTATAATCCGACCGCATGGGATTTTAACTATATTCAG
AGCCTGGACAACCAGTACAAAAAGAACGTTATAGCACCCGTCATGCAGAACTGAC
CGTTCAGGTTAAAAAACTGCTGGAAGAAGAAATGGAAGCCGTTCAGAAACTGGAAC
TGATTGAGGATCTGAAAAATCTGGGTATCAGCTATCCGTTCAAAGATAACATTCAG
CAGATCCTGAACCAGATTTACAACGAACATAAATGCTGCCATAACAGCGAAGTGGA
AGAAAAAGACCTGTATTTTACCGCACTGCGTTTTCGTCTGCTGCGTCAGCAGGGTT
TTGAAGTTAGCCAAGAAGTTTTCGACCACTTCAAAAATGAAAAAGGCACCGATTTTA
AACCGAACCTGGCAGATGATACCAAAGGTCTGCTGCAGCTGTATGAAGCAAGCTTT
CTGCTGCGCGAAGCCGAAGATACCCTGGAACTGGCACGTCAGTTTAGCACCAAAC
TGCTGCAGAAAAAAGTGGATGAAAATGGCGACGATAAAATCGAAGATAATCTGCTG
CTGTGGATTCGTCGTAGTCTGGAACTGCCGCTGCATTGGCGTGTTCAGCGTCTGG
AAGCACGTGGTTTTCTGGATGCCTATGTTCGTCGTCCGGATATGAATCCGATTGTT
TTTGAACTGGCAAAGCTGGATTTCAATATTACCCAGGCAACCCAGCAAGAAGAACT
GAAAGACCTGAGCCGTTGGTGGAATAGCACCGGTCTGGCAGAAAAACTGCCGTTT
GCACGTGATCGTGTTGTTGAAAGCTATTTTTGGGCAATGGGCACCTTTGAACCGCA
TCAGTATGGTTATCAGCGTGAACTGGTTGCAAAAATCATTGCACTGGCAACCGTTG
TTGATGATGTCTATGATGTTTATGGCACCCTGGAAGAATTAGAACTGTTTACCGATG
CAATTCGTCGTTGGGATCGTGAAAGCATTGATCAGCTGCCGTATTATATGCAGCTG
TGTTTTCTGACCGTGAACAACTTTGTGTTTGAGCTGGCACATGATGTGCTGAAAGA
TAAAAGCTTTAATTGTCTGCCGCATCTGCAGCGTAGCTGGCTGGATCTGGCCGAAG
CATATCTGGTTGAAGCAAAATGGTATCATAGCCGTTATACCCCGAGCCTGGAAGAG
TATCTGAATATTGCACGTGTTAGCGTTACCTGTCCGACCATTGTTAGCCAGATGTAT
TTTGCACTGCCGATTCCGATTGAAAAACCGGTGATTGAAATCATGTACAAATACCA
CGATATCCTGTATCTGAGCGGTATGCTGCTGCGCCTGCCGGATGATCTGGGCACC
GCATCATTTGAACTGAAACGTGGTGATGTTCAGAAAGCGGTTCAGTGCTATATGAA
AGAACGTAATGTTCCGGAAAATGAAGCCCGTGAACATGTGAAATTTCTGATTCGTG
AAGCCAGCAAGCAGATCAATACCGCAATGGCCACCGATTGTCCGTTTACCGAAGA
TTTTGCAGTTGCAGCAGCCAATCTGGGTCGTGTTGCAAATTTTGTTTATGTGGATG
GTGATGGTTTTGGTGTTCAGCACAGCAAAATCTATGAGCAGATTGGTACACTGATG
TTTGAACCGTATCCGTAAGTCGACGTCGAC

**352_pJL1-(CAT7aa)-LS_Sly**

TCTAGAAATAATTTTGTTTAACTTTAAGAAGGAGATATACATATGCATATGGAGAAAA
AAATCCGTCGTAGCGGTAATTATGAACCGACCATGTGGAACTATGAATATATCCAG
AGCACCCATAATCATCATGTGGGCGAGAAATATATGAAACGCTTTAATGAACTGAA
AGCCGAAATGAAAAAGCATCTGATGATGATGCTGCACGAAGAAAGCCAAGAACTG
GAAAAACTGGAACTGATTGATAATCTGCAGCGTCTGGGTGTTAGCTATCACTTTAA
AGATGAAATCATTCAGATCCTGCGCAGCATTCATGATCAGTCAAGCAGCGAAGCAA
CCAGCGCAAATAGCCTGTATTATACCGCACTGAAATTTCGTATTCTGCGTCAGCAT

GGCTTTTATATCAGCCAGGATATTCTGAACGACTTCAAAGATGAGCAGGGCCATTT
TAAACAGAGCCTGTGTAAAGATACCAAAGGTCTGCTGCAGCTGTATGAAGCAAGCT
TTCTGAGCACCAAAAGCGAAACCAGCACACTGCTGGAAAGCGCCAATACCTTTGC
AATGAGCCATCTGAAAAACTATCTGAATGGTGGTGATGAAGAGAACAACTGGATGG
TTAAACTGGTTCGTCATGCACTGGAAGTTCCGCTGCATTGTATGATGCTTCGTGTT
GAAACCCGTTGGTATATCGACATCTATGAAAATATTCCGAATGCCAATCCGCTGCT
GATTGAACTGGCCAAACTGGATTTTAACTTTGTTCAGGCAATGCACCAGCAAGAAC
TGCGTAATCTGAGCCGTTGGTGGAAAAAAAGCATGCTGGCAGAAAAACTGCCGTTT
GCACGTGATCGTATTGTTGAAGCATTTCAGTGGATTACCGGCATGATTTTTGAGAG
CCAAGAAAATGAATTTTGCCGCATCATGCTGACCAAAGTTACCGCAATGGCAACCG
TTATTGATGATATCTATGATGTTTATGGCACCCTGGATGAGCTGGAAATCTTTACCC
ATGCAATTCAGCGCATGGAAATTAAAGCAATGGATGAACTGCCGCACTACATGAAA
CTGTGTTATCTGGCACTGTTTAATACCAGCAGCGAAATTGCATATCAGGTGCTGAA
AGAACAGGGCATTAACATTATGCCGTATCTGACCAAAAGCTGGGCTGATCTGAGCA
AAAGTTATCTGCAAGAAGCACGTTGGTATTATAGCGGTTATACCCCGAGCCTGGAT
GAATACATGGAAAATGCATGGATTAGCGTTGGTAGCCTGGTTATGGTTGTTAATGC
ATTTTTTCTGGTGACCAATCCGATCACCAAAGAAGTTCTGGAATACCTGTTCAGCAA
CAAATATCCGGACATTATTCGTTGGCCTGCAACCATTATTCGCCTGACCGATGATC
TGGCGACCAGCAGCAATGAAATGAAACGTGGTGATGTTCCGAAAAGCATTCAGTG
CTATATGAAAGAAAATGGTGCCAGCGAAGAAGAAGCCCGTAAACATATTAACCTGA
TGATCAAAGAAACCTGGAAAATGATCAATACCGCACAGCATGATAACAGCCTGTTT
TGCGAAAAATTCATGGGTTGTGCAGTTAATATTGCACGTACCGGTCAGACCATTTA
TCAGCATGGTGATGGTCATGGTATCCAGAATTACAAAATTCAGAACCGCATCAGCA
AACTGTTTTTTGAACCGATTACCATCAGCATGCCGTAAGTCGACGTCGAC

**353_pJL1-(CAT7aa)-LS-Pab**

TCTAGAAATAATTTTGTTTAACTTTAAGAAGGAGATATACATATGCATATGGAGAAAA
AAATCCGTCGTCGTGGTAATTATCATAGCAATCTGTGGGATGATGATTTCATTCAGA
GCCTGAGCACCCCGTATGGTGAACCGAGCTATCGTGAACGTGCAGAACGTCTGAA
AGGTGAGATCAAAAAAATGTTTCGCAGCATGAGCAAAGATGATGGCGAACTGATTA
CACCGCTGAATGATCTGATTCAGCGTCTGTGGATGGTTGATAGCGTGCAGCGTCT
GGGTATTGATCGTCATTTCAAAAACGAAATCAAAAGCGCACTGGACTATGTGTATA
GCTATTGGAATGAAAAAGGTATTGGTTGCGGTCGTGATAGCGTTGTTGCCGATCTG
AATAGCACCGCACTGGGTTTTCGTACCCTGCGTCTGCATGGTTATAATGTTAGCAG
CGAAGTTCTGAAAGTGTTCGAAGATCAGAATGGTCAGTTTGCATGTAGCCCGAGCA
AAACCGAAGGTGAAATTCGTAGCGCACTGAATCTGTATCGTGCAAGCCTGATTGCA
TTTCCGGGTGAAAAAGTTATGGATGATGCAGAAATCTTTAGCAGCCGCTATCTGAA
AGAAGCCGTTCAAGAAATTCCGGATTGTAGCCTGAGCCAAGAAATTGCCTATGCAC
TGGAATATGGTTGGCATACCAATATGCCTCGTCTGGAAGCACGCAATTATATGGAT
GTTTTTGGTCATCCGAGCAGCCCGTGGCTGAAAAAAAACAAAACACAGTATATGGA
CGGCGAGAAACTGCTGGAACTGGCAAAACTGGAATTTAACATTTTTCACAGCCTGC
AGCAAGAGGAACTGCAGTATATTAGCCGTTGGTGGAAAGATAGTGGTCTGCCGAA

ACTGGCATTTAGCCGTCATCGTCATGTTGAGTATTATACCCTGGGTAGCTGTATTG
CAACCGATCCGAAACATCGTGCATTTCGTCTGGGTTTTGTTAAAACCTGTCATCTGA
ATACCGTGCTGGATGATATCTATGATACCTTTGGCACCATGGATGAAATCGAACTG
TTTACCGAAGCAGTTCGTCGTTGGGACCCGAGTGAAACCGAAAGCCTGCCGGATT
ATATGAAAGGTGTTTATATGGTTCTGTATGAAGCCCTGACCGAAATGGCACAAGAA
GCACAGAAAACACAGGGTCGCGATACCCTGAATTATGCACGTAAAGCATGGGAAA
TTTATCTGGACAGCTATATCCAAGAGGCAAAATGGATTGCAAGCGGTTATCTGCCG
ACCTTTCAAGAATATTTTGAGAACGGTAAAATCAGCAGCGCATATCGTGCAGCAGC
ACTGACCCCGATTCTGACCCTGGATGTTCCGCTGCCGGAATACATTCTGAAAGGC
ATTGATTTTCCGAGCCGCTTTAATGATCTGGCAAGCAGCTTTCTGCGCCTGCGTGG
TGATACCCGTTGTTATAAAGCAGATCGTGCACGTGGTGAAGAAGCAAGCTGTATTA
GCTGTTACATGAAAGATAATCCGGGTAGCACCGAAGAAGATGCCCTGAATCATATT
AACAGCATGATCAACGAGATCATCAAAGAACTGAATTGGGAACTGCTGCGTCCGG
ATAGCAATATTCCGATGCCAGCGCGTAAACATGCATTTGATATTACCCGTGCACTG
CATCACCTGTATAAATACCGTGATGGTTTTAGCGTTGCAACCAAAGAAACCAAAAG
TCTGGTTAGCCGCATGGTTCTGGAACCGGTGCCGCTGTAAGTCGACGTCGAC

**320_pJL1-(CAT5aa)-BS_Agr**

TCTAGAAATAATTTTGTTTAACTTTAAGAAGGAGATATACATATGGAGAAAAAAATCA
GCGCGGGTGTTTCTGCGGTTTCTAAAGTTTCTTCTCTGGTTTGCGACCTGTCTTCT
ACCTCTGGTCTGATCCGTCGTACCGCGAACCCGCACCCGAACGTTTGGGGTTACG
ACCTGGTTCACTCTCTGAAATCTCCGTACATCGACTCTTCTTACCGTGAACGTGCG
GAAGTTCTGGTTTCTGAAATCAAAGCGATGCTGAACCCGGCGATCACCGGTGACG
GTGAATCTATGATCACCCCGTCTGCGTACGACACCGCGTGGGTTGCGCGTGTTCC
GGCGATCGACGGTTCTGCGCGTCCGCAGTTCCCGCAGACCGTTGACTGGATTCTG
AAAAACCAGCTGAAAGACGGTTCTTGGGGTATCCAGTCTCACTTCCTGCTGTCTGA
CCGTCTGCTGGCGACCCTGTCTTGCGTTCTGGTTCTGCTGAAATGGAACGTTGGT
GACCTGCAGGTTGAACAGGGTATCGAGTTCATCAAATCTAACCTGGAACTGGTTAA
AGACGAAACCGACCAGGACTCTCTGGTTACCGACTTCGAAATCATCTTCCCGTCTC
TGCTGCGTGAAGCGCAGTCTCTGCGTCTGGGTCTGCCGTACGACCTGCCGTACAT
CCACCTGCTGCAGACCAAACGTCAGGAACGTCTGGCGAAACTGTCTCGTGAAGAA
ATCTACGCGGTTCCGTCTCCGCTGCTGTACTCTCTGGAAGGTATCCAGGACATCGT
TGAATGGGAACGTATCATGGAAGTTCAGTCTCAGGACGGTTCTTTCCTGTCTTCTC
CGGCGTCTACCGCGTGCGTTTTCATGCACACCGGTGACGCGAAATGCCTGGAGTT
CCTGAACTCTGTTATGATCAAATTCGGTAACTTCGTTCCGTGCCTGTACCCGGTTG
ACCTGCTGGAACGTCTGCTGATCGTTGACAACATCGTTCGTCTGGGTATCTACCGT
CACTTCGAAAAAGAAATCAAAGAAGCGCTGGACTACGTTTACCGTCACTGGAACGA
ACGTGGTATCGGTTGGGGTCGTCTGAACCCGATCGCGGACCTGGAAACCACCGC
GCTGGGTTTCCGTCTGCTGCGTCTGCACCGTTACAACGTTTCTCCGGCGATCTTCG
ACAACTTCAAAGACGCGAACGGTAAATTCATCTGCTCTACCGGTCAGTTCAACAAA
GACGTTGCGTCTATGCTGAACCTGTACCGTGCGTCTCAGCTGGCGTTTCCGGGTG
AAAACATCCTGGACGAAGCGAAATCTTTCGCGACCAAATACCTGCGTGAAGCGCT

GGAAAAATCTGAAACCTCTTCTGCGTGGAACAACAAACAGAACCTGTCTCAGGAAA
TCAAATACGCGCTGAAAACCTCTTGGCACGCGTCTGTTCCGCGTGTTGAAGCGAA
ACGTTACTGCCAGGTTTACCGTCCGGACTACGCGCGTATCGCGAAATGCGTTTACA
AACTGCCGTACGTGAACAACGAAAAATTCCTGGAACTGGGTAAACTGGACTTCAAC
ATCATCCAGTCTATCCACCAGGAAGAAATGAAAAACGTTACCTCTTGGTTCCGTGA
CTCTGGTCTGCCGCTGTTCACCTTCGCGCGTGAACGTCCGCTGGAGTTCTACTTC
CTGGTTGCGGCGGGTACCTACGAACCGCAGTACGCGAAATGCCGTTTCCTGTTCA
CCAAAGTTGCGTGCCTGCAGACCGTTCTGGACGACATGTACGACACCTACGGTAC
CCTGGACGAACTGAAACTGTTCACCGAAGCGGTTCGTCGTTGGGACCTGTCTTTCA
CCGAAAACCTGCCGGACTACATGAAACTGTGCTACCAGATTTACTACGACATCGTT
CACGAAGTTGCGTGGGAAGCGGAAAAAGAACAGGGTCGTGAACTGGTTTCTTTCT
TCCGTAAAGGTTGGGAAGACTACCTGCTGGGTTACTACGAAGAAGCGGAATGGCT
GGCGGCGGAATACGTTCCGACCCTGGACGAATACATCAAAAACGGTATCACCTCT
ATCGGTCAGCGTATCCTGCTGCTGTCTGGTGTTCTGATCATGGACGGTCAGCTGCT
GTCTCAGGAAGCGCTGGAAAAAGTTGACTATCCGGGTCGTCGTGTTCTGACCGAA
CTGAACTCTCTGATCTCTCGTCTGGCGGACGACACCAAAACCTACAAAGCGGAAAA
AGCGCGTGGTGAACTGGCGTCTTCTATCGAATGCTACATGAAAGACCACCCGGAA
TGCACCGAAGAAGAAGCGCTGGACCACATCTACTCTATCCTGGAACCGGCGGTTA
AAGAACTGACCCGTGAGTTCCTGAAACCGGACGACGTTCCGTTCGCGTGCAAAAA
AATGCTGTTCGAAGAAACCCGTGTTACTATGGTTATCTTCAAAGACGGTGACGGTT
TCGGTGTTTCTAAACTGGAAGTTAAAGACCACATCAAAGAATGCCTGATCGAACCG
CTGCCGCTGTAAGTCGAC

**249_11a_pJL1_CATrbs(5aa)_PS_Agr**

TCTAGAAATAATTTTGTTTAACTTTAAGAAGGAGATATACATATGGAGAAAAAAATCC
GTCGTGGCAAATCCATTACCCCGAGTATCTCGATGAGCTCTACGACGGTTGTGACA
GACGACGGCGTCCGCCGTCGTATGGGTGATTTTCATTCCAATTTATGGGACGACG
ATGTGATTCAGAGCTTACCAACAGCGTACGAAGAAAAATCCTATCTTGAACGTGCG
GAAAAACTGATTGGAGAAGTAAAAAACATGTTTAATTCCATGAGCCTTGAGGATGG
TGAACTTATGAGTCCACTGAATGATCTGATCCAGCGCCTGTGGATCGTAGACAGCC
TGGAGCGGTTAGGCATCCACCGTCACTTCAAAGATGAGATTAAATCGGCTTTGGAT
TATGTGTATTCATACTGGGGAGAGAATGGTATTGGGTGTGGCGGGAAAGTGTGG
TAACAGACTTGAACAGCACTGCACTGGGCCTGCGGACCCTGCGCCTCCACGGCTA
TCCAGTCAGCTCTGATGTGTTTAAGGCTTTTAAAGGACAAAACGGACAGTTTTCTTG
TTCCGAGAATATTCAAACCGATGAAGAAATCCGTGGCGTGCTGAATTTGTTCCGTG
CAAGCCTGATTGCCTTTCCGGGTGAGAAAATCATGGACGAGGCCGAAATCTTTTCA
ACCAAATACCTGAAGGAAGCTCTGCAAAAGATCCCTGTATCCTCCTTATCCCGGGA
AATCGGCGATGTGTTGGAATACGGCTGGCATACTTATTTGCCGCGTTTAGAGGCAC
GCAATTATATTCAGGTGTTTGGTCAGGACACCGAAAATACCAAAAGCTACGTTAAA
AGCAAAAAACTTTTGGAACTGGCTAAGCTGGAATTTAACATTTTCCAGAGCTTGCA
GAAACGTGAACTGGAATCGTTGGTTCGCTGGTGGAAAGAAAGCGGCTTTCCGGAG
ATGACCTTTTGTCGTCATCGTCATGTGGAATATTATACCCTGGCCTCCTGCATTGC

GTTCGAACCGCAGCATAGTGGTTTCCGCCTCGGATTTGCGAAAACTTGTCATCTGA
TTACCGTACTGGATGATATGTATGATACCTTCGGCACGGTTGATGAATTAGAACTGT
TCACCGCCACTATGAAACGCTGGGACCCGTCTTCGATCGATTGCTTACCGGAATAC
ATGAAAGGCGTATATATCGCCGTATACGACACCGTGAATGAGATGGCGCGCGAAG
CGGAGGAGGCCCAAGGGCGCGATACCTTGACCTATGCACGTGAGGCGTGGGAAG
CCTATATCGATAGTTACATGCAAGAAGCCCGTTGGATCGCCACCGGCTATTTACCT
TCATTTGATGAGTATTATGAAAATGGCAAAGTCAGTTGCGGACATCGTATCTCGGC
CCTGCAGCCTATCTTAACCATGGACATTCCGTTTCCCGACCACATCCTGAAAGAAG
TTGATTTCCCGAGCAAACTCAATGATCTCGCGTGTGCCATTCTCCGCCTGCGCGGT
GATACGCGCTGCTATAAGGCGGATCGTGCACGTGGCGAAGAAGCGAGCAGCATC
TCCTGTTATATGAAAGATAACCCTGGCGTGAGCGAGGAAGATGCTTTAGATCACAT
TAATGCCATGATCTCAGATGTAATTAAAGGGCTTAACTGGGAGTTATTAAAACCTGA
TATTAATGTGCCCATCAGTGCGAAAAAACACGCGTTCGACATTGCGCGCGCTTTCC
ACTATGGTTATAAATATCGCGATGGTTATAGCGTGGCGAATGTAGAGACGAAATCG
CTGGTGACCCGGACCCTGCTGGAATCTGTGCCCTTATGAGGATCCTGACTCGAGG
TCGAC

**250_11b_pJL1_CATrbs(5aa)_PS_Pab**

TCTAGAAATAATTTTGTTTAACTTTAAGAAGGAGATATACATATGGAGAAAAAAATCC
GCCGTATGGGTGATTTTCACTCTAATCTGTGGAACGACGATTTTATTCAAAGCCTGA
GCACGTCGTACGGCGAACCTAGTTACCGCGAACGCGCTGAGCGTCTCATTGGCGA
AGTCAAGAAAATGTTTAATAGCATGTCATCTGAAGACGGCGAGCTGATTAGCCCCC
ATAATGACTTAATTCAACGCGTGTGGATGGTGGACAGCGTCGAACGTCTGGGCATT
GAACGCCACTTCAAGAATGAGATTAAAAGCGCTCTCGATTATGTCTATTCCTACTG
GTCAGAAAAAGGCATCGGCTGCGGGCGCGAATCGGTGGTAGCCGACCTGAACAG
TACGGCCTTAGGCTTACGTACACTGCGCCTGCATGGATATGCAGTGTCCGCGGAC
GTTCTCAATTTGTTTAAAGATCAGAATGGTCAATTTGCATGCTCACCATCTCAGACC
GAAGAAGAAATCCGTAGTGTATTAAACCTTTATCGCGCAAGTTTAATTGCCTTCCCG
GGGGAAAAAGTGATGGAAGAGGCGGAAATTTTCTCGGCCAAGTATCTGGAAGAGG
CCCTGCAGAAAATCTCCGTCAGCTCACTTAGCCAAGAAATCCGTGATGTGCTGGAA
TATGGTTGGCATACCTACCTGCCCCGGATGGAGGCGCGCAATCATATTGACGTGT
TTGGCCAGGACACACAGAACTCTAAAAGCTGCATTAATACCGATAAATTATTAGAG
CTTGCGAAATTGGAGTTTAATATCTTTCACTCGTTGCAGAAACGCGAGCTGGAATA
TCTGGTGCGGTGGTGGAAAGACAGCGGCTCCCCGCAGATGACCTTTGGCCGCCA
TCGGCATATTGAATACTATACCCTGGCTAGCTGTATTGCATTTGAACCACAGCACTC
TGGCTTTCGTCTCGGCTTTGCGAAAACTTGCCATATCATCACCATCCTGGACGACA
TGTATGATACCTTTGGCACCGTTGATGAGTTAGAACTTTTTACGGCAGCCATGAAG
CGCTGGGACCCGAGCGCAGCAGACTGCCTTCCTGAGTACATGAAAGTAATGTATA
TGATCGTGTACGACACCGTGAACGAAATGTGTCAGGAGGCGGAAAAAGCGCAGG
GTCGTGACACCCTGGATTATGCCCGTCAGGCGTGGGAAGACTACCTGGATTCTTA
TATGCAGGAAGCTAAGTGGATCGCCACCGGTTACTTACCGACCTTCGAAGAATACT
ATGAGAATGGGAAAGTTTCATCGGGGCATCGTGTGGCTGCACTGCAGCCGATCTT

GACCATGGACATTCCGTTTCCTCCGCACATTTTGAAAGAAGTGGATTTTCCCTCAAA
GCTGAGTGATCTGGCCTGTGCTATCCTGCGCCTGCGTGGAGACACTCGTTGTTAC
AAAGCCGACCGTGCCCGTGGCGAAGAAGCTTCATCGATTTCGTGTTACATGAAAG
ATAATCCTGGCGCGACGGAAGAAGATGCCTTGGATCATATTAATGCCATGATCAGC
GATGTAATTCGTGGGTTGAACTGGGAGCTGTTAAAGCCTAATTCGAGCGTGCCGAT
CAGCAGCAAAAAGCACGTTTTCGATATTTCACGCGCTTTTCATTACGGCTACAAATA
CCGCGATGGCTACTCAGTGGCAAATATCGAAACCAAAAGCTTGGTCAAACGTACA
GTTATTGATCCGGTGACTTTATAAGGATCCTGACTCGAGGTCGAC
